# DNA-LLM2Vec: Architecture-Aware Task-Conditioned Adaptation of Decoder DNA Foundation Models

**DOI:** 10.64898/2026.10.07.757259

**Authors:** Yuan Liang, Ke Zhai, Yujian Gan, Siyou Li, Juexi Shao, Grant H. Watson, Massimo Poesio

## Abstract

Decoder DNA foundation models pretrain with dense next-token prediction and scale to long sequences, but their representations are directional. This is a poor fit for genomic tasks that need sequence context both upstream and downstream of a locus. Bidirectional masked-language-model encoders supply this context, but they pretrain more sparsely and scale less easily to long windows. We introduce DNA-LLM2Vec, a staged decoder-to-encoder framework that opens bidirectional context, repairs the resulting mismatch with masked next-token prediction, and applies contrastive views encoding strand and positional invariance. We instantiate it on two architecturally distinct backbones, DNAGPT and HyenaDNA, and evaluate seven biological task families (grouped from GB, NT, GUE, and GUE+ EPI) with frozen linear probing. The benefits of conversion are architecture-aware and task-conditioned rather than uniform. Crop-based positional contrastive learning helps most for the extended-context families (enhancer–promoter interaction and Virus-COVID), raising frozen Avg-EPI MCC to 27.5% for DNAGPT and 26.7% for HyenaDNA, whereas for shorter, local families the results depend on backbone and tokenisation. Beyond classification, bidirectional context and MNTP improve causal-eQTL variant-effect prediction specifically at distance, and a base-pair mutagenesis diagnostic shows that adaptation organises motif-aware contact structure while reverse-complement training imprints strand-consistent geometry, yet neither predicts the best downstream view. DNA-LLM2Vec therefore produces frozen bidirectional representations whose most effective adaptation view depends on the task structure and the backbone. Code and checkpoints are available at `semi-nothing/dna-llm2vec`.

**Key Points:**

- DNA-LLM2Vec converts pretrained causal DNA foundation models into frozen, bidirectional encoders via bidirectional patching, masked next-token prediction (MNTP) repair, and contrastive adaptation.
- Frozen linear probing (our primary protocol, reusable representation quality) and full fine-tuning (a supplementary analysis, downstream adaptation potential) are not interchangeable. Full fine-tuning reshapes rather than preserves the frozen task-family response, in an architecture-dependent way.
- A base-pair mutagenesis motif-contact analysis provides an orthogonal, non-classification view: adaptation strengthens motif-aware regulatory structure, but this peaks at different adaptation stages across the two architectures and does not, by itself, predict the best downstream variant.
- Ablations and controls support the design: the pretrained backbone supports the broad (Fam-mean) gains, not the objectives alone; the staged pipeline slightly outperforms its shortcuts; and results are stable under alternative pooling.

## Introduction

Modelling DNA places representational demands distinct from those of natural language and proteins. First, cellular phenotypes depend on base pairs both *upstream and downstream* of a locus, so many genomic tasks require access to both flanks of a position at once. Second, the genome is double-stranded: a sequence and its reverse complement carry the same information, and encoding this strand symmetry has been reported to improve genomic models (Mallet and Vert, 2021; Zhou et al., 2022; Schiff et al., 2024). Third, regulation is often *long-range*. Enhancers and other regulatory elements can act on genes tens to hundreds of kilobases away, and occasionally up to a megabase. Many phenotype-relevant variants act at these distances (Furlong and Levine, 2018; Schoenfelder and Fraser, 2019; Maurano et al., 2012). Bidirectional context, strand symmetry, and long-range modelling are therefore three inductive biases generally considered desirable for a DNA representation model.

Reflecting the first of these motivations, most current DNA foundation models adopt bidirectional encoder-style objectives. A recent survey of 15 representative DNA foundation models classifies 9 as masked-language-modelling (MLM) encoders and 6 as causal-language-modelling (CLM) decoders (Shu et al., 2026). Encoder-style models—DNABERT (Ji et al., 2021), DNABERT-2 (Zhou et al., 2024), the Nucleotide Transformer (Dalla-Torre et al., 2025) and Caduceus (Schiff et al., 2024)—produce bidirectional embeddings that match the bidirectional-context demands of many genomic tasks. This alignment comes with two practical costs. First, MLM pretraining supervises only the small fraction (≈15%) of positions that are masked (Devlin et al., 2019). The `[MASK]` token used in training also never appears at inference, which creates a train– test mismatch (BehnamGhader et al., 2024; Safari et al., 2025). Second, bidirectional attention-based encoders are expensive over long genomic windows, so they are hard to scale to the lengths at which distal regulation occurs. State-space encoders such as Caduceus ease this scaling cost, but they are still MLM-pretrained.

Causal decoders sit at the opposite corner of this trade-off. Models such as DNAGPT (Zhang et al., 2023) and HyenaDNA (Nguyen et al., 2023) supervise *every* token through next-token prediction, which makes pretraining dense and data-efficient. Some of their operators, such as the sub-quadratic Hyena long convolution, also scale more naturally to long contexts. What they lack is bidirectionality. Their representations are *directional*, and this geometry can be mismatched with tasks that need both flanks of a regulatory element at once, or a stable embedding of strand-equivalent sequences (Feng et al., 2025). In short, encoders give bidirectional context but pretrain less efficiently, while decoders pretrain densely and scale to long context but stay directional. Each family supplies only part of what a genomic representation model would ideally combine.

This motivates a different route. Rather than pretrain yet another bidirectional model from scratch, we ask whether an existing causal DNA decoder can be *converted* into a bidirectional encoder. Such a conversion would keep the dense supervision of causal pretraining, and where available its long-context scaling, while adding the bidirectional context that DNA calls for. In NLP, LLM2Vec (BehnamGhader et al., 2024) showed that decoder-only LLMs can be turned into strong sentence encoders in three steps: open bidirectional attention, repair the resulting mismatch with Masked Next-Token Prediction (MNTP), and refine embedding geometry with SimCSE-style contrastive learning (Gao et al., 2021). We adapt this recipe to genomics and introduce **DNA-LLM2Vec**, a staged decoder-to-encoder conversion framework for pretrained DNA decoders. We instantiate it on two architecturally distinct backbones: DNAGPT, a GPT-style Transformer decoder, and HyenaDNA, a long-context Hyena decoder. Using two backbones lets us test whether the recipe behaves consistently across distinct backbone and tokenisation regimes. Beyond the base LLM2Vec stages, we add contrastive views that encode the symmetries of DNA: strand invariance through reverse-complement pairs, and positional invariance within genomic windows through crop and local-shift pairs. Dropout SimCSE serves as a biology-agnostic baseline. We treat these invariances as task-dependent hypotheses rather than universal improvements, because a transformation that preserves one regulatory signal may erase another.

We validate DNA-LLM2Vec through a series of evaluation and ablation experiments on both backbones. The evaluation setup is described in section 3, the main results and ablations in section 4 and section 5, and the overall discussion in section 6.

## Methods

We adapt pretrained decoder-style DNA language models into bidirectional sequence encoders through a shared staged pipeline. For DNAGPT, the stages produce model variants M0–M6; for HyenaDNA, the corresponding variants are denoted H0–H6. The index has the same meaning in both families: 0 is the original causal decoder, 1 opens bidirectional context, 2 adds masked next-token adaptation, and 3–6 are dropout, reverse-complement, crop, and local-shift SimCSE variants. Stages 3–6 are parallel: each contrastive variant branches independently from the same Stage-2 checkpoint (M2/H2). Section 2 illustrates the overall design.

### Stage 1: Opening bidirectional context (M0/H0 → M1/H1)

#### DNAGPT

GPT-2 self-attention applies an upper-triangular causal mask so that position *t* attends only to positions 1, …, *t*. We remove it—setting the mask to **0**, disabling the causal flag, and zeroing the causal bias buffer—so no parameters change; M1 retains M0’s weights and differs only in attention behaviour.

#### HyenaDNA

HyenaDNA replaces self-attention with the Hyena operator, a gated *causal* long convolution, so there is no attention mask to edit. We instead make it bidirectional by running the *same* operator forward and on the reversed input, then merging the two streams. Writing *g*(·) for the causal Hyena operator and *R* for sequence reversal, we combine the forward stream *g*(**u**) and the backward stream *R g*(*R* **u**) with a learned projection **W**_*d*_ ∈ ℝ^*d×*2*d*^:

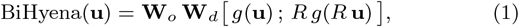

where [·; ·] is channel-wise concatenation and **W**_*o*_ is the operator’s existing output projection. Both directions share the same pretrained Hyena filter, so **W**_*d*_ is the only new parameter. We concatenate rather than sum so the projection can weight the two directions asymmetrically instead of averaging away orientation. Initialising 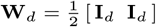 makes the patch-only checkpoint H1 a deterministic average of the forward and backward passes, so the H0→H1 comparison isolates the effect of bidirectional context rather than of an untrained projection. Stage 2 then adapts **W**_*d*_ with the backbone.

### Stage 2: Masked Next-Token Prediction (M1/H1 → M2/H2)

Standard MLM fine-tuning of a decoder creates a vocabulary mismatch: the model never saw a `[MASK]` token during autoregressive pretraining, so its `[MASK]` embedding is uninitialised and it has no learned behaviour for predicting masked positions (BehnamGhader et al., 2024). We instead use the Masked Next-Token Prediction (MNTP) objective of LLM2Vec. Given tokens **x** = (*x*_1_, …, *x*_*T*_), we mask a subset ℳ ⊂ {2, …, *T*} (with probability *p*_mask_). Let 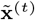 denote **x** with position *t* replaced by `[MASK]`. For each masked position *t*, the model predicts *x*_*t*_ from the hidden state at the *preceding* position, 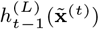

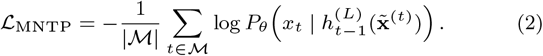

The key difference from standard MLM is where the prediction is read out: MLM predicts a masked token from its own position, whereas MNTP retains the causal convention that *t* − 1 predicts *t*. This reuses the pretrained next-token head, so after Stage 2 the *t* − 1 representation carries bidirectional context at the *token level*, while prediction still runs through weights the decoder already learned. **Mask token design**. The two backbones represent the mask differently, reflecting their tokenisation. Unlike the Llama/Mistral setting in LLM2Vec, DNAGPT has no neutral token analogous to : its BPE vocabulary is dominated by ATGC *k*-mers and structural special tokens, so we add a dedicated `[MASK]` token via `add_special_tokens` and resize the embedding matrix. During MNTP we update *only* this newly added row and keep the pretrained BPE embeddings fixed (mask-only; a full-embedding variant is compared in the Supplementary Material Section S6). HyenaDNA is tokenised at near single-nucleotide resolution, so for H2 we use *span-masked* adaptation with the same shifted-prediction objective. Contiguous nucleotide spans are replaced by a `[MASK]` token, and the preceding position predicts each masked nucleotide under bidirectional context. In both cases the `[MASK]` token is never an input during downstream evaluation (Steps 4 and 5), where sequences are encoded without masking.

### Stage 3: Contrastive learning (M2/H2 → M3–M6/H3–H6)

Whereas Stage 2 (MNTP) reshapes representations at the *token level*, it does not directly optimise the geometry of the *sequence-level* embedding space. We therefore add a SimCSE-style (Gao et al., 2021) contrastive stage that operates on pooled sequence embeddings. For each anchor **x**_*i*_ we build a single *positive* view 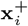 that is assumed to preserve the sequence’s biological meaning, while all other sequences in the mini-batch act as *negatives*. Let *h*_*θ*_(**x**) ∈ ℝ^*d*^ be the mean-pooled backbone representation; during training a two-layer projection head maps it to a 256-dimensional L2-normalised vector *f*_*θ*_(**x**) (discarded afterwards; downstream evaluations use *h*_*θ*_). We minimise a symmetric InfoNCE loss, the average of two directional terms *L*_*a*→*b*_ and *L*_*b*→*a*_ (the latter swapping anchor and positive). Writing *z*_*i*_ = *f*_*θ*_(**x**_*i*_) for the projected embedding,

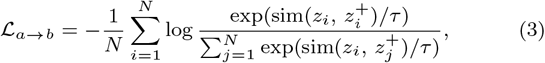

where sim is cosine similarity and *τ* a temperature.

The four variants below differ *only* in how the positive view 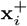 is constructed; the negatives are always the remaining in-batch sequences. Each positive-construction rule therefore encodes a distinct biological-equivalence assumption, which we test empirically rather than assume to hold universally.

#### M3/H3:Dropout augmentation

A biology-agnostic baseline. The positive view is the same sequence, 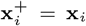, passed through the encoder twice with independent dropout, so the two views differ only by stochastic noise; it tests whether generic representation smoothing alone is enough.

#### M4/H4:Reverse-complement augmentation

Strand invariance. The positive view is the reverse complement of **x**_*i*_ (the complementary strand read 5’→3’). Because DNA is double-stranded, many sequence-level labels—enhancer, promoter, histone marks—should not depend on which strand is presented (Schiff et al., 2024), so matching a sequence to its reverse complement injects strand invariance without architectural change.

#### M5/H5:Crop augmentation

Invariance to window boundaries. From a long DNA sequence window we independently crop two overlapping sub-sequences of length *L*_crop_ (crop-start displacement up to *L*_crop_(1 − *r*_overlap_), so they share at least an *r*_overlap_ fraction) and treat them as the positive pair. This makes crop a natural fit for extended-context tasks such as EPI.

#### M6/H6:Local-shift augmentation

Fine-grained positional invariance. From a window of length *L*_crop_ + 2⌊*r*_shift_*L*_crop_⌋, the anchor starts after the left margin and the positive shifts its start by a random offset in [−*r*_shift_*L*_crop_, *r*_shift_*L*_crop_]. This is a finer-grained crop: the views are more similar than in M5, making the task easier but more regularising.

## Experimental Setup

The experiments follow a three-axis design (*backbone* × *adaptation recipe* × *evaluation protocol*). The following subsections detail the backbones and adaptation recipes, the data, and the evaluation protocols used to answer them.

### Backbones and adaptation

#### DNAGPT

We use DNAGPT (`dnagpt/human_gpt2-v1`) as the Transformer-decoder backbone: a GPT-2 decoder (Radford et al., 2019) with 105 M parameters, pretrained autoregressively (public human-only version) with Byte-Pair Encoding (BPE) tokenisation and a maximum context length of 1024 tokens.

#### HyenaDNA

We use HyenaDNA (*L*ongSafari/hyenadna-sma ll-32k-seqlen-hf) as the operator-based decoder backbone: a nucleotide-level long-context genomic language model built on the Hyena operator. This 32K checkpoint is used for all experiments (Sections 4.1–4.4), including the VEP probe, with evaluation inputs capped at 8192 bp (section A).

#### Adaptation

To preserve pretrained knowledge we adapt each backbone differently: DNAGPT uses Low-Rank Adaptation (LoRA) (Hu et al., 2022) on the GPT-2 `c_attn`/`c_proj` modules (query/key/value, attention-output and MLP projections), whereas HyenaDNA is fully fine-tuned. This matches the backbones at the recipe rather than the parameter level, avoiding a Hyena-specific adapter. Full training hyperparameters are in Appendix A.

### Data

#### Adaptation data

All decoder-to-encoder adaptation stages (MNTP/masked adaptation in Stage 2 and contrastive training in Stage 3) are trained on the hg38 (GRCh38) human reference genome (Schneider et al., 2017). We tile each chromosome into fixed-length windows and discard any window containing `N` or ambiguous bases. DNAGPT is adapted on non-overlapping 1024-token windows (≈4 kb under its BPE tokenisation), and HyenaDNA on 8192 bp nucleotide windows tiled with stride 4096; the crop and local-shift variants (M5/M6, H5/H6) draw their positive pairs from slightly longer source windows so that two offset crops still overlap (crop overlap ratio 0.5, local-shift ratio 0.1; section 2). Because adaptation is self-supervised, we do not hold out a genomic partition: a random 1% of windows is reserved only for monitoring the training objective, and all downstream evaluation uses the separate benchmark datasets below.

#### Evaluation benchmarks and task families

We evaluate on four public benchmark suites: Genomics Benchmarks (GB) (Grešová et al., 2023), the Nucleotide Transformer downstream tasks (NT) (Dalla-Torre et al., 2025), the Genome Understanding Evaluation (GUE) (Zhou et al., 2024), and its GUE+ Enhancer– Promoter Interaction extension (EPI) (Zhou et al., 2024). We use every task in each suite: all 8 GB tasks, all 18 NT tasks, 18 GUE tasks (excluding EMP yeast), and all 6 GUE+ EPI cell-line tasks. Rather than aggregate by source suite, our main analyses regroup these tasks into seven biological task families (EPI, Virus-COVID, histone/chromatin marks, TF-binding, promoter, splice, and local regulatory elements) plus one sequence-type control group (table 1); the suites serve only as data sources. The control group (coding-vs-intergenomic and human-vs-worm) is a near-ceiling sanity check rather than a regulatory family, so we exclude it from the task-family analysis and report its per-seed scores in the Supplementary Material Section S1. Per-task sequence lengths and split sizes are listed in sections C.1 to C.4, and the family-to-suite mapping with native metrics in table 1.

**Table 1.** Task-family grouping used for task-conditioned analyses. The grouping is used for aggregation and interpretation only; all models are evaluated on the original benchmark tasks.

| Task family | Source tasks | n | Biological structure / metric |
| --- | --- | --- | --- |
| EPI | GUE+ EPI cell-line tasks | 6 | extended enhancer–promoter pair relation / MCC |
| Virus-COVID | GUE Virus-COVID | 1 | extended viral single-sequence classification / macro-F1 |
| Histone/chromatin marks | NT histone/chromatin-mark tasks | 10 | local epigenomic mark prediction / MCC |
| TF-binding | GUE TF-Human and TF-Mouse tasks | 10 | local motif occupancy across TF/cell contexts / MCC |
| Promoter | GB Non-TATA, GUE core/300 promoter, NT promoter | 10 | short promoter/core-promoter classification / accuracy or MCC |
| Splice | NT splice (all, acceptor, donor) and GUE splice | 4 | position-sensitive splice recognition / MCC |
| Local regulatory element | GB enhancer/regulatory/OCR and NT enhancer tasks | 7 | local regulatory annotation / accuracy or MCC |
| Sequence-type controls | GB coding-vs-intergenomic and human-vs-worm | 2 | sequence-type / species discrimination / accuracy |

We additionally evaluate variant-effect prediction (VEP) (InstaDeep Ltd., 2024) on the causal-eQTL SNP task of the Genomics Long-Range Benchmark, where each SNP is labelled by whether the variant causally affects expression (fine-mapped GTEx eQTLs).

### Evaluation protocols

Our primary protocol is a frozen linear probe (Step 4), which tests frozen representation quality and underlies the architecture-aware task-conditioned analysis. We also fine-tune the adapted checkpoints end-to-end (Step 5) as a secondary check that they remain useful under task-specific supervision, and report it in the Supplementary Material Section S2&S3. Because the benchmark suites use different official metrics, task-family aggregates are computed in each task’s native metric as absolute deltas over the backbone-matched causal baseline (M0 for DNAGPT, H0 for HyenaDNA), averaged within each family; the baseline contributes its deterministic checkpoint value and each stochastic variant its mean across training seeds.

#### Linear probe (Step 4)

Backbone weights are frozen and a logistic regression classifier (L2 regularisation, *C* = 1.0, solver lbfgs) is fitted on top of mean-pooled, L2-normalised representations using scikit-learn (Pedregosa et al., 2011). We keep *C* at the scikit-learn default without per-task tuning, so the probe protocol is identical across all models and tasks. The lbfgs solver is deterministic given fixed data, so we report a single classifier fit per model–task pair.

#### Variant-effect prediction probe

VEP uses the same frozen-representation principle with a task-appropriate head and no backbone updates: for each SNP we encode its reference and alternate windows, form a paired feature (the concatenation [**e**_ref_; **e**_alt_] or the difference **e**_alt_ − **e**_ref_), standardise it, and classify with a RBF-SVM probe, reporting AUROC per distance-to-TSS stratum with paired bootstrap deltas against the causal baseline.

#### Reproducibility

Unless noted otherwise, all stochastic stages use three seeds (42–44), reported as mean ± std.

### Mechanistic motif-contact diagnostic

In addition to downstream classification, we use *in-silico* base-pair mutagenesis as an orthogonal diagnostic of whether adapted representations organise regulatory sequence structure around known motif instances; it is not used to train or select checkpoints. For each backbone variant, we compute position-by-position perturbation contact maps on held-out hg38 windows and test whether the top-scoring positive contact pairs are enriched for endpoints overlapping a strict IUPAC regulatory motif panel, comparing observed frequencies against distance-matched permutation backgrounds to control for the distance distribution of high-scoring contacts. We report Stouffer-combined *z* statistics and enrichment effect sizes, both pooled and group-specific (promoter-associated and TFBS motifs). The full protocol is given in Appendix B.

## Results

We organise the results from reusable frozen representations to transfer and mechanism. We first map task- and context-dependent frozen-probe behaviour, then test single-nucleotide variant-effect transfer and mechanistic geometry.

### Frozen linear probing

The Step 4 linear-probe results, aggregated by task family (fig. 2), give in each cell the mean task-level margin of an adapted variant over its backbone-matched causal baseline (M2–M6 vs M0 for DNAGPT, H2–H6 vs H0 for HyenaDNA), in each task’s native metric. From fig. 2, adaptation is not a universal improvement: the benefit depends on both the task’s biological structure and the backbone, and no single recipe dominates. Two patterns organise the matrix: crop contrastive learning is the shared optimum for the extended-context EPI and Virus-COVID families in both backbones, whereas the shorter regulatory and sequence-processing families are architecture-dependent. Per-seed task tables are in the Supplementary Material Section S1.

**Figure 1.**
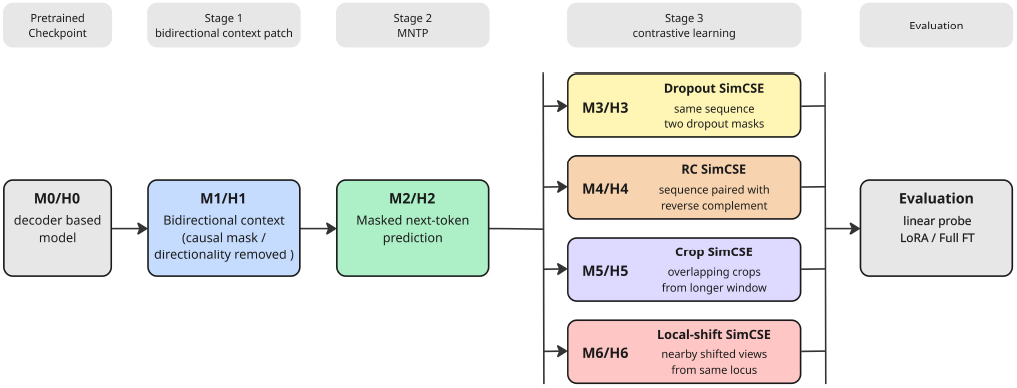
DNA-LLM2Vec adaptation pipeline: a pretrained causal DNA decoder is staged into a frozen bidirectional encoder (bidirectional patch, MNTP, then four parallel contrastive variants), instantiated on DNAGPT (M0–M6) and HyenaDNA (H0–H6).

**Figure 2.**
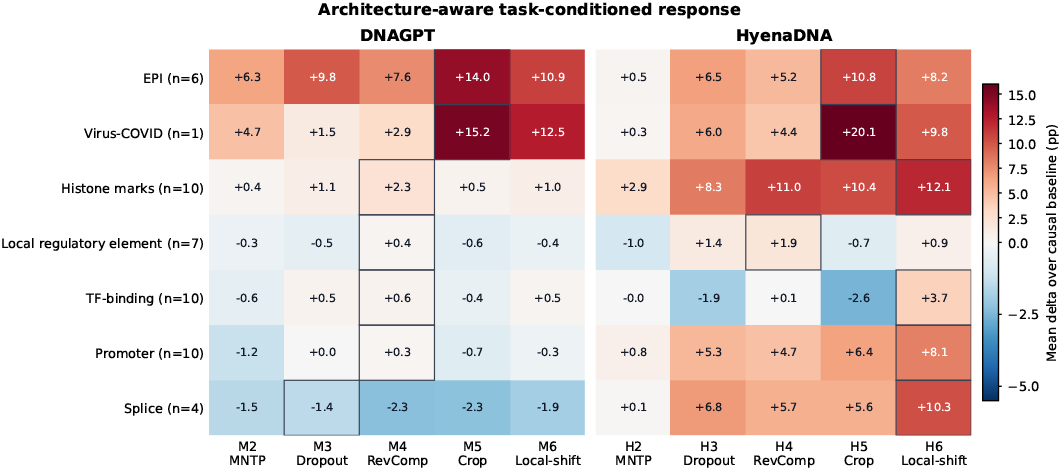
Task-family by variant response under frozen linear probing. Cells show the mean task-level margin over the backbone-matched causal baseline, in percentage points of each task’s native metric. Positive values indicate improvement; negative values indicate transfer cost.

Because the clearest positive families are EPI and Virus-COVID, we ask whether the gains track the sequence context available to the probe (fig. 3). For DNAGPT, the best adapted margin grows with sequence length (Pearson *r* = 0.64, *n* = 48): all six EPI tasks improve by +7.8 to +22.6 pp MCC and Virus-COVID by +15.2 pp macro-F1. HyenaDNA also improves on all EPI tasks (+9.2 to +13.9 pp MCC) and Virus-COVID (+20.2 pp macro-F1), but its length trend is weaker (*r* = 0.36, *n* = 48), because several short histone, promoter and splice tasks also gain substantially.

**Figure 3.**
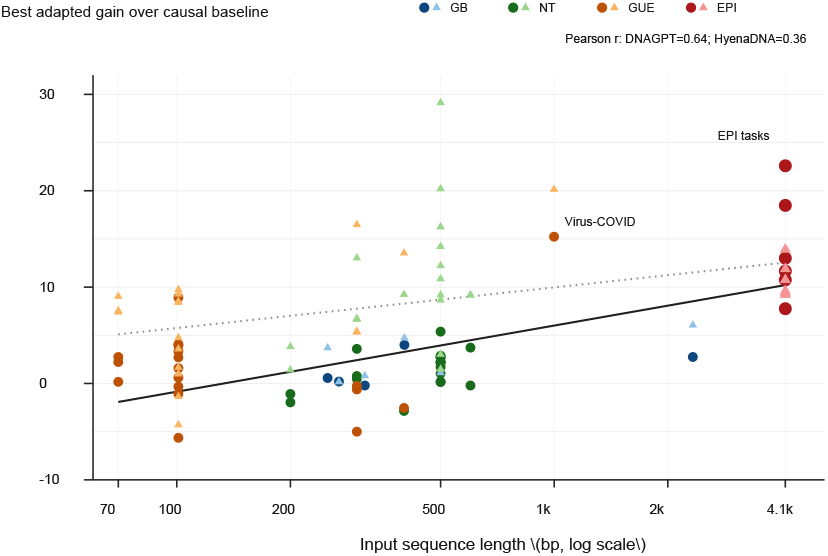
Gain of the best per-task adaptation variant (maximum over variants) over the causal baseline across Step 4 biological tasks; an upper envelope, since the best variant is task-dependent. Each point is one task, with values reported as percentage-point differences in the task metric. Colours denote benchmark suites; within each suite, • + denote DNAGPT, while ▲ + · · · denote HyenaDNA.

#### Short-context task-family trade-offs

Among the short families— ordered biologically from motif-proximal TF-binding, through cis-regulatory local-element tasks, to broader histone/chromatin-state readouts—agreement across families suggests transfer beyond a single assay type. For DNAGPT (BPE tokens over genomic *k*-mers), reverse-complement contrastive learning (M4) gives the strongest mean response for histone marks, local regulatory elements, TF-binding and promoter tasks, though the margins are far smaller than in EPI or Virus-COVID. Splice is the clearest negative-transfer family: every adapted DNAGPT variant falls below M0, most under reverse-complement, crop and local-shift views.

The HyenaDNA panel differs. With nucleotide-level tokenisation and longer context, it responds positively on histone, promoter and splice families: local shift (H6) gives the largest promoter and splice margins, while crop (H5) stays strongest for the extended-context EPI and Virus-COVID tasks. TF-binding stays near zero on average and is clearly positive only under H6, and local regulatory elements remain more mixed than the histone or promoter families.

### Variant-effect prediction (VEP)

We probe the frozen HyenaDNA encoders on the causal-eQTL VEP task using the Caduceus-matched protocol of section 3.3: paired reference/alternate SNP windows are mean-pooled, classified by an RBF-SVM, and scored by AUROC within three distance-to-TSS strata (table 2, mean over three seeds). We restrict this analysis to HyenaDNA to maintain nucleotide-level correspondence between the reference and alternate windows and to avoid changes in BPE segmentation following a single-base substitution, which under DNAGPT’s tokenisation re-segments the surrounding sequence so that the two token windows no longer align position-for-position (Zhou et al., 2024), whereas nucleotide-level tokenisation, used here and by Caduceus (Schiff et al., 2024), keeps the probe well defined at single-base resolution. The two adaptation stages help specifically at distance: H0 is best only within 0–30 kb, whereas MNTP (H2) is best in both distal strata (0.668 and 0.604 AUROC), with bidirectional patching (H1) already improving over H0 there. The contrastive variants (H3–H6) give no further gain, indicating that for causal-eQTL transfer the benefit comes from bidirectional context and MNTP rather than from the contrastive variants.

**Table 2.** Frozen VEP probe (RBF-SVM) on the Genomics Long-Range Benchmark causal-eQTL SNP task, by distance-to-TSS stratum (AUROC, mean± std over 3 seeds; HyenaDNA only). Best per column in bold.

| Variant | 0–30 kb | 30–100 kb | >100 kb |
| --- | --- | --- | --- |
| H0 (causal) | **0.690**±0.003 | 0.633±0.002 | 0.569±0.000 |
| H1 (bidir. patch) | 0.677±0.003 | 0.647±0.002 | 0.586±0.000 |
| H2 (MNTP) | 0.687±0.003 | **0.668**±0.006 | **0.604**±0.015 |
| H3 (dropout) | 0.638±0.011 | 0.640±0.006 | 0.586±0.013 |
| H4 (reverse-complement) | 0.665±0.005 | 0.620±0.012 | 0.563±0.020 |
| H5 (crop) | 0.658±0.001 | 0.630±0.014 | 0.576±0.011 |
| H6 (local-shift) | 0.669±0.004 | 0.591±0.008 | 0.543±0.030 |

### Reflection-equivariance diagnostics

The frozen-probe results show that converting HyenaDNA changes its downstream behaviour but not *how*. We therefore probe the converted backbone directly: reversing the input (a pure positional flip *R*, not a reverse complement) should not collapse a model that has genuinely learned sequence orientation. On held-out hg38 windows we measure three quantities per variant (mean over three seeds; full table in Supplementary Section S13), writing *H*(**x**) for the pre-pooling embedding sequence and *f* (**x**) for its mean pool: token-level equivariance tok eq, the mean per-position cosine between *R H*(**x**) and *H*(*R* **x**); pooled invariance pooled_refl = cos(*f* (**x**), *f* (*R***x**)); and anisotropy anchor, the mean cosine between two distinct window embeddings. For all three a *lower* value is the informative direction. We report HyenaDNA only, since its symmetrically initialised bidirectional patch is an even, reflection-equivariant operator that training must break, whereas DNAGPT’s absolute position embeddings and attention cannot become orientation-blind when the causal mask is removed.

Three patterns follow. *The conversion behaves as designed* : the untrained patch H1 is exactly reflection-equivariant (tok eq = pooled refl = 1.00), and MNTP begins to break this symmetry (tok eq = 0.74 for H2). *The contrastive stage de-anisotropises the space*: anchor falls from 0.95–0.99 (H0–H2) to 0.29–0.45 (H3– H6), consistent with improved embedding uniformity and helping explain the frozen-probe gains in section 4.1. *The views differ mechanistically* : dropout (H3), crop (H5) and local shift (H6) break token-level equivariance hard (tok eq = 0.11–0.19, far below the causal 0.86), whereas reverse-complement (H4) retains the most reflection structure (tok eq = 0.59), as expected since it explicitly trains invariance to the reverse complement, the symmetry closest to the positional reversal measured here.

### Contact-map diagnostics

The task-family heatmap measures downstream utility; we now ask two mechanistic questions: whether the adaptation recipes organise perturbation structure around known regulatory motifs, and whether reverse-complement training makes that structure strand-consistent. Both use the base-pair-mutagenesis motif-contact diagnostic of section 3.4.

#### Motif-contact enrichment

For DNAGPT, the M0→M1→M2 chain is informative: opening bidirectional context alone reduces the pooled motif signal (Stouffer *z*: 27.9 → 8.2), and MNTP then raises it above baseline (8.2 → 34.2). Among contrastive variants (initialised from M2), reverse-complement SimCSE (M4) gives the strongest motif-contact signal (*z* = 41.6, mean enrichment 1.23), while dropout, crop and local-shift stay below M2—contrastive adaptation is not uniformly beneficial; the invariance encoded by the positive pair matters.

#### Reverse-complement consistency

We next test whether the reverse-complement objective yields strand-consistent perturbation geometry—a probe of biological strand symmetry, distinct from the pure positional reversal in section 4.3. The effect is specific and large in both backbones. M4 raises RC consistency from 0.073 ± 0.022 (M0) to 0.271 ± 0.067 (Δ = +0.199, paired Wilcoxon *q* = 1.07 × 10^*−*6^, 23/24 windows improving; fig. 4(A)), exceeding every other DNAGPT variant (all *q <* 10^*−*6^). In HyenaDNA, H4 is again strongest, raising consistency from 0.224±0.051 (H0) to 0.935±0.018 (Δ = +0.711, *q* = 1.79 × 10^*−*7^, all 24 windows; fig. 4(A)), again above every other variant (all *q <* 10^*−*6^). Per-comparison paired tests for both backbones are in the Supplementary Material Section S16&S17. Reverse-complement contrastive learning therefore does not merely improve a downstream score; it makes base-pair-level contact structure strand-consistent.

**Figure 4.**
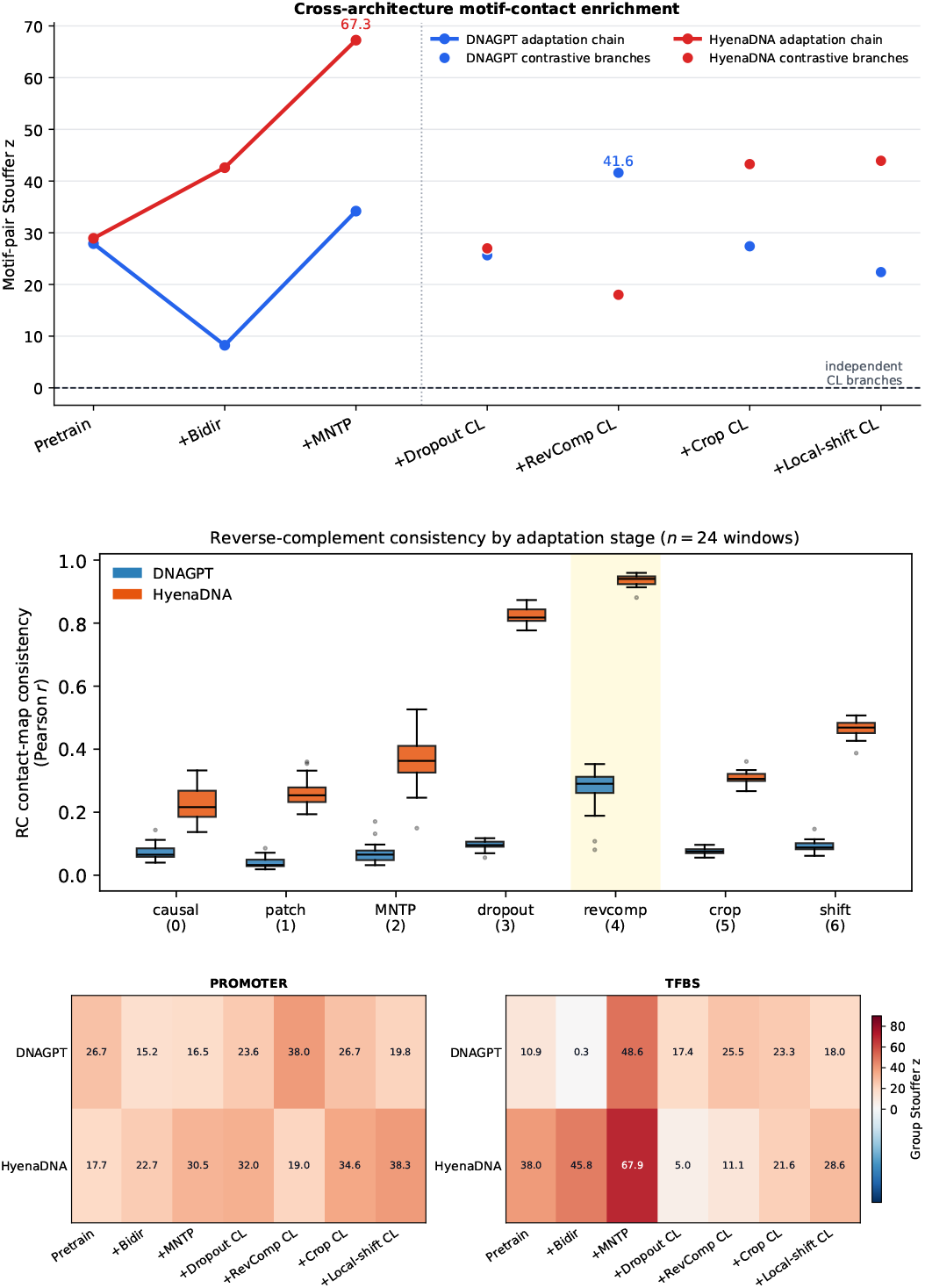
Contact-map diagnostics. (A) Pooled motif-pair Stouffer *z*. (B) Reverse-complement consistency of base-pair mutagenesis contact maps by adaptation stage (Pearson correlation across *n* = 24 hg38 windows; blue DNAGPT, orange HyenaDNA); reverse complement (M4/H4) gives the largest, most consistent increase in both backbones.(C) Promoter and TFBS group-level Stouffer *z*. Effect sizes track the pooled *z* (full per-variant enrichment in the Supplementary Material Section S14).

#### Cross-architecture comparison

Repeating the motif-contact analysis across the matched H0–H6 gives a two-way grid: seven recipes on two very different backbones. Adaptation strengthens motif-contact organisation in both (fig. 4(B, C)), but the peak stage is not shared. DNAGPT peaks at reverse-complement (M4, above); HyenaDNA peaks right after MNTP, its pooled signal rising *z* = 28.9 → 42.6 → 67.3 (H0→H1→H2, enrichment 1.38 at H2), with contrastive branches lower but positive (H6 *z* = 43.9, H5 *z* = 43.3). Group-level results separate pooled from motif-type effects: for DNAGPT, M4 is strongest for promoters (*z* = 38.0) while TFBS peaks at MNTP (M2 *z* = 48.6); for HyenaDNA, H6 is strongest for promoters (*z* = 38.3) while TFBS again peaks at MNTP (H2 *z* = 67.9). Masked adaptation thus dominates the pooled and TFBS signals, while the strongest promoter view is architecture-dependent (M4 for DNAGPT, H6 for HyenaDNA).

#### Diagnostics vs downstream optima

Motif and EPI optima differ because different invariances organise different geometry: in DNAGPT, reverse complement (M4) organises local strand-symmetric motif structure while crop (M5) helps long-window tasks; in HyenaDNA, MNTP (H2) drives pooled and TFBS organisation, local shift (H6) sharpens promoters, and crop (H5) remains best for EPI. The two contact-level diagnostics also decouple differently across backbones: in DNAGPT, M4 gives both the strongest strand consistency and the strongest contrastive motif-pair enrichment, whereas in HyenaDNA H4 gives the strongest strand consistency but the *lowest* pooled enrichment of the contrastive variants (*z* = 18.0), with pooled organisation peaking at H2.

## Ablation and controls

We conduct three ablation studies on the frozen linear-probe results, varying the pretrained backbone, the crop variant’s initialisation, and the pooling operator. We report each as EPI (Avg-EPI MCC) and the mean over all seven task families (Fam-mean).

### Value of pretraining

We re-run the Stage-2/3 adaptation from a *randomly initialised* backbone (table 3). The pretrained backbone is required for the broad multi-family competence: the best Fam-mean falls from 45.28 to 32.90 for DNAGPT and from 40.94 to 36.20 for HyenaDNA. Its role in the EPI signal is architecture-dependent. Crop still lifts EPI on a random backbone in both models (by +8.40 for DNAGPT and +8.36 for HyenaDNA over the random causal baseline), but the ceiling differs sharply: DNAGPT’s random crop reaches 23.39 Avg-EPI MCC, close to its pretrained 27.46, whereas HyenaDNA’s random crop reaches only 15.24 against a pretrained 26.65, with a random causal baseline that is itself very low (6.88). Pretraining is therefore essential for the broad family response in both backbones and for HyenaDNA’s EPI signal, while DNAGPT’s EPI gain is largely recoverable from a random backbone through crop.

**Table 3.**
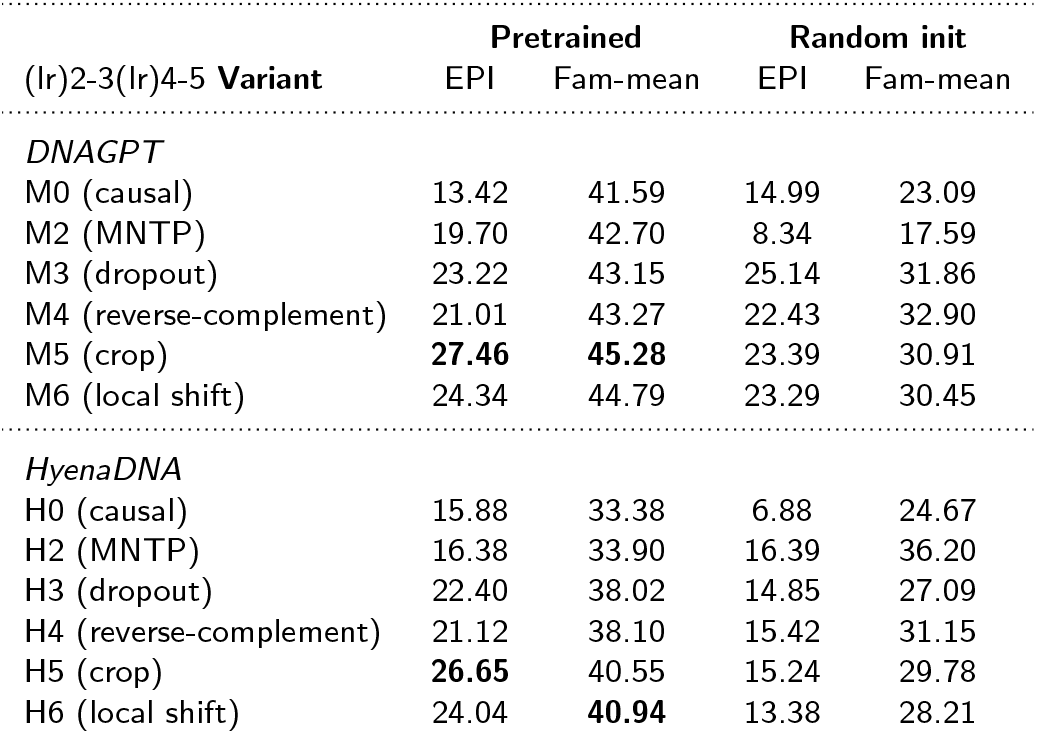
Value of pretraining. Frozen family scores (%) when the Stage-2/3 adaptation is applied to a *randomly initialised* backbone versus the pretrained backbone used in the main results (single seed, r1). EPI is Avg-EPI MCC; Fam-mean is the mean over the seven task families.

### Crop-variant initialisation

The crop variant (M5/H5) branches from the MNTP checkpoint (stage 2) in the main pipeline. We compare branching instead from the causal backbone (stage 0) or the bidirectional patch (stage 1) (table 4). Branching from stage 2 gives the best Fam-mean in both backbones (45.28 vs. 44.73 and 43.91 for DNAGPT; 40.55 vs. 38.28 and 38.80 for HyenaDNA), while EPI is nearly unchanged by the branch point (25.93–27.46 for DNAGPT; 26.15–26.65 for HyenaDNA). The crop EPI gain is thus robust to where adaptation starts, while the staged initialisation adds a modest Fam-mean improvement; we discuss the complementary roles of the stages in section 6.

**Table 4.** Crop-variant initialisation. Frozen family scores (%) when the crop variant branches from stage 0, stage 1, or stage 2 (the main M5/H5), single seed.

| Branches from | EPI | Fam-mean |
| --- | --- | --- |
| *DNAGPT (M5)* |  |  |
| Stage 0 (causal) | 25.93 | 43.91 |
| Stage 1 (bidir. patch) | 27.44 | 44.73 |
| Stage 2 (MNTP, main) | **27.46** | **45.28** |
| *HyenaDNA (H5)* |  |  |
| Stage 0 (causal) | 26.57 | 38.80 |
| Stage 1 (bidir. patch) | 26.15 | 38.28 |
| Stage 2 (MNTP, main) | **26.65** | **40.55** |

### Pooling

We replace mean pooling over token hidden states with an attention-weighted pooling alternative in the frozen probe (table 5). The per-variant ordering is preserved (crop, M5/H5, strongest on EPI; M5/M6 and H5/H6 on Fam-mean), and Fam-mean stays within about 1 pp of mean pooling (DNAGPT M5 45.28 vs. 46.08; HyenaDNA H5 40.55 vs. 40.91). The frozen-probe conclusions do not depend on the pooling operator, so we keep the simpler mean pooling throughout.

**Table 5.** Pooling robustness. Frozen family scores (%) under mean versus attention-weighted pooling (mean over three seeds). EPI is Avg-EPI MCC; Fam-mean is the mean over the seven task families.

| (lr)2-3(lr)4-5 Variant | Mean pool |  | Weighted pool |  |
| --- | --- | --- | --- | --- |
|  | EPI | Fam-mean | EPI | Fam-mean |
| *DNAGPT* |  |  |  |  |
| M2 (MNTP) | 19.70 | 42.70 | 20.02 | 42.22 |
| M3 (dropout) | 23.22 | 43.15 | 23.57 | 42.79 |
| M4 (reverse-complement) | 21.01 | 43.27 | 21.77 | 43.98 |
| M5 (crop) | **27.46** | **45.28** | **29.05** | **46.08** |
| M6 (local shift) | 24.34 | 44.79 | 25.06 | 45.14 |
| *HyenaDNA* |  |  |  |  |
| H2 (MNTP) | 16.38 | 33.90 | 16.90 | 33.63 |
| H3 (dropout) | 22.40 | 38.02 | 23.46 | 38.15 |
| H4 (reverse-complement) | 21.12 | 38.10 | 23.08 | 38.54 |
| H5 (crop) | **26.65** | 40.55 | **28.75** | 40.91 |
| H6 (local shift) | 24.04 | **40.94** | 25.61 | **41.19** |

**Table 6.** DNAGPT training and evaluation hyperparameters used for the main experiments. Full command-line examples are provided in the public repository README.

| Stage | Input/window | Optimisation | Batch | LR | Other settings |
| --- | --- | --- | --- | --- | --- |
| Stage 2 MNTP (M2) | 1024 tokens | LoRA $r = 16$, $\alpha = 32$, drop 0.05 | 24×2 | 1e−4 | mask prob 0.15; warmup 500; 1 epoch; mask-only embeddings |
| Stage 3 dropout (M3) | 1024 tokens | LoRA $r = 16$, $\alpha = 32$, drop 0.05 | 64×2 | 1e−4 | dropout positives; $\tau = 0.05$; 3000 steps |
| Stage 3 revcomp (M4) | 1024 tokens | LoRA $r = 16$, $\alpha = 32$, drop 0.05 | 64×2 | 1e−4 | reverse-complement positives; $\tau = 0.05$; 1 epoch |
| Stage 3 crop (M5) | chunk 4096 bp | LoRA $r = 16$, $\alpha = 32$, drop 0.05 | 64×2 | 1e−4 | overlap 0.5; $\tau = 0.05$; 1 epoch |
| Stage 3 local shift (M6) | chunk 4096 bp | LoRA $r = 16$, $\alpha = 32$, drop 0.05 | 64×2 | 1e−4 | shift 0.1; $\tau = 0.05$; 1 epoch |
| Step 4 linear probe | EPI crop 4096 bp | frozen backbone + logistic regression | 128 | – | mean pooling; L2-normalised embeddings |
| Step 5 full FT | 1024 tokens | full fine-tuning | 8 | 3e−5 | GUE+ EPI: 20 epochs; warmup 50; gradient checkpointing |

**Table 7.** HyenaDNA training and evaluation hyperparameters used for the cross-architecture replication experiments.

| Stage | Input/window | Optimisation | Batch | LR | Other settings |
| --- | --- | --- | --- | --- | --- |
| Stage 2 MNTP (H2) | 8192 bp, stride 4096 | full-model span-masked adaptation | 32×2 | 1e−4 | mask prob 0.15; span 3–20 bp; warmup 500; 1 epoch |
| Stage 3 dropout (H3) | 8192 bp, stride 4096 | full-model contrastive adaptation | 16×2 | 1e−5 | dropout positives; $\tau = 0.05$; 1 epoch |
| Stage 3 revcomp (H4) | 8192 bp, stride 4096 | full-model contrastive adaptation | 16×2 | 1e−5 | reverse-complement positives; $\tau = 0.05$; 1 epoch |
| Stage 3 crop (H5) | chunk 8192 bp, stride 4096 | full-model contrastive adaptation | 16×2 | 1e−5 | overlap 0.5; $\tau = 0.05$; 1 epoch |
| Stage 3 local shift (H6) | chunk 8192 bp, stride 4096 | full-model contrastive adaptation | 16×2 | 1e−5 | shift 0.1; $\tau = 0.05$; 1 epoch |
| Step 4 linear probe | EPI crop 8192 bp | frozen backbone + logistic regression | 128 | – | mean pooling; L2-normalised embeddings |
| Step 5 full FT | 8192 bp | full fine-tuning | 4 | 3e−5 | GUE/GUE+ only; 20 epochs; warmup 50; gradient checkpointing |

**Table 8.** Approximate adaptation wall-clock cost on one NVIDIA A100 40GB GPU. Times are rounded estimates from the main runs and exclude downstream Step 4/5 evaluation. Stage 1 is a structural patch; Stage 2/3 costs are per seed/run for the listed training length.

| Backbone | Stage / variant | Training length | Wall-clock |
| --- | --- | --- | --- |
| DNAGPT | Stage 1 bidirectional patch (M1) | – | ~10 min |
| DNAGPT | Stage 2 MNTP (M2) | 1 epoch | ~2.5 h |
| DNAGPT | Stage 3 dropout (M3) | 1 epoch | ~10 h |
| DNAGPT | Stage 3 revcomp (M4) | 1 epoch | ~6 h |
| DNAGPT | Stage 3 crop (M5) | 1 epoch | ~5.5 h |
| DNAGPT | Stage 3 local shift (M6) | 1 epoch | ~8 h |
| HyenaDNA | Stage 1 bidirectional patch (H1) | – | ~10 min |
| HyenaDNA | Stage 2 masked adaptation (H2) | 1 epoch | ~3 h |
| HyenaDNA | Stage 3 dropout (H3) | 1 epoch | ~9.75 h |
| HyenaDNA | Stage 3 revcomp (H4) | 1 epoch | ~8.15 h |
| HyenaDNA | Stage 3 crop (H5) | 1 epoch | ~15 h |
| HyenaDNA | Stage 3 local shift (H6) | 1 epoch | ~13.4 h |

Together, these three controls support the design: (i) the frozen gains mostly reorganise a pretrained backbone rather than learn it from scratch, though a random backbone still recovers part of the gain (table 3), so pretraining and adaptation contribute jointly; (ii) the staged pipeline gives the crop variant a small additional edge over its shortcuts; and (iii) the frozen-probe conclusions do not depend on the pooling operator.

## Discussion

Our results are best read as a regime map: decoder-to-encoder conversion injects specific biological invariances, and each helps only where it matches the task family and the backbone. This echoes the original LLM2Vec observation that embedding adaptation helps geometry-sensitive tasks more than standard classification; here the relevant axis is biological structure rather than the benchmark of origin. The extended-context regime is the clearest case, and its cross-backbone size gap invites a mechanistic reading. The larger frozen EPI margin for DNAGPT than HyenaDNA need not mean the Transformer is intrinsically more long-range capable: DNAGPT’s causal baseline starts weaker (13.4% vs 15.9% Avg-EPI MCC), leaving more for bidirectional repair and crop-based invariance to recover, whereas HyenaDNA is already a long-context nucleotide model. We keep this interpretation at the sequence-window level, since the GUE+ EPI files provide sequences but not genomic coordinates.

The short families expose an interaction between contrastive view and tokenisation. Splice recognition depends on nucleotide-precise donor and acceptor signals that BPE can merge into multi-base tokens, consistent with DNAGPT’s splice regressions and HyenaDNA’s splice gains; short core-promoter windows likewise expose few BPE tokens for crop or shift to act on. Yet neither backbone shows a uniform TF-binding gain, so motif resolution alone does not explain the short-task map. We cannot fully separate tokenisation from sequence operator or context length without matched controls, so these remain plausible contributors rather than a proven mechanism.

The base-pair-mutagenesis diagnostics are useful precisely because they are orthogonal to classification. Reverse-complement adaptation most directly imprints strand-consistent contact geometry, yet motif enrichment does not predict the best downstream view (crop, not reverse complement, wins EPI), and the two diagnostics decouple differently across backbones.

The ablation controls sharpen the causal reading: the gains reorganise a pretrained backbone rather than emerge from the objectives alone, most stringently for HyenaDNA’s EPI signal, while full supervision reshapes rather than preserves the frozen family response. Crop recovers most of the EPI gain even from an earlier start (section 5.2), but does not make MNTP redundant: the two objectives act at different representation scales. MNTP repairs the token- and position-level mismatch left by opening bidirectional attention, whereas contrastive learning directly optimises the pooled *sequence-level* geometry that frozen sequence-level classification such as EPI relies on; MNTP’s own contribution instead surfaces in position-resolved, token-sensitive diagnostics, namely the motif-contact analysis and the distal-VEP improvements at H1/H2. The method’s value therefore lies in frozen embedding geometry.

### Limitations

We evaluate one human-only DNAGPT scale and one HyenaDNA backbone, so comparisons with multi-species encoders are contextual rather than controlled, and we cannot isolate tokenisation from operator or context-length effects. The pretraining and crop-start controls are single-seed and should be read as indicative of the main effects rather than as bariance-quantified estimates. The GUE+ EPI benchmark is sequence-level rather than coordinate-resolved, and small families (Virus-COVID, splice) are descriptive observations requiring broader validation.

## Conclusion

DNA-LLM2Vec turns pretrained causal DNA decoders into frozen bidirectional encoders whose value is task- and architecture-conditioned rather than universal. The broader implication is methodological: genomic embedding methods should select or learn contrastive views by both task structure and architecture, and report positive and negative transfer regimes rather than a single aggregate score.

## Data availability

All code is available at `semi-nothing/dna-llm2vec`. The human reference genome (hg38) was obtained from the UCSC Genome Browser. GUE benchmark data were obtained from the DNABERT-2 GitHub repository (`MAGICS-LAB/DNABERT_2`); Genomics Benchmarks data via the `genomic-benchmarks` Python package (Grešová et al., 2023); and Nucleotide Transformer downstream-task data from Hugging Face (`InstaDeepAI/nucleotide_transformer_downstream_tasks`). All datasets are publicly available. Additional per-seed tables, ablations, and diagnostic analyses are provided in the Supplementary Material.

## Author contributions statement

Y.L. conceived the study and designed the method; Y.L. implemented the pipeline and ran the experiments; Y.L., K.Z., S.L. and J.S. analysed the results; Y.L. and K.Z. wrote the manuscript; Y.G., G.W. and M.P. supervised the work. All authors reviewed and approved the final manuscript.

## Acknowledgments

We thank Apocrita, Queen Mary University of London’s high-performance computing cluster, and its ITS Research team for providing and maintaining the computing resources used in this work (King et al., 2017).

During the preparation of this work we used Claude and ChatGPT to assist with writing parts of the analysis code, processing and tabulating experimental results, editing the language of text we had written, and checking internal consistency across the manuscript. We reviewed all such output and verified it against the underlying data and code, and we take full responsibility for the content of this article. The study design, the experiments, the interpretation of results and all scientific claims are our own, and no part of the manuscript was drafted by AI from a prompt. Further details are given in the Supplementary Material.

## Training Hyperparameters

*Precision, input length and hardware*. DNAGPT adaptation and downstream embedding evaluation use bfloat16; the base-pair-mutagenesis contact-map diagnostics (section 3.4) instead load DNAGPT checkpoints in float32 to avoid precision-sensitive perturbation artefacts. HyenaDNA runs in float32 throughout, because the Hyena operator casts activations to float32 internally regardless of input dtype. Because all performance comparisons are made between variants of the same backbone, these per-backbone precision choices do not affect our conclusions. For the short-context benchmarks we cap the 32K HyenaDNA inputs at 8192 bp; every sequence there is shorter than 8192 bp (the longest is the 5000 bp EPI input), so the cap truncates nothing while allowing a larger training batch size. All DNAGPT experiments use a single NVIDIA A100-PCIE-40GB GPU. All adaptation stages are trained on hg38, excluding windows that contain ambiguous nucleotides.

## Base-Pair Mutagenesis Motif-Enrichment Protocol

For the contact-map diagnostics reported in the Results and Supplementary Material, we used the same held-out hg38 windows for all variants within each backbone and the same 24-window set for the cross-architecture comparison. The reported DNAGPT motif-enrichment and reverse-complement-consistency contact maps were generated with the checkpoints loaded in float32. For each window, we computed base-pair mutagenesis effects by substituting each position with alternative nucleotides, measuring the resulting change in the mean-pooled sequence embedding, and converting these perturbation effects into a symmetric position-by-position contact score matrix. Concretely, for each mutable position *i* and alternative base *b* we record the pooled-embedding delta 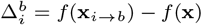, where *f* is the L2-normalised mean-pooled embedding and **x**_*i*→*b*_ is the sequence with position *i* substituted by base *b*. The contact score between positions *i* and *j* is the cosine similarity of these deltas, averaged over the alternative bases *B*,

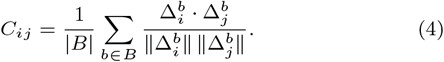

We then zero its diagonal, and apply average-product correction (APC). We used the positive contact score for the primary analysis, because motif co-occurrence is expected to increase same-direction perturbation structure; absolute-score analyses are treated as sensitivity checks.

Motifs were scanned directly on the stored input sequence using a fixed panel of eleven canonical regulatory consensus sequences in IUPAC notation (table 9): four core promoter elements and seven transcription-factor binding consensus sequences. Scanning used the reverse complement as well as the forward strand unless otherwise stated. We restrict the panel to consensus sequences of at least six positions, because shorter patterns match held-out hg38 windows at rates that make the distance-matched background comparison uninformative; this excludes otherwise canonical but short elements such as the DPE, the CCAAT box, the CTCF core and the ETS core.

**Table 9.** Regulatory motif panel used for the base-pair mutagenesis contact-enrichment analysis. Patterns are IUPAC consensus sequences, scanned on both strands with no flank expansion in the main analysis.

| Motif | IUPAC pattern | Length | Group |
| --- | --- | --- | --- |
| TATA box | TATAWAWR | 8 | promoter |
| Initiator | YYANWYY | 7 | promoter |
| BRE<sub>u</sub> | SSRCGCC | 7 | promoter |
| GC box | GGGCGG | 6 | promoter |
| E-box | CACGTG | 6 | TFBS |
| AP-1 | TGASTCA | 7 | TFBS |
| CRE | TGACGTCA | 8 | TFBS |
| NF-κB | GGGRNNYYCC | 10 | TFBS |
| GATA | WGATAR | 6 | TFBS |
| RUNX | TGTGGT | 6 | TFBS |
| YY1 | CCATNTT | 7 | TFBS |

**Table 10.**
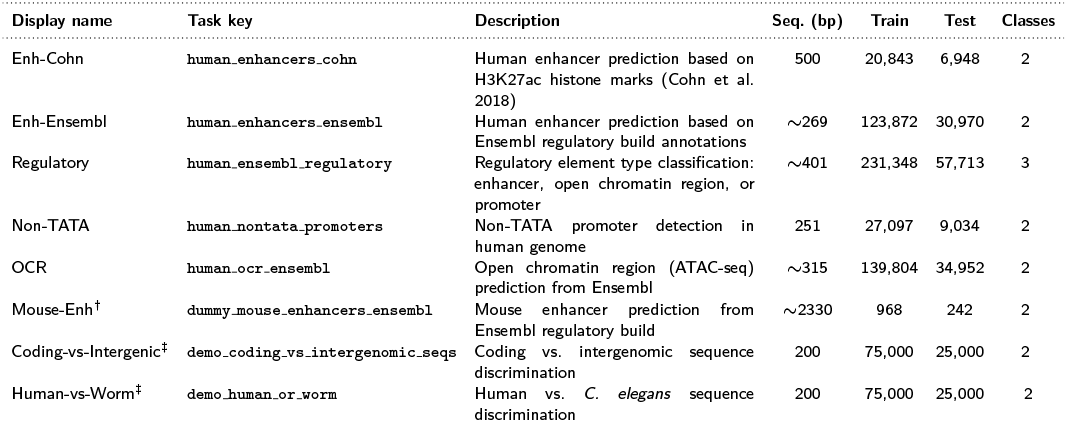
Genomics Benchmarks (GB) task details (all 8 tasks). Counts are the train/test split sizes from the `genomic-benchmarks` package; Seq. length is the median sequence length in base pairs. ^†^ mouse-derived; ^‡^ sequence-type control tasks (the remaining tasks are human regulatory tasks).

**Table 11.** Nucleotide Transformer downstream task details (all 18 tasks). Histone/chromatin-mark tasks use human ENCODE ChIP-seq data (GRCh38); splice-site tasks derive from the Spliceator dataset. Sequence lengths follow the released NT downstream datasets.

| Display name | Task key | Description | Seq. (bp) | Train | Test | Classes |
| --- | --- | --- | --- | --- | --- | --- |
| *Histone / chromatin marks (10 tasks)* |  |  |  |  |  |  |
| H3 | H3 | Core histone H3 occupancy | 500 | 13,468 | 1,497 | 2 |
| H4 | H4 | Core histone H4 occupancy | 500 | 13,140 | 1,461 | 2 |
| H3K4me1 | H3K4me1 | H3K4 monomethylation | 500 | 28,509 | 3,168 | 2 |
| H3K4me2 | H3K4me2 | H3K4 dimethylation | 500 | 27,614 | 3,069 | 2 |
| H3K4me3 | H3K4me3 | H3K4 trimethylation (active promoter) | 500 | 33,119 | 3,680 | 2 |
| H3K9ac | H3K9ac | H3K9 acetylation (active promoter/enhancer) | 500 | 25,003 | 2,779 | 2 |
| H3K14ac | H3K14ac | H3K14 acetylation | 500 | 29,743 | 3,305 | 2 |
| H3K36me3 | H3K36me3 | H3K36 trimethylation (active transcription) | 500 | 31,392 | 3,488 | 2 |
| H3K79me3 | H3K79me3 | H3K79 trimethylation | 500 | 25,953 | 2,884 | 2 |
| H4ac | H4ac | H4 acetylation | 500 | 30,685 | 3,410 | 2 |
| *Promoter (3 tasks)* |  |  |  |  |  |  |
| Promoter-All | promoter.all | Promoter vs. non-promoter | 300 | 53,276 | 5,920 | 2 |
| Promoter-TATA | promoter.tata | TATA-box promoter subset | 300 | 5,509 | 621 | 2 |
| Promoter-noTATA | promoter.no.tata | TATA-box-free promoter subset | 300 | 47,767 | 5,299 | 2 |
| *Enhancer (2 tasks)* |  |  |  |  |  |  |
| Enhancers | enhancers | Enhancer detection | 200 | 14,968 | 400 | 2 |
| Enhancer-Types | enhancers.types | Enhancer type classification | 200 | 14,968 | 400 | 3 |
| *Splice site (3 tasks)* |  |  |  |  |  |  |
| Splice-All | splice_sites.all | Donor, acceptor, or neither (background) | 400 | 27,000 | 3,000 | 3 |
| Splice-Acceptor | splice_sites.acceptors | Acceptor site detection | 600 | 19,961 | 2,218 | 2 |
| Splice-Donor | splice_sites.donors | Donor site detection | 600 | 19,775 | 2,198 | 2 |

**Table 12.**
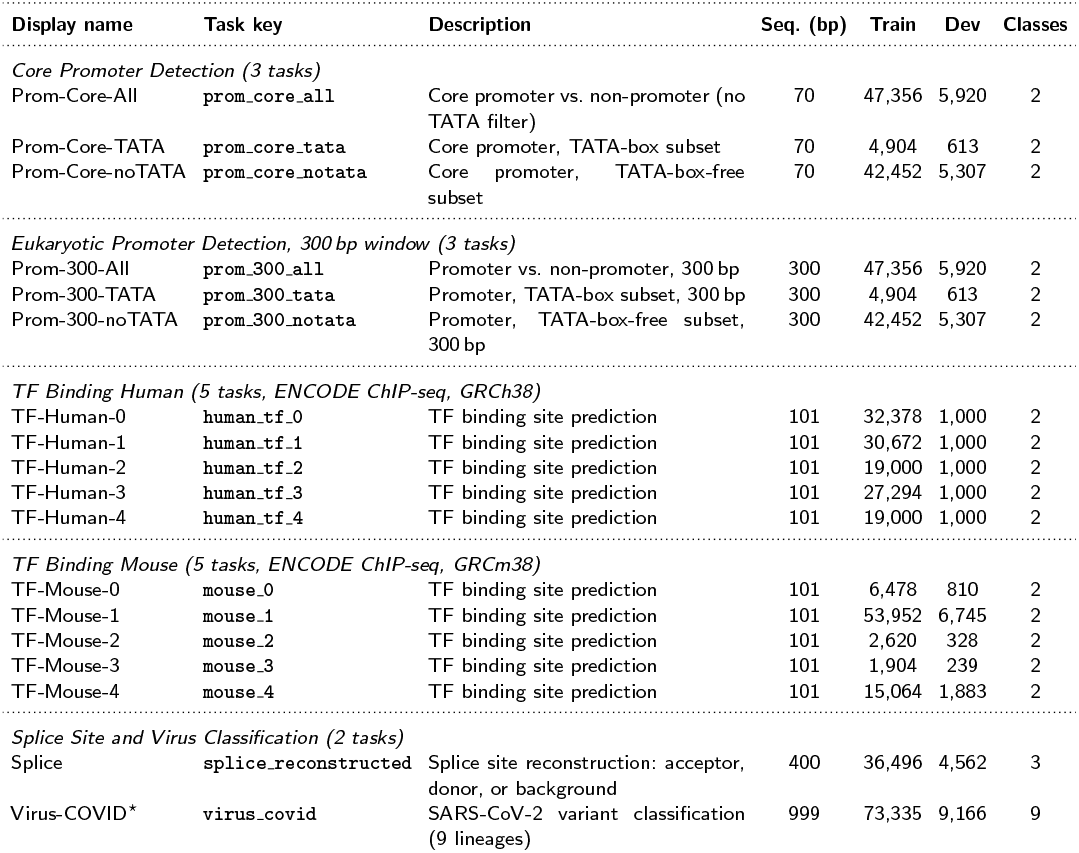
GUE task details. *Prom-Core*: mammalian core promoter window. *Prom-300* : eukaryotic promoter, extended 300 bp window. *TF-Human* and *TF-Mouse*: transcription factor binding site prediction across 5 independent cell-line / condition splits each. The TATA / noTATA splits test performance on sequences with or without a TATA box motif. ^*⋆*^ Virus-COVID uses macro-F1 as the primary metric; all other tasks use MCC.

**Table 13.** GUE+ Enhancer-Promoter Interaction (EPI) task details. Each input concatenates a local enhancer window and a local promoter window into a 5000 bp sequence with a binary label (interacting vs. non-interacting); we junction-crop it to 4096 bp for DNAGPT, while for HyenaDNA the full 5000 bp is retained within an 8192 bp budget. The released files provide only train and dev splits (the test split is not publicly available), so we evaluate on dev. The release does not include genomic coordinates or pair distances, so enhancer–promoter genomic distances are not recoverable. Split statistics reflect the released `GUE_v2.zip` files.

| Display name | Task key | Cell line | Seq. (bp) | Train | Dev | Classes |
| --- | --- | --- | --- | --- | --- | --- |
| EPI-GM12878 | `epi.0` | GM12878 (B-lymphoblastoid) | 5000 | 10,000 | 2,000 | 2 |
| EPI-HeLa-S3 | `epi.1` | HeLa-S3 (cervical carcinoma) | 5000 | 10,000 | 2,000 | 2 |
| EPI-HUVEC | `epi.2` | HUVEC (umbilical vein endothelial) | 5000 | 10,000 | 2,000 | 2 |
| EPI-IMR90 | `epi.3` | IMR90 (lung fibroblast) | 5000 | 10,000 | 2,000 | 2 |
| EPI-K562 | `epi.4` | K562 (chronic myelogenous leukaemia) | 5000 | 10,000 | 2,000 | 2 |
| EPI-NHEK | `epi.5` | NHEK (normal human epidermal keratinocyte) | 5000 | 10,000 | 2,000 | 2 |

The main analysis used no motif-flank expansion, so a contact endpoint was marked as motif-covered only when its retained base-pair position overlapped the motif instance itself. For each contact map, we considered finite upper-triangular position pairs with minimum matrix-index separation 50 and selected the top 1% by positive contact score. The observed frequency of motif–motif endpoint pairs was compared with a distance-matched permutation background using 1,000 permutations per window. Per-window empirical *P* values therefore have a minimum resolution of 1*/*(1000 + 1), and we report enrichment effect sizes and *z* statistics as the primary quantities.

Aggregate model-level summaries were computed across the 24 windows. For each model, window-level *z* statistics were combined with Stouffer’s method, and empirical *P* values were also combined with Fisher’s method as a secondary summary. Group-level summaries repeat the same procedure after restricting endpoint coverage to motif groups such as promoter-associated motifs and TFBS motifs. The cross-architecture figure uses matched recipe labels M0–M6 and H0–H6, where H0–H6 follow the same pretrain, bidirectional, masked-adaptation, dropout, reverse-complement, crop, and local-shift recipe order as M0–M6.

## Benchmark Dataset Details

This appendix provides detailed statistics for all benchmark tasks used in this study. Dataset sizes are taken from the respective original papers and data repositories and may differ slightly from those obtained by later package versions.

### Genomics Benchmarks (GB)

The Genomics Benchmarks suite (Grešová et al., 2023) provides sequence classification tasks spanning regulatory element prediction in human and mouse genomes. Data are obtained via the `genomic-benchmarks` Python package (version 0 splits). The primary evaluation metric is accuracy.

### Nucleotide Transformer Downstream Tasks (NT)

The NT downstream benchmark (Dalla-Torre et al., 2025) consists of 18 tasks: 10 histone/chromatin-mark tasks, 3 promoter tasks, 2 enhancer tasks, and 3 splice-site tasks, sourced from ENCODE ChIP-seq data, eukaryotic promoter/enhancer annotations, and the Spliceator dataset. Data are downloaded from Hugging Face (`InstaDeepAI/nucleotide_transformer_downstream_tasks`). The primary evaluation metric is MCC.

### Genome Understanding Evaluation (GUE)

From the GUE benchmark (Zhou et al., 2024), we use 18 sequence classification tasks (excluding the EMP yeast tasks), covering core promoter detection, eukaryotic promoter detection, transcription factor (TF) binding site prediction, splice site reconstruction, and SARS-CoV-2 variant classification. Data are downloaded from `leannmlindsey/GUE` on Hugging Face. Primary metrics are MCC and macro-F1 (COVID task). Models are trained on the `train` split and evaluated on the `dev` split, following the original DNABERT-2 protocol.

### GUE+ Enhancer-Promoter Interaction (EPI)

The GUE+ EPI benchmark (Zhou et al., 2024) provides six cell-line-specific binary classification tasks predicting whether a distal enhancer interacts with a target promoter. Data originate from 3C/Hi-C chromatin conformation capture experiments and are sourced from the DNABERT-2 GitHub repository (MAGICS-*L*AB/DNABERT_2). Each model input is a sequence-level simplification of the underlying chromatin-contact pair: a local enhancer window and a local promoter window are concatenated into a 5000 bp sequence. For DNAGPT we junction-crop this to 4096 bp (to fit its 1024-token BPE context window); for HyenaDNA the full 5000 bp sequence is retained within an 8192 bp budget. Although the positive pairs originate from 3C/Hi-C-identified interactions that may span much larger genomic distances, the released benchmark files contain only paired sequences and binary labels, not genomic coordinates or pair distances. We therefore inherit the dataset-level simplification and cannot recover or report the underlying enhancer–promoter genomic distance distribution from this release. The primary evaluation metric is MCC. In the released `GUE_v2.zip` files used in this study, each EPI cell-line task contains 10,000 training examples and a 2,000-example development split, which we use for evaluation (the test split is not publicly released).

## References

P. BehnamGhader, V. Adlakha, M. Mosbach, D. Bahdanau, N. Chapados, and S. Reddy. LLM2Vec: Large language models are secretly powerful text encoders. arXiv preprint 2404.05961, 2024. URL https://arxiv.org/abs/2404.05961.

H. Dalla-Torre, L. Gonzalez, J. Mendoza-Revilla, N. Lopez Carranza, A. H. Grzywaczewski, F. Oteri, C. Dallago, E. Trop, B. P. de Almeida, H. Sirelkhatim, G. Richard, M. Skwark, K. Beguir, M. Lopez, and T. Pierrot. Nucleotide transformer: Building and evaluating robust foundation models for human genomics. Nature Methods, 2025. URL https://www.nature.com/articles/s41592-024-02523-z.

J. Devlin, M.-W. Chang, K. Lee, and K. Toutanova. BERT: Pre-training of deep bidirectional transformers for language understanding. In Proceedings of NAACL, 2019.

H. Feng, L. Wu, B. Zhao, C. Huff, J. Zhang, J. Wu, L. Lin, P. Wei, and C. Wu. Benchmarking DNA foundation models for genomic and genetic tasks. Nature Communications, 16:10780, 2025. doi: 10.1038/s41467-025-65823-8. URL https://www.nature.com/articles/s41467-025-65823-8.

E. E. M. Furlong and M. Levine. Developmental enhancers and chromosome topology. Science, 361(6409):1341–1345, 2018. doi: 10.1126/science.aau0320.

T. Gao, X. Yao, and D. Chen. SimCSE: Simple contrastive learning of sentence embeddings. In Proceedings of the 2021 Conference on Empirical Methods in Natural Language Processing, pages 6894–6910, 2021.

K. Grešová, V. Martinek, D. Čechák, P. Šimeček, and P. Alexiou. Genomic benchmarks: a collection of datasets for genomic sequence classification. BMC Genomic Data, 24:25, 2023. URL https://bmcgenomdata.biomedcentral.com/articles/10.1186/s12863-023-01123-8.

E. J. Hu, Y. Shen, P. Wallis, Z. Allen-Zhu, Y. Li, S. Wang, L. Wang, and W. Chen. LoRA: Low-rank adaptation of large language models. In International Conference on Learning Representations, 2022. URL https://arxiv.org/abs/2106.09685.

InstaDeep Ltd. The genomics long-range benchmark: Advancing DNA language models. Hugging Face Datasets, 2024. https://huggingface.co/datasets/InstaDeepAI/genomics-long-range-benchmark.

Y. Ji, Z. Zhou, H. Liu, and R. V. Davuluri. DNABERT: pretrained bidirectional encoder representations from transformers model for DNA-language in genome. Bioinformatics, 37(15):2112–2120, 2021. URL https://academic.oup.com/bioinformatics/article/37/15/2112/6128680.

T. King, S. Butcher, and L. Zalewski. Apocrita – high performance computing cluster for queen mary university of london. Queen Mary University of London, 2017. URL 10.5281/zenodo.438045.

V. Mallet and J.-P. Vert. Reverse-complement equivariant networks for DNA sequences. In Advances in Neural Information Processing Systems, volume 34, 2021. URL https://proceedings.neurips.cc/paper/2021/hash/706608cfdbcc1886bb7eea5513f90133-Abstract.html.

M. T. Maurano, R. Humbert, E. Rynes, R. E. Thurman, E. Haugen, H. Wang, A. P. Reynolds, R. Sandstrom, H. Qu, J. Brody, et al. Systematic localization of common disease-associated variation in regulatory DNA. Science, 337(6099): 1190–1195, 2012. doi: 10.1126/science.1222794.

E. Nguyen, M. Poli, M. Faizi, A. Thomas, C. Birch-Sykes, M. Wornow, A. Patel, C. Rabideau, S. Massaroli, Y. Bengio, S. Ermon, S. A. Baccus, and C. Ré. HyenaDNA: Long-range genomic sequence modeling at single nucleotide resolution. In Advances in Neural Information Processing Systems, 2023. URL https://pmc.ncbi.nlm.nih.gov/articles/PMC10327243/.

F. Pedregosa, G. Varoquaux, A. Gramfort, V. Michel, B. Thirion, O. Grisel, M. Blondel, A. Müller, J. Nothman, G. Louppe, P. Prettenhofer, R. Weiss, V. Dubourg, J. Vanderplas, A. Passos, D. Cournapeau, M. Brucher, M. Perrot, and É. Duchesnay. Scikit-learn: Machine learning in Python. Journal of Machine Learning Research, 12:2825–2830, 2011.

A. Radford, J. Wu, R. Child, D. Luan, D. Amodei, and I. Sutskever. Language models are unsupervised multitask learners. 2019. URL https://cdn.openai.com/better-language-models/language_models_are_unsupervised_multitask_learners.pdf.

M. Safari, P. Millan Arias, S. C. Lowe, L. Kari, A. X. Chang, and G. W. Taylor. Enhancing DNA foundation models to address masking inefficiencies. arXiv preprint 2502.18405, 2025. URL https://arxiv.org/abs/2502.18405.

Y. Schiff, C.-H. Kao, A. Gokaslan, T. Dao, A. Gu, and V. Kuleshov. Caduceus: Bi-directional equivariant long-range DNA sequence modeling. In International Conference on Machine Learning, 2024. URL https://arxiv.org/abs/2403.03234.

V. A. Schneider, T. Graves-Lindsay, K. Howe, N. Bouk, H.-C. Chen, P. A. Kitts, T. D. Murphy, K. D. Pruitt, F. Thibaud-Nissen, D. Albracht, R. S. Fulton, M. Kremitzki, V. Magrini, C. Markovic, S. McGrath, K. M. Steinberg, K. Auger, W. Chow, J. Collins, G. Harden, T. Hubbard, S. Pelan, J. T. Simpson, G. Threadgold, J. Torrance, J. M. Wood, L. Clarke, S. Koren, M. Boitano, P. Peluso, H. Li, C.-S. Chin, A. M. Phillippy, R. Durbin, R. K. Wilson, P. Flicek, E. E. Eichler, and D. M. Church. Evaluation of GRCh38 and de novo haploid genome assemblies demonstrates the enduring quality of the reference assembly. Genome Research, 27(5):849–864, 2017. doi: 10.1101/gr.213611.116.

S. Schoenfelder and P. Fraser. Long-range enhancer-promoter contacts in gene expression control. Nature Reviews Genetics, 20:437–455, 2019. doi: 10.1038/s41576-019-0128-0.

L. Shu J. Tang X. Guan, and D. Zhang. A comprehensive survey of genome language models in bioinformatics. Briefings in Bioinformatics, 27(1):bbaf724, 2026. doi: 10.1093/bib/bbaf724. URL https://academic.oup.com/bib/article/27/1/bbaf724/8426124.

D. Zhang, W. Zhang, Y. Zhao, J. Zhang, B. He, C. Qin, and J. Yao. DNAGPT: A generalized pre-trained tool for versatile DNA sequence analysis tasks. arXiv preprint 2307.05628, 2023. URL https://arxiv.org/abs/2307.05628.

H. Zhou, A. Shrikumar, and A. Kundaje. Towards a better understanding of reverse-complement equivariance for deep learning models in genomics. In Proceedings of the 16th Machine Learning in Computational Biology Meeting, volume 165 of Proceedings of Machine Learning Research, pages 1–33, 2022. URL https://proceedings.mlr.press/v165/zhou22a.html.

Z. Zhou, Y. Ji, W. Li, P. Dutta, R. Davuluri, and H. Liu. DNABERT-2: Efficient foundation model and benchmark for multi-species genome. In International Conference on Learning Representations, 2024. URL https://arxiv.org/abs/2306.15006.

