## Supplementary Material for "DNA-LLM2Vec: Architecture-Aware Task-Conditioned Adaptation of Decoder DNA Foundation Models"

October 7, 2026

#### S1 Step 4 Linear Probe Full Per-Seed Results

Tables S1–S6 report DNAGPT step 4 linear probe results for all model variants and all benchmark suites at the individual seed level, under the *mask-only* embedding-table update policy used in the main paper; the full-embedding comparison is included below. M0 and M1 involve no stochastic training and are therefore deterministic (labelled "det."). M2–M6 are each run with three independent random seeds (s42, s43, s44); shaded rows give the mean ± std across the three seeds.

##### S1.1 DNAGPT Step 4 Linear Probe Per-Seed Results

##### S1.2 HyenaDNA Step 4 Linear Probe Per-Seed Results

Tables S8–S14 report the corresponding HyenaDNA Step 4 linear-probe results used for the cross-architecture task-family analyses. H0 and H1 are deterministic in this evaluation; H2–H6 report three independent adaptation seeds and mean ± std rows.

#### S2 Step 5 Full Fine-Tuning Per-Seed Results

Tables S15–S22 report the Step 5 full fine-tuning results at the individual-seed level. DNAGPT adapted rows use the mask-only policy matching the main-paper full fine-

Table S1: Genomics Benchmarks (GB) per-seed linear probe accuracy (%), *mask-only* setting, expanded to all 8 GB tasks.

| **Model** | **Run** | **Coding-vs-Intergenic** | **Human-vs-Worm** | **Enh-Cohn** | **Enh-Ensembl** | **Regulatory** | **Non-TATA** | **OCR** | **Mouse-Enh** | **Avg-GB** |
|---|---|---|---|---|---|---|---|---|---|---|
| M0 (DNAGPT causal) | (det.) | 89.54±0.00 | 93.87±0.00 | 73.65±0.00 | 74.06±0.00 | 63.70±0.00 | 82.77±0.00 | 68.10±0.00 | 78.93±0.00 | 78.08±0.00 |
| M1 (+bidir attn) | (det.) | 89.52±0.00 | 94.12±0.00 | 73.20±0.00 | 74.03±0.00 | 69.51±0.00 | 83.05±0.00 | 67.26±0.00 | 76.03±0.00 | 78.34±0.00 |
| M2 (+MNTP) | s42 | 89.48 | 93.86 | 74.04 | 73.77 | 66.14 | 83.22 | 67.67 | 77.69 | 78.23 |
| M2 (+MNTP) | s43 | 89.44 | 93.83 | 73.70 | 73.74 | 66.20 | 83.35 | 67.40 | 77.69 | 78.17 |
| M2 (+MNTP) | s44 | 89.33 | 93.74 | 73.69 | 73.76 | 66.69 | 83.38 | 67.89 | 78.51 | 78.38 |
| M2 (+MNTP) | mean±std | 89.42±0.08 | 93.81±0.06 | 73.81±0.20 | 73.76±0.02 | 66.34±0.30 | 83.32±0.09 | 67.65±0.25 | 77.96±0.47 | 78.26±0.11 |
| M3 (+dropout SimCSE) | s42 | 89.45 | 94.22 | 73.62 | 74.10 | 66.82 | 83.25 | 67.80 | 78.10 | 78.42 |
| M3 (+dropout SimCSE) | s43 | 89.67 | 94.17 | 73.98 | 73.86 | 67.10 | 83.26 | 67.29 | 78.10 | 78.43 |
| M3 (+dropout SimCSE) | s44 | 89.40 | 93.94 | 73.86 | 73.99 | 67.46 | 83.51 | 67.92 | 77.69 | 78.47 |
| M3 (+dropout SimCSE) | mean±std | 89.51±0.14 | 94.11±0.15 | 73.82±0.18 | 73.98±0.12 | 67.13±0.32 | 83.34±0.15 | 67.67±0.33 | 77.96±0.24 | 78.44±0.03 |
| M4 (+RC SimCSE) | s42 | 89.69 | 94.25 | 73.46 | 74.38 | 67.44 | 82.67 | 67.78 | 81.82 | 78.94 |
| M4 (+RC SimCSE) | s43 | 89.85 | 94.48 | 73.59 | 74.18 | 67.31 | 82.93 | 67.82 | 80.58 | 78.84 |
| M4 (+RC SimCSE) | s44 | 89.73 | 94.26 | 73.68 | 74.21 | 67.15 | 83.08 | 68.06 | 82.64 | 79.10 |
| M4 (+RC SimCSE) | mean±std | 89.76±0.08 | 94.33±0.13 | 73.58±0.11 | 74.26±0.11 | 67.30±0.15 | 82.89±0.21 | 67.89±0.15 | 81.68±1.04 | 78.96±0.13 |
| M5 (+crop SimCSE) | s42 | 88.81 | 93.64 | 72.90 | 73.43 | 66.73 | 82.82 | 66.99 | 78.93 | 78.03 |
| M5 (+crop SimCSE) | s43 | 89.04 | 93.82 | 73.03 | 73.87 | 66.78 | 83.22 | 67.17 | 78.93 | 78.23 |
| M5 (+crop SimCSE) | s44 | 88.62 | 93.48 | 72.64 | 73.15 | 67.32 | 82.99 | 67.09 | 79.75 | 78.13 |
| M5 (+crop SimCSE) | mean±std | 88.82±0.21 | 93.65±0.17 | 72.86±0.20 | 73.48±0.36 | 66.94±0.33 | 83.01±0.20 | 67.08±0.09 | 79.20±0.47 | 78.13±0.10 |
| M6 (+shift SimCSE) | s42 | 89.11 | 93.78 | 73.65 | 74.05 | 67.66 | 82.90 | 67.44 | 78.93 | 78.44 |
| M6 (+shift SimCSE) | s43 | 89.05 | 93.72 | 73.82 | 73.82 | 67.88 | 83.16 | 67.42 | 80.17 | 78.63 |
| M6 (+shift SimCSE) | s44 | 88.90 | 93.69 | 73.20 | 73.67 | 67.57 | 82.99 | 67.80 | 80.58 | 78.55 |
| M6 (+shift SimCSE) | mean±std | 89.02±0.11 | 93.73±0.05 | 73.56±0.32 | 73.85±0.19 | 67.70±0.16 | 83.02±0.13 | 67.55±0.21 | 79.89±0.86 | 78.54±0.10 |

Table S2: Nucleotide Transformer downstream tasks (NT) per-seed linear probe MCC (%), *mask-only* setting, part A: promoter, enhancer, splice, and aggregate tasks.

| **Model** | **Run** | **Prom-All** | **Prom-TATA** | **Prom-noTA** | **Enhancers** | **Enh-Types** | **Splice-All** | **Splice-Acc** | **Splice-Donor** | **Avg-NT** |
|---|---|---|---|---|---|---|---|---|---|---|
| M0 (DNAGPT causal) | (det.) | 84.39±0.00 | 80.96±0.00 | 84.04±0.00 | 48.02±0.00 | 33.99±0.00 | 31.92±0.00 | 35.45±0.00 | 34.02±0.00 | 45.99±0.00 |
| M1 (+bidir attn) | (det.) | 84.09±0.00 | 80.46±0.00 | 84.01±0.00 | 44.00±0.00 | 32.45±0.00 | 25.22±0.00 | 37.39±0.00 | 27.74±0.00 | 44.50±0.00 |
| M2 (+MNTP) | s42 | 82.46 | 81.65 | 82.20 | 48.52 | 32.97 | 28.07 | 37.04 | 31.55 | 45.81 |
| M2 (+MNTP) | s43 | 82.63 | 82.90 | 82.50 | 44.01 | 32.64 | 28.43 | 39.31 | 30.77 | 45.54 |
| M2 (+MNTP) | s44 | 82.33 | 81.57 | 82.58 | 47.51 | 30.48 | 28.28 | 38.96 | 33.40 | 45.92 |
| M2 (+MNTP) | mean±std | 82.47±0.15 | 82.04±0.75 | 82.43±0.20 | 46.68±2.37 | 32.03±1.35 | 28.26±0.18 | 38.44±1.22 | 31.91±1.35 | 45.76±0.20 |
| M3 (+dropout SimCSE) | s42 | 84.11 | 82.92 | 84.20 | 46.00 | 30.79 | 28.45 | 38.45 | 29.99 | 46.15 |
| M3 (+dropout SimCSE) | s43 | 84.00 | 85.56 | 83.70 | 45.01 | 31.81 | 29.46 | 39.55 | 29.29 | 46.08 |
| M3 (+dropout SimCSE) | s44 | 84.85 | 85.17 | 84.50 | 43.03 | 32.39 | 29.30 | 37.54 | 32.59 | 46.63 |
| M3 (+dropout SimCSE) | mean±std | 84.32±0.46 | 84.55±1.43 | 84.13±0.40 | 44.68±1.51 | 31.66±0.81 | 29.07±0.54 | 38.51±1.01 | 30.62±1.74 | 46.29±0.30 |
| M4 (+RC SimCSE) | s42 | 84.59 | 83.62 | 84.13 | 46.06 | 32.04 | 24.28 | 38.36 | 33.85 | 46.82 |
| M4 (+RC SimCSE) | s43 | 85.56 | 83.96 | 85.41 | 48.09 | 33.69 | 25.96 | 38.82 | 33.24 | 47.41 |
| M4 (+RC SimCSE) | s44 | 84.45 | 85.96 | 84.89 | 46.63 | 28.99 | 22.70 | 40.34 | 34.34 | 47.11 |
| M4 (+RC SimCSE) | mean±std | 84.87±0.60 | 84.51±1.26 | 84.81±0.64 | 46.93±1.05 | 31.57±2.38 | 24.31±1.63 | 39.17±1.04 | 33.81±0.55 | 47.11±0.30 |
| M5 (+crop SimCSE) | s42 | 83.46 | 82.36 | 83.33 | 46.06 | 31.48 | 23.26 | 38.42 | 31.36 | 45.32 |
| M5 (+crop SimCSE) | s43 | 84.66 | 84.98 | 84.25 | 45.67 | 33.04 | 24.11 | 39.31 | 32.63 | 46.31 |
| M5 (+crop SimCSE) | s44 | 84.11 | 83.04 | 83.08 | 43.18 | 30.43 | 23.16 | 39.76 | 34.30 | 45.60 |
| M5 (+crop SimCSE) | mean±std | 84.08±0.60 | 83.46±1.36 | 83.55±0.62 | 44.97±1.56 | 31.65±1.31 | 23.51±0.52 | 39.16±0.68 | 32.76±1.47 | 45.74±0.51 |
| M6 (+shift SimCSE) | s42 | 84.79 | 81.96 | 84.91 | 45.51 | 29.82 | 24.56 | 38.22 | 31.51 | 45.75 |
| M6 (+shift SimCSE) | s43 | 84.30 | 83.99 | 84.36 | 43.00 | 31.07 | 26.42 | 39.93 | 32.04 | 46.16 |
| M6 (+shift SimCSE) | s44 | 84.27 | 83.96 | 83.90 | 45.11 | 31.11 | 25.87 | 39.04 | 35.96 | 46.53 |
| M6 (+shift SimCSE) | mean±std | 84.45±0.29 | 83.30±1.16 | 84.39±0.51 | 44.54±1.35 | 30.67±0.73 | 25.62±0.96 | 39.06±0.86 | 33.17±2.43 | 46.15±0.39 |

Table S3: Nucleotide Transformer downstream tasks (NT) per-seed linear probe MCC (%), *mask-only* setting, part B: histone-mark tasks and aggregate.

| **Model** | **Run** | **H3** | **H4** | **H3K9ac** | **H3K14ac** | **H4ac** | **H3K4me1** | **H3K4me2** | **H3K4me3** | **H3K36me3** | **H3K79me3** | **Avg-NT** |
|---|---|---|---|---|---|---|---|---|---|---|---|---|
| M0 (DNAGPT causal) | (det.) | 64.58±0.00 | 69.43±0.00 | 38.43±0.00 | 30.62±0.00 | 27.74±0.00 | 31.69±0.00 | 25.49±0.00 | 19.89±0.00 | 37.47±0.00 | 49.64±0.00 | 45.99±0.00 |
| M1 (+bidir attn) | (det.) | 63.64±0.00 | 66.20±0.00 | 36.51±0.00 | 28.27±0.00 | 26.86±0.00 | 29.15±0.00 | 25.22±0.00 | 18.72±0.00 | 39.34±0.00 | 51.74±0.00 | 44.50±0.00 |
| M2 (+MNTP) | s42 | 64.57 | 72.26 | 38.32 | 29.12 | 28.70 | 29.96 | 25.83 | 21.53 | 38.43 | 51.32 | 45.81 |
| M2 (+MNTP) | s43 | 63.23 | 69.76 | 38.05 | 28.96 | 28.23 | 28.53 | 27.43 | 20.79 | 38.31 | 53.20 | 45.54 |
| M2 (+MNTP) | s44 | 64.59 | 72.04 | 38.07 | 30.58 | 28.48 | 29.19 | 26.95 | 19.51 | 39.47 | 52.49 | 45.92 |
| M2 (+MNTP) | mean±std | 64.13±0.78 | 71.35±1.38 | 38.15±0.15 | 29.55±0.89 | 28.47±0.24 | 29.23±0.72 | 26.74±0.82 | 20.61±1.02 | 38.74±0.64 | 52.34±0.95 | 45.76±0.20 |
| M3 (+dropout SimCSE) | s42 | 65.37 | 71.45 | 38.82 | 31.62 | 29.01 | 30.47 | 26.41 | 20.56 | 38.49 | 53.55 | 46.15 |
| M3 (+dropout SimCSE) | s43 | 64.17 | 71.58 | 37.88 | 30.21 | 28.25 | 31.33 | 25.90 | 20.18 | 38.14 | 53.48 | 46.08 |
| M3 (+dropout SimCSE) | s44 | 64.70 | 73.38 | 38.51 | 32.21 | 29.12 | 30.80 | 27.36 | 20.24 | 38.91 | 54.73 | 46.63 |
| M3 (+dropout SimCSE) | mean±std | 64.75±0.60 | 72.14±1.08 | 38.40±0.48 | 31.35±1.03 | 28.79±0.47 | 30.87±0.43 | 26.56±0.74 | 20.33±0.20 | 38.51±0.39 | 53.92±0.70 | 46.29±0.30 |
| M4 (+RC SimCSE) | s42 | 67.12 | 71.69 | 40.16 | 31.55 | 30.59 | 31.09 | 27.78 | 22.38 | 38.93 | 54.53 | 46.82 |
| M4 (+RC SimCSE) | s43 | 66.59 | 72.25 | 38.66 | 32.23 | 30.77 | 32.55 | 28.27 | 23.06 | 39.53 | 54.81 | 47.41 |
| M4 (+RC SimCSE) | s44 | 66.84 | 72.14 | 39.55 | 33.24 | 30.00 | 31.92 | 26.93 | 22.85 | 40.47 | 55.71 | 47.11 |
| M4 (+RC SimCSE) | mean±std | 66.85±0.27 | 72.03±0.30 | 39.46±0.75 | 32.34±0.85 | 30.45±0.40 | 31.85±0.73 | 27.66±0.68 | 22.76±0.35 | 39.64±0.78 | 55.02±0.62 | 47.11±0.30 |
| M5 (+crop SimCSE) | s42 | 64.17 | 66.11 | 37.83 | 30.75 | 28.50 | 30.30 | 26.22 | 22.29 | 38.65 | 51.18 | 45.32 |
| M5 (+crop SimCSE) | s43 | 65.37 | 66.97 | 39.15 | 31.95 | 28.87 | 31.98 | 28.16 | 22.32 | 39.60 | 50.61 | 46.31 |
| M5 (+crop SimCSE) | s44 | 63.63 | 67.38 | 38.48 | 31.99 | 28.89 | 30.91 | 26.14 | 22.35 | 39.73 | 50.25 | 45.60 |
| M5 (+crop SimCSE) | mean±std | 64.39±0.89 | 66.82±0.65 | 38.49±0.66 | 31.56±0.70 | 28.75±0.22 | 31.06±0.85 | 26.84±1.14 | 22.32±0.03 | 39.33±0.59 | 50.68±0.47 | 45.74±0.51 |
| M6 (+shift SimCSE) | s42 | 66.46 | 69.51 | 38.55 | 30.91 | 29.36 | 29.42 | 25.81 | 21.37 | 38.74 | 52.14 | 45.75 |
| M6 (+shift SimCSE) | s43 | 65.78 | 70.45 | 38.16 | 31.43 | 28.49 | 29.57 | 27.08 | 20.98 | 39.96 | 53.89 | 46.16 |
| M6 (+shift SimCSE) | s44 | 66.73 | 71.02 | 37.72 | 33.19 | 29.47 | 30.04 | 27.38 | 21.55 | 38.82 | 52.42 | 46.53 |
| M6 (+shift SimCSE) | mean±std | 66.32±0.49 | 70.33±0.76 | 38.14±0.42 | 31.84±1.19 | 29.11±0.54 | 29.68±0.32 | 26.76±0.83 | 21.30±0.29 | 39.17±0.68 | 52.82±0.94 | 46.15±0.39 |

Table S4: GUE+ EPI per-seed linear probe MCC (%), *mask-only* setting. Six cell-line tasks plus average shown.

| **Model** | **Run** | **GM12878** | **HeLa-S3** | **HUVEC** | **IMR90** | **K562** | **NHEK** | **Avg-EPI** |
|---|---|---|---|---|---|---|---|---|
| M0 (DNAGPT causal) | (det.) | 13.84 | 11.65 | 17.85 | 15.10 | 13.01 | 9.07 | 13.42 |
| M1 (+bidir attn) | (det.) | 12.67 | 10.64 | 18.42 | 17.19 | 11.70 | 13.11 | 13.96 |
| M2 (+MNTP) | s42 | 16.74 | 18.46 | 23.94 | 22.80 | 16.39 | 17.20 | 19.26 |
| M2 (+MNTP) | s43 | 16.90 | 18.14 | 25.26 | 22.70 | 17.75 | 16.21 | 19.49 |
| M2 (+MNTP) | s44 | 17.12 | 18.79 | 25.07 | 23.76 | 18.44 | 18.90 | 20.35 |
| M2 (+MNTP) | mean±std | 16.92±0.19 | 18.46±0.33 | 24.76±0.71 | 23.09±0.59 | 17.53±1.04 | 17.44±1.36 | 19.70±0.57 |
| M3 (+dropout SimCSE) | s42 | 21.30 | 19.55 | 26.39 | 27.35 | 20.19 | 23.93 | 23.12 |
| M3 (+dropout SimCSE) | s43 | 20.72 | 21.14 | 27.52 | 28.77 | 20.15 | 22.75 | 23.51 |
| M3 (+dropout SimCSE) | s44 | 18.17 | 19.33 | 27.31 | 31.30 | 20.05 | 21.98 | 23.02 |
| M3 (+dropout SimCSE) | mean±std | 20.06±1.67 | 20.01±0.99 | 27.07±0.60 | 29.14±2.00 | 20.13±0.07 | 22.89±0.98 | 23.22±0.26 |
| M4 (+RC SimCSE) | s42 | 18.75 | 17.74 | 27.90 | 25.48 | 16.72 | 19.85 | 21.07 |
| M4 (+RC SimCSE) | s43 | 19.80 | 17.85 | 27.39 | 25.90 | 17.19 | 18.33 | 21.08 |
| M4 (+RC SimCSE) | s44 | 17.83 | 19.45 | 26.87 | 25.84 | 16.48 | 18.74 | 20.87 |
| M4 (+RC SimCSE) | mean±std | 18.79±0.99 | 18.35±0.96 | 27.39±0.52 | 25.74±0.23 | 16.80±0.36 | 18.97±0.79 | 21.01±0.12 |
| M5 (+crop SimCSE) | s42 | 21.38 | 25.19 | 30.58 | 32.71 | 23.97 | 31.86 | 27.61 |
| M5 (+crop SimCSE) | s43 | 20.98 | 24.49 | 29.89 | 33.72 | 25.08 | 31.06 | 27.54 |
| M5 (+crop SimCSE) | s44 | 22.45 | 24.27 | 28.05 | 34.32 | 22.24 | 32.06 | 27.23 |
| M5 (+crop SimCSE) | mean±std | 21.60±0.76 | 24.65±0.48 | 29.51±1.31 | 33.58±0.81 | 23.76±1.43 | 31.66±0.53 | 27.46±0.20 |
| M6 (+shift SimCSE) | s42 | 19.44 | 19.75 | 26.97 | 27.16 | 20.08 | 23.38 | 22.80 |
| M6 (+shift SimCSE) | s43 | 20.61 | 21.28 | 28.67 | 30.08 | 22.01 | 25.02 | 24.61 |
| M6 (+shift SimCSE) | s44 | 19.44 | 21.66 | 30.08 | 32.18 | 22.52 | 27.80 | 25.61 |
| M6 (+shift SimCSE) | mean±std | 19.83±0.68 | 20.90±1.01 | 28.57±1.56 | 29.81±2.52 | 21.54±1.29 | 25.40±2.23 | 24.34±1.42 |

Table S5: GUE benchmark per-seed MCC (%), *mask-only* setting, part A: core promoter (PC), promoter-300 (P3), and TF binding in human.

| **Model** | **Run** | **PC-All** | **PC-TATA** | **PC-noTA** | **P3-All** | **P3-TATA** | **P3-noTA** | **TF-H0** | **TF-H1** | **TF-H2** | **TF-H3** | **TF-H4** |
|---|---|---|---|---|---|---|---|---|---|---|---|---|
| M0 (DNAGPT causal) | (det.) | 54.17 | 47.18 | 61.10 | 78.55 | 44.73 | 85.93 | 50.35 | 49.25 | 49.05 | 41.28 | 52.96 |
| M1 (+bidir attn) | (det.) | 57.88 | 43.46 | 62.37 | 76.97 | 35.07 | 85.33 | 55.33 | 52.32 | 47.84 | 43.68 | 52.23 |
| M2 (+MNTP) | s42 | 56.45 | 42.33 | 62.86 | 76.59 | 40.00 | 84.12 | 53.94 | 51.69 | 49.72 | 45.49 | 53.33 |
| M2 (+MNTP) | s43 | 56.69 | 42.26 | 63.24 | 76.56 | 37.04 | 84.11 | 53.37 | 51.07 | 48.48 | 43.28 | 54.49 |
| M2 (+MNTP) | s44 | 56.05 | 42.36 | 63.58 | 75.71 | 39.99 | 83.79 | 53.98 | 51.02 | 46.92 | 40.29 | 52.90 |
| M2 (+MNTP) | mean±std | 56.40±0.32 | 42.32±0.05 | 63.23±0.36 | 76.29±0.50 | 39.01±1.71 | 84.01±0.19 | 53.76±0.34 | 51.26±0.37 | 48.37±1.40 | 43.02±2.61 | 53.57±0.82 |
| M3 (+dropout SimCSE) | s42 | 55.10 | 48.57 | 63.84 | 77.60 | 39.70 | 85.32 | 53.79 | 52.43 | 47.47 | 43.08 | 53.51 |
| M3 (+dropout SimCSE) | s43 | 56.32 | 45.93 | 63.16 | 77.10 | 38.78 | 84.79 | 53.77 | 51.24 | 45.91 | 45.49 | 52.50 |
| M3 (+dropout SimCSE) | s44 | 56.32 | 44.40 | 64.41 | 77.77 | 38.73 | 85.03 | 55.38 | 52.22 | 46.91 | 45.29 | 53.88 |
| M3 (+dropout SimCSE) | mean±std | 55.91±0.70 | 46.30±2.11 | 63.80±0.63 | 77.49±0.35 | 39.07±0.55 | 85.05±0.27 | 54.31±0.92 | 51.96±0.64 | 46.76±0.79 | 44.62±1.34 | 53.30±0.71 |
| M4 (+RC SimCSE) | s42 | 55.57 | 50.09 | 64.15 | 77.91 | 38.53 | 84.66 | 52.88 | 53.24 | 50.45 | 41.27 | 52.77 |
| M4 (+RC SimCSE) | s43 | 56.89 | 43.87 | 63.47 | 78.24 | 41.15 | 85.64 | 54.81 | 51.01 | 49.24 | 39.66 | 51.29 |
| M4 (+RC SimCSE) | s44 | 55.47 | 48.09 | 63.92 | 77.63 | 39.52 | 85.31 | 53.18 | 51.40 | 46.46 | 40.47 | 51.44 |
| M4 (+RC SimCSE) | mean±std | 55.98±0.79 | 47.35±3.18 | 63.85±0.35 | 77.93±0.31 | 39.73±1.32 | 85.20±0.50 | 53.62±1.04 | 51.88±1.19 | 48.72±2.05 | 40.47±0.81 | 51.83±0.81 |
| M5 (+crop SimCSE) | s42 | 54.70 | 47.62 | 61.45 | 76.23 | 36.41 | 84.83 | 51.93 | 51.25 | 44.39 | 42.48 | 52.53 |
| M5 (+crop SimCSE) | s43 | 54.73 | 44.59 | 62.09 | 77.47 | 40.47 | 85.64 | 51.95 | 52.04 | 44.98 | 44.29 | 52.70 |
| M5 (+crop SimCSE) | s44 | 54.40 | 48.96 | 62.31 | 76.66 | 36.48 | 84.27 | 52.12 | 50.23 | 44.78 | 43.30 | 50.87 |
| M5 (+crop SimCSE) | mean±std | 54.61±0.18 | 47.06±2.24 | 61.95±0.45 | 76.79±0.63 | 37.79±2.32 | 84.91±0.69 | 52.00±0.10 | 51.17±0.91 | 44.72±0.30 | 43.36±0.91 | 52.03±1.01 |
| M6 (+shift SimCSE) | s42 | 55.41 | 47.02 | 63.77 | 77.37 | 36.11 | 85.49 | 52.41 | 53.69 | 47.24 | 42.28 | 52.77 |
| M6 (+shift SimCSE) | s43 | 54.25 | 47.05 | 63.17 | 77.84 | 39.74 | 86.50 | 51.95 | 50.62 | 47.79 | 42.68 | 53.75 |
| M6 (+shift SimCSE) | s44 | 54.93 | 45.77 | 63.13 | 76.43 | 39.07 | 85.04 | 54.14 | 51.02 | 47.47 | 43.68 | 51.67 |
| M6 (+shift SimCSE) | mean±std | 54.86±0.58 | 46.61±0.73 | 63.36±0.36 | 77.21±0.72 | 38.31±1.93 | 85.68±0.75 | 52.83±1.15 | 51.78±1.67 | 47.50±0.28 | 42.88±0.72 | 52.73±1.04 |

Table S6: GUE benchmark per-seed MCC (%), *mask-only* setting, part B: TF binding in mouse, splice site reconstruction, SARS-CoV-2 MCC, and non-COVID GUE average.

| **Model** | **Run** | **TF-M0** | **TF-M1** | **TF-M2** | **TF-M3** | **TF-M4** | **Splice** | **COVID-MCC** | **Avg-GUE17** |
|---|---|---|---|---|---|---|---|---|---|
| M0 (DNAGPT causal) | (det.) | 35.36 | 68.47 | 60.99 | 36.43 | 27.99 | 19.79 | 20.27 | 50.80 |
| M1 (+bidir attn) | (det.) | 29.97 | 67.36 | 61.58 | 26.36 | 26.54 | 11.02 | 3.97 | 49.14 |
| M2 (+MNTP) | s42 | 27.19 | 64.88 | 62.89 | 41.55 | 26.08 | 16.03 | 23.31 | 50.30 |
| M2 (+MNTP) | s43 | 25.47 | 64.73 | 62.83 | 34.01 | 24.71 | 16.27 | 24.48 | 49.33 |
| M2 (+MNTP) | s44 | 25.59 | 65.12 | 59.81 | 38.14 | 26.96 | 17.16 | 23.81 | 49.37 |
| M2 (+MNTP) | mean±std | 26.08±0.96 | 64.91±0.20 | 61.84±1.76 | 37.90±3.78 | 25.92±1.13 | 16.49±0.60 | 23.87±0.59 | 49.67±0.55 |
| M3 (+dropout SimCSE) | s42 | 28.91 | 65.95 | 63.42 | 39.87 | 25.56 | 17.63 | 21.22 | 50.69 |
| M3 (+dropout SimCSE) | s43 | 25.18 | 65.72 | 64.02 | 47.31 | 26.26 | 16.38 | 20.96 | 50.58 |
| M3 (+dropout SimCSE) | s44 | 28.64 | 66.72 | 61.59 | 44.80 | 25.17 | 17.73 | 20.64 | 50.88 |
| M3 (+dropout SimCSE) | mean±std | 27.58±2.08 | 66.13±0.52 | 63.01±1.27 | 43.99±3.79 | 25.66±0.55 | 17.25±0.75 | 20.94±0.29 | 50.72±0.15 |
| M4 (+RC SimCSE) | s42 | 26.37 | 69.86 | 64.63 | 41.44 | 27.78 | 12.93 | 23.56 | 50.85 |
| M4 (+RC SimCSE) | s43 | 28.81 | 70.16 | 67.08 | 36.41 | 26.10 | 16.02 | 21.43 | 50.58 |
| M4 (+RC SimCSE) | s44 | 31.55 | 70.17 | 63.46 | 43.21 | 26.85 | 15.12 | 22.60 | 50.78 |
| M4 (+RC SimCSE) | mean±std | 28.91±2.59 | 70.06±0.18 | 65.06±1.85 | 40.35±3.53 | 26.91±0.84 | 14.69±1.59 | 22.53±1.07 | 50.74±0.14 |
| M5 (+crop SimCSE) | s42 | 27.88 | 65.13 | 61.58 | 46.51 | 23.95 | 15.63 | 34.58 | 49.68 |
| M5 (+crop SimCSE) | s43 | 29.11 | 65.02 | 61.61 | 43.98 | 25.82 | 17.61 | 35.95 | 50.24 |
| M5 (+crop SimCSE) | s44 | 30.71 | 65.53 | 62.21 | 41.45 | 24.05 | 16.74 | 34.77 | 49.71 |
| M5 (+crop SimCSE) | mean±std | 29.23±1.42 | 65.23±0.27 | 61.80±0.36 | 43.98±2.53 | 24.61±1.05 | 16.66±0.99 | 35.10±0.74 | 49.88±0.32 |
| M6 (+shift SimCSE) | s42 | 30.59 | 67.91 | 62.26 | 51.46 | 25.53 | 14.29 | 30.45 | 50.92 |
| M6 (+shift SimCSE) | s43 | 30.85 | 66.13 | 62.23 | 39.75 | 24.60 | 17.20 | 32.83 | 50.36 |
| M6 (+shift SimCSE) | s44 | 27.72 | 65.54 | 59.76 | 44.85 | 27.89 | 15.88 | 33.54 | 50.23 |
| M6 (+shift SimCSE) | mean±std | 29.72±1.74 | 66.53±1.23 | 61.42±1.43 | 45.35±5.87 | 26.01±1.70 | 15.79±1.46 | 32.27±1.62 | 50.50±0.36 |

Table S7: Step 4 Virus-COVID macro-F1 under the main mean-pooling linear-probe protocol. Virus-COVID uses macro-F1 as its official metric, whereas the other GUE tasks use MCC. M2–M6 report the mask-only setting used in the main Step 4 results and are mean ± std over three seeds.

| **Model** | **Virus-COVID F1** |
|---|---|
| M0 (DNAGPT causal) | 27.40 |
| M1 (+bidir attn) | 11.81 |
| M2 (+MNTP LoRA) | 32.10±0.87 |
| M3 (+dropout SimCSE) | 28.85±0.28 |
| M4 (+revcomp SimCSE) | 30.28±1.18 |
| M5 (+crop SimCSE) | 42.63±0.68 |
| M6 (+local-shift SimCSE) | 39.93±1.41 |

Table S8: HyenaDNA Genomics Benchmarks (GB) per-seed linear probe accuracy (%), expanded to all 8 GB tasks.

| **Model** | **Run** | **Coding-vs-Intergenic** | **Human-vs-Worm** | **Enh-Cohn** | **Enh-Ensembl** | **Regulatory** | **Non-TATA** | **OCR** | **Mouse-Enh** | **Avg-GB** |
|---|---|---|---|---|---|---|---|---|---|---|
| H0 (HyenaDNA causal) | (det.) | 84.28 | 85.72 | 70.68 | 70.87 | 86.95 | 78.51 | 64.72 | 73.97 | 76.96 |
| H1 (+bidirectional patch) | (det.) | 81.86 | 80.48 | 68.77 | 68.14 | 79.17 | 79.47 | 62.75 | 75.62 | 74.53 |
| H2 (+MNTP) | r1 | 85.37 | 85.54 | 70.60 | 71.57 | 82.31 | 80.01 | 65.07 | 76.45 | 77.11 |
| H2 (+MNTP) | r2 | 85.37 | 84.62 | 70.75 | 71.38 | 81.61 | 80.13 | 64.66 | 76.45 | 76.87 |
| H2 (+MNTP) | r3 | 85.48 | 84.51 | 70.68 | 70.16 | 83.41 | 80.21 | 64.75 | 75.21 | 76.80 |
| H2 (+MNTP) | mean±std | 85.41±0.06 | 84.89±0.57 | 70.68±0.08 | 71.04±0.77 | 82.44±0.91 | 80.12±0.10 | 64.83±0.22 | 76.04±0.72 | 76.93±0.16 |
| H3 (+dropout SimCSE) | r1 | 84.92 | 87.34 | 69.13 | 69.40 | 92.76 | 81.02 | 64.05 | 74.79 | 77.93 |
| H3 (+dropout SimCSE) | r2 | 85.18 | 88.27 | 70.05 | 69.36 | 90.36 | 81.27 | 64.16 | 78.10 | 78.34 |
| H3 (+dropout SimCSE) | r3 | 85.63 | 86.84 | 71.56 | 70.24 | 91.89 | 82.24 | 65.40 | 81.40 | 79.40 |
| H3 (+dropout SimCSE) | mean±std | 85.24±0.36 | 87.48±0.73 | 70.25±1.23 | 69.67±0.50 | 91.67±1.22 | 81.51±0.64 | 64.54±0.75 | 78.10±3.31 | 78.56±0.76 |
| H4 (+revcomp SimCSE) | r1 | 85.07 | 90.01 | 71.88 | 71.37 | 87.53 | 81.65 | 65.20 | 80.17 | 79.11 |
| H4 (+revcomp SimCSE) | r2 | 86.30 | 90.73 | 71.49 | 71.48 | 88.04 | 82.17 | 65.95 | 79.75 | 79.49 |
| H4 (+revcomp SimCSE) | r3 | 85.92 | 90.01 | 71.99 | 68.17 | 88.46 | 81.76 | 64.43 | 80.17 | 78.86 |
| H4 (+revcomp SimCSE) | mean±std | 85.76±0.63 | 90.25±0.42 | 71.79±0.26 | 70.34±1.88 | 88.01±0.47 | 81.86±0.27 | 65.19±0.76 | 80.03±0.24 | 79.15±0.32 |
| H5 (+crop SimCSE) | r1 | 83.65 | 84.88 | 71.37 | 69.71 | 81.39 | 81.50 | 63.85 | 76.45 | 76.60 |
| H5 (+crop SimCSE) | r2 | 84.38 | 85.50 | 70.77 | 69.91 | 81.56 | 81.76 | 64.59 | 75.21 | 76.71 |
| H5 (+crop SimCSE) | r3 | 84.54 | 84.91 | 70.84 | 68.38 | 82.84 | 82.12 | 64.66 | 75.21 | 76.69 |
| H5 (+crop SimCSE) | mean±std | 84.19±0.47 | 85.10±0.35 | 70.99±0.33 | 69.33±0.83 | 81.93±0.79 | 81.79±0.31 | 64.37±0.45 | 75.62±0.72 | 76.67±0.06 |
| H6 (+local-shift SimCSE) | r1 | 85.61 | 88.09 | 71.20 | 70.91 | 91.22 | 82.28 | 64.61 | 76.45 | 78.80 |
| H6 (+local-shift SimCSE) | r2 | 87.11 | 89.57 | 71.19 | 71.66 | 90.06 | 81.86 | 66.13 | 76.86 | 79.30 |
| H6 (+local-shift SimCSE) | r3 | 86.44 | 88.64 | 71.92 | 69.96 | 90.68 | 82.49 | 65.79 | 77.69 | 79.20 |
| H6 (+local-shift SimCSE) | mean±std | 86.39±0.75 | 88.77±0.75 | 71.44±0.42 | 70.84±0.85 | 90.65±0.58 | 82.21±0.32 | 65.51±0.80 | 77.00±0.63 | 79.10±0.26 |

Table S9: HyenaDNA Nucleotide Transformer downstream tasks (NT) per-seed linear probe MCC (%), part A: promoter, enhancer, splice, and aggregate tasks.

| **Model** | **Run** | **Prom-All** | **Prom-TATA** | **Prom-noTA** | **Enhancers** | **Enh-Types** | **Splice-All** | **Splice-Acc** | **Splice-Donor** | **Avg-NT** |
|---|---|---|---|---|---|---|---|---|---|---|
| H0 (HyenaDNA causal) | (det.) | 74.03 | 63.42 | 74.14 | 46.53 | 29.00 | 22.00 | 23.48 | 21.87 | 35.29 |
| H1 (+bidirectional patch) | (det.) | 71.92 | 66.38 | 73.19 | 50.50 | 32.25 | 17.91 | 20.43 | 19.57 | 37.98 |
| H2 (+MNTP) | r1 | 74.21 | 60.13 | 73.82 | 44.75 | 26.23 | 19.27 | 24.18 | 23.86 | 36.83 |
| H2 (+MNTP) | r2 | 73.44 | 58.93 | 72.41 | 45.27 | 28.64 | 19.64 | 28.36 | 24.63 | 36.50 |
| H2 (+MNTP) | r3 | 73.54 | 60.28 | 73.50 | 44.27 | 23.65 | 19.41 | 25.30 | 23.35 | 36.21 |
| H2 (+MNTP) | mean±std | 73.73±0.42 | 59.78±0.74 | 73.24±0.74 | 44.76±0.50 | 26.17±2.50 | 19.44±0.19 | 25.95±2.16 | 23.95±0.64 | 36.51±0.31 |
| H3 (+dropout SimCSE) | r1 | 77.43 | 63.71 | 78.18 | 50.50 | 32.24 | 31.88 | 30.44 | 27.95 | 42.03 |
| H3 (+dropout SimCSE) | r2 | 76.63 | 68.65 | 76.23 | 46.06 | 31.55 | 29.22 | 28.54 | 26.28 | 41.80 |
| H3 (+dropout SimCSE) | r3 | 78.04 | 68.57 | 77.95 | 47.00 | 28.10 | 28.04 | 27.58 | 24.24 | 40.95 |
| H3 (+dropout SimCSE) | mean±std | 77.37±0.71 | 66.98±2.83 | 77.45±1.07 | 47.85±2.34 | 30.63±2.22 | 29.71±1.97 | 28.85±1.46 | 26.16±1.86 | 41.59±0.57 |
| H4 (+revcomp SimCSE) | r1 | 77.28 | 70.02 | 77.10 | 49.06 | 33.63 | 25.77 | 26.52 | 28.63 | 43.75 |
| H4 (+revcomp SimCSE) | r2 | 77.41 | 69.28 | 77.25 | 48.12 | 34.11 | 28.79 | 30.40 | 25.32 | 43.77 |
| H4 (+revcomp SimCSE) | r3 | 75.15 | 66.74 | 75.21 | 46.63 | 30.73 | 27.74 | 30.18 | 30.10 | 42.07 |
| H4 (+revcomp SimCSE) | mean±std | 76.61±1.27 | 68.68±1.72 | 76.52±1.14 | 47.94±1.23 | 32.82±1.83 | 27.43±1.53 | 29.03±2.18 | 28.02±2.45 | 43.20±0.98 |
| H5 (+crop SimCSE) | r1 | 78.15 | 74.16 | 78.22 | 45.83 | 33.15 | 24.58 | 26.37 | 27.03 | 42.70 |
| H5 (+crop SimCSE) | r2 | 79.22 | 71.60 | 80.20 | 42.31 | 29.67 | 23.16 | 30.69 | 26.15 | 42.99 |
| H5 (+crop SimCSE) | r3 | 78.53 | 75.06 | 78.50 | 47.07 | 29.08 | 24.91 | 28.81 | 27.80 | 42.96 |
| H5 (+crop SimCSE) | mean±std | 78.63±0.54 | 73.61±1.80 | 78.97±1.07 | 45.07±2.47 | 30.63±2.20 | 24.22±0.93 | 28.62±2.17 | 26.99±0.83 | 42.88±0.16 |
| H6 (+local-shift SimCSE) | r1 | 80.32 | 77.39 | 80.31 | 45.18 | 29.34 | 29.02 | 33.20 | 29.16 | 45.01 |
| H6 (+local-shift SimCSE) | r2 | 80.68 | 75.84 | 80.98 | 45.57 | 28.66 | 34.92 | 34.45 | 35.00 | 45.48 |
| H6 (+local-shift SimCSE) | r3 | 81.05 | 76.12 | 81.35 | 43.59 | 27.48 | 29.77 | 30.33 | 28.97 | 44.21 |
| H6 (+local-shift SimCSE) | mean±std | 80.68±0.37 | 76.45±0.83 | 80.88±0.53 | 44.78±1.05 | 28.49±0.94 | 31.24±3.21 | 32.66±2.11 | 31.04±3.43 | 44.90±0.64 |

Table S10: HyenaDNA Nucleotide Transformer downstream tasks (NT) per-seed linear probe MCC (%), part B: histone-mark tasks and aggregate.

| Model | Run | H3 | H4 | H3K9ac | H3K14ac | H4ac | H3K4me1 | H3K4me2 | H3K4me3 | H3K36me3 | H3K79me3 | Avg-NT |
|---|---|---|---|---|---|---|---|---|---|---|---|---|
| H0 (HyenaDNA causal) | (det.) | 48.25 | 40.71 | 30.52 | 12.99 | 20.18 | 18.90 | 22.54 | 18.32 | 27.73 | 40.70 | 35.29 |
| H1 (+bidirectional patch) | (det.) | 57.48 | 58.80 | 35.65 | 19.50 | 21.47 | 23.34 | 23.31 | 14.08 | 32.80 | 45.15 | 37.98 |
| H2 (+MNTP) | r1 | 55.49 | 52.41 | 30.72 | 15.64 | 21.68 | 26.49 | 23.04 | 15.47 | 29.57 | 45.93 | 36.83 |
| H2 (+MNTP) | r2 | 55.52 | 49.96 | 31.10 | 14.51 | 22.24 | 23.44 | 23.32 | 15.11 | 27.29 | 43.20 | 36.50 |
| H2 (+MNTP) | r3 | 56.29 | 48.52 | 30.75 | 13.89 | 22.23 | 23.68 | 24.26 | 15.59 | 28.85 | 44.39 | 36.21 |
| H2 (+MNTP) | mean±std | 55.77±0.45 | 50.30±1.97 | 30.86±0.21 | 14.68±0.89 | 22.05±0.32 | 24.54±1.70 | 23.54±0.64 | 15.39±0.25 | 28.57±1.17 | 44.51±1.37 | 36.51±0.31 |
| H3 (+dropout SimCSE) | r1 | 65.08 | 63.06 | 37.98 | 22.89 | 24.40 | 30.24 | 24.24 | 16.18 | 33.22 | 46.89 | 42.03 |
| H3 (+dropout SimCSE) | r2 | 62.39 | 64.06 | 38.78 | 24.06 | 25.44 | 29.60 | 24.93 | 18.65 | 33.78 | 47.59 | 41.80 |
| H3 (+dropout SimCSE) | r3 | 63.19 | 61.38 | 34.37 | 23.30 | 25.61 | 28.25 | 24.03 | 18.15 | 34.27 | 44.99 | 40.95 |
| H3 (+dropout SimCSE) | mean±std | 63.55±1.38 | 62.83±1.35 | 37.04±2.35 | 23.42±0.59 | 25.15±0.66 | 29.36±1.02 | 24.40±0.47 | 17.66±1.31 | 33.76±0.53 | 46.49±1.35 | 41.59±0.57 |
| H4 (+revcomp SimCSE) | r1 | 68.46 | 69.67 | 40.49 | 28.37 | 28.62 | 29.56 | 25.58 | 18.99 | 38.51 | 51.16 | 43.75 |
| H4 (+revcomp SimCSE) | r2 | 70.45 | 69.83 | 40.04 | 26.59 | 29.42 | 28.16 | 25.59 | 18.68 | 37.36 | 51.10 | 43.77 |
| H4 (+revcomp SimCSE) | r3 | 66.44 | 63.67 | 38.59 | 22.70 | 28.14 | 28.48 | 23.23 | 20.33 | 35.16 | 48.01 | 42.07 |
| H4 (+revcomp SimCSE) | mean±std | 68.45±2.01 | 67.72±3.51 | 39.71±0.99 | 25.89±2.90 | 28.73±0.65 | 28.73±0.73 | 24.80±1.36 | 19.33±0.88 | 37.01±1.70 | 50.09±1.80 | 43.20±0.98 |
| H5 (+crop SimCSE) | r1 | 64.98 | 68.54 | 37.05 | 24.41 | 26.92 | 30.14 | 23.86 | 17.03 | 38.18 | 49.90 | 42.70 |
| H5 (+crop SimCSE) | r2 | 66.99 | 69.18 | 37.20 | 24.73 | 29.61 | 30.35 | 23.75 | 19.68 | 40.08 | 49.34 | 42.99 |
| H5 (+crop SimCSE) | r3 | 64.98 | 64.35 | 38.84 | 25.88 | 27.79 | 32.18 | 25.08 | 18.92 | 36.77 | 48.78 | 42.96 |
| H5 (+crop SimCSE) | mean±std | 65.65±1.16 | 67.36±2.62 | 37.70±0.99 | 25.01±0.77 | 28.11±1.37 | 30.89±1.12 | 24.23±0.74 | 18.54±1.36 | 38.34±1.66 | 49.34±0.56 | 42.88±0.16 |
| H6 (+local-shift SimCSE) | r1 | 65.94 | 72.71 | 38.53 | 29.73 | 28.93 | 32.90 | 25.72 | 20.66 | 39.59 | 51.51 | 45.01 |
| H6 (+local-shift SimCSE) | r2 | 65.39 | 68.10 | 38.27 | 30.34 | 29.01 | 33.96 | 25.62 | 18.73 | 41.03 | 52.00 | 45.48 |
| H6 (+local-shift SimCSE) | r3 | 66.04 | 68.72 | 37.89 | 27.66 | 28.54 | 32.49 | 25.34 | 20.00 | 39.23 | 51.17 | 44.21 |
| H6 (+local-shift SimCSE) | mean±std | 65.79±0.35 | 69.84±2.50 | 38.23±0.32 | 29.24±1.40 | 28.83±0.25 | 33.12±0.76 | 25.56±0.20 | 19.80±0.98 | 39.95±0.95 | 51.56±0.42 | 44.90±0.64 |

Table S11: HyenaDNA GUE+ EPI per-seed linear probe MCC (%).

| Model | Run | GM12878 | HeLa-S3 | HUVEC | IMR90 | K562 | NHEK | Avg-EPI |
|---|---|---|---|---|---|---|---|---|
| H0 (HyenaDNA causal) | (det.) | 12.62 | 13.10 | 21.82 | 21.86 | 13.76 | 12.15 | 15.88 |
| H1 (+bidirectional patch) | (det.) | 11.15 | 11.19 | 17.57 | 15.26 | 8.97 | 4.20 | 11.39 |
| H2 (+MNTP) | r1 | 12.40 | 11.84 | 21.00 | 23.61 | 14.90 | 13.54 | 16.22 |
| H2 (+MNTP) | r2 | 13.67 | 12.21 | 21.21 | 23.94 | 14.52 | 12.08 | 16.27 |
| H2 (+MNTP) | r3 | 12.31 | 12.79 | 22.53 | 24.22 | 13.56 | 14.47 | 16.65 |
| H2 (+MNTP) | mean±std | 12.79±0.76 | 12.28±0.48 | 21.58±0.83 | 23.92±0.31 | 14.33±0.69 | 13.36±1.20 | 16.38±0.24 |
| H3 (+dropout SimCSE) | r1 | 21.76 | 17.47 | 26.48 | 23.17 | 25.86 | 20.67 | 22.57 |
| H3 (+dropout SimCSE) | r2 | 19.68 | 21.61 | 23.20 | 25.51 | 23.44 | 21.37 | 22.47 |
| H3 (+dropout SimCSE) | r3 | 19.44 | 19.63 | 25.34 | 25.71 | 24.24 | 18.53 | 22.15 |
| H3 (+dropout SimCSE) | mean±std | 20.29±1.28 | 19.57±2.07 | 25.01±1.67 | 24.80±1.41 | 24.51±1.23 | 20.19±1.48 | 22.40±0.22 |
| H4 (+revcomp SimCSE) | r1 | 20.41 | 15.75 | 24.83 | 22.93 | 17.81 | 19.31 | 20.17 |
| H4 (+revcomp SimCSE) | r2 | 19.48 | 18.31 | 25.91 | 27.49 | 20.23 | 20.26 | 21.95 |
| H4 (+revcomp SimCSE) | r3 | 18.11 | 16.63 | 28.25 | 26.65 | 21.25 | 16.48 | 21.23 |
| H4 (+revcomp SimCSE) | mean±std | 19.33±1.16 | 16.90±1.30 | 26.33±1.75 | 25.69±2.43 | 19.76±1.77 | 18.68±1.97 | 21.12±0.90 |
| H5 (+crop SimCSE) | r1 | 17.98 | 23.42 | 31.18 | 30.59 | 27.30 | 28.12 | 26.43 |
| H5 (+crop SimCSE) | r2 | 24.46 | 26.12 | 31.60 | 31.63 | 23.78 | 24.29 | 26.98 |
| H5 (+crop SimCSE) | r3 | 23.09 | 25.43 | 30.86 | 32.15 | 21.92 | 25.78 | 26.54 |
| H5 (+crop SimCSE) | mean±std | 21.84±3.42 | 24.99±1.40 | 31.21±0.37 | 31.46±0.79 | 24.33±2.73 | 26.06±1.93 | 26.65±0.29 |
| H6 (+local-shift SimCSE) | r1 | 21.05 | 20.26 | 25.66 | 26.83 | 21.63 | 21.69 | 22.85 |
| H6 (+local-shift SimCSE) | r2 | 23.72 | 20.02 | 28.64 | 27.78 | 24.60 | 23.67 | 24.74 |
| H6 (+local-shift SimCSE) | r3 | 20.46 | 20.98 | 25.69 | 32.12 | 25.57 | 22.34 | 24.53 |
| H6 (+local-shift SimCSE) | mean±std | 21.74±1.74 | 20.42±0.50 | 26.66±1.71 | 28.91±2.82 | 23.93±2.05 | 22.57±1.01 | 24.04±1.04 |

Table S12: HyenaDNA GUE per-seed linear probe MCC (%), promoter and human TF-binding tasks.

| Model | Run | PC-All | PC-TATA | PC-noTA | P3-All | P3-TATA | P3-noTA | TF-H0 | TF-H1 | TF-H2 | TF-H3 | TF-H4 |
|---|---|---|---|---|---|---|---|---|---|---|---|---|
| H0 (HyenaDNA causal) | (det.) | 45.07 | 35.73 | 50.78 | 68.73 | 25.68 | 75.95 | 43.74 | 47.29 | 42.77 | 38.63 | 48.84 |
| H1 (+bidirectional patch) | (det.) | 46.21 | 37.70 | 52.66 | 65.36 | 22.18 | 74.19 | 45.75 | 43.65 | 40.51 | 24.55 | 39.60 |
| H2 (+MNTP) | r1 | 48.14 | 41.29 | 53.23 | 66.47 | 26.16 | 75.15 | 47.87 | 48.13 | 42.70 | 37.62 | 49.02 |
| H2 (+MNTP) | r2 | 48.64 | 42.90 | 53.94 | 66.46 | 26.76 | 74.68 | 47.69 | 47.72 | 40.32 | 37.91 | 49.73 |
| H2 (+MNTP) | r3 | 48.13 | 42.61 | 53.84 | 67.07 | 28.75 | 74.08 | 49.84 | 46.72 | 39.00 | 36.27 | 48.67 |
| H2 (+MNTP) | mean±std | 48.30±0.29 | 42.27±0.86 | 53.67±0.38 | 66.67±0.35 | 27.22±1.36 | 74.64±0.54 | 48.47±1.19 | 47.52±0.73 | 40.67±1.88 | 37.27±0.88 | 49.14±0.54 |
| H3 (+dropout SimCSE) | r1 | 50.92 | 39.33 | 57.87 | 71.82 | 41.68 | 78.00 | 51.52 | 47.04 | 43.00 | 27.48 | 42.60 |
| H3 (+dropout SimCSE) | r2 | 52.74 | 43.46 | 59.51 | 70.34 | 43.01 | 76.13 | 50.32 | 46.06 | 45.81 | 32.54 | 42.62 |
| H3 (+dropout SimCSE) | r3 | 52.04 | 39.47 | 59.24 | 72.57 | 39.84 | 78.66 | 50.57 | 51.87 | 41.25 | 35.78 | 46.87 |
| H3 (+dropout SimCSE) | mean±std | 51.90±0.92 | 40.75±2.35 | 58.87±0.88 | 71.58±1.13 | 41.51±1.59 | 77.60±1.31 | 50.80±0.63 | 48.32±3.11 | 43.35±2.30 | 31.93±4.18 | 44.03±2.46 |
| H4 (+revcomp SimCSE) | r1 | 51.10 | 40.52 | 59.41 | 70.51 | 34.01 | 78.62 | 48.81 | 51.18 | 40.50 | 35.90 | 47.20 |
| H4 (+revcomp SimCSE) | r2 | 53.36 | 41.65 | 61.54 | 71.19 | 33.47 | 78.84 | 51.51 | 49.96 | 43.47 | 35.26 | 48.81 |
| H4 (+revcomp SimCSE) | r3 | 51.40 | 41.51 | 58.52 | 69.83 | 34.20 | 76.63 | 50.38 | 49.46 | 38.73 | 36.76 | 47.22 |
| H4 (+revcomp SimCSE) | mean±std | 51.95±1.23 | 41.23±0.62 | 59.82±1.55 | 70.51±0.68 | 33.89±0.38 | 78.03±1.22 | 50.23±1.36 | 50.20±0.88 | 40.90±2.40 | 35.97±0.75 | 47.74±0.92 |
| H5 (+crop SimCSE) | r1 | 50.22 | 42.63 | 56.79 | 71.49 | 45.15 | 78.64 | 49.32 | 47.20 | 42.91 | 30.34 | 41.99 |
| H5 (+crop SimCSE) | r2 | 50.45 | 45.53 | 58.05 | 71.96 | 40.20 | 80.45 | 50.12 | 46.20 | 42.63 | 29.32 | 42.01 |
| H5 (+crop SimCSE) | r3 | 50.37 | 41.67 | 55.75 | 71.76 | 39.86 | 78.95 | 48.51 | 43.20 | 42.82 | 27.01 | 40.99 |
| H5 (+crop SimCSE) | mean±std | 50.35±0.12 | 43.28±2.01 | 56.86±1.15 | 71.74±0.24 | 41.74±2.96 | 79.35±0.97 | 49.32±0.81 | 45.53±2.08 | 42.79±0.14 | 28.89±1.71 | 41.66±0.58 |
| H6 (+local-shift SimCSE) | r1 | 51.78 | 41.00 | 57.88 | 74.76 | 41.13 | 81.33 | 51.63 | 52.51 | 44.47 | 33.48 | 48.42 |
| H6 (+local-shift SimCSE) | r2 | 53.23 | 45.87 | 60.75 | 74.39 | 46.97 | 81.39 | 54.96 | 50.33 | 44.12 | 36.65 | 52.26 |
| H6 (+local-shift SimCSE) | r3 | 52.42 | 42.33 | 59.63 | 73.28 | 38.40 | 81.08 | 50.19 | 49.77 | 42.69 | 33.03 | 50.82 |
| H6 (+local-shift SimCSE) | mean±std | 52.48±0.73 | 43.07±2.52 | 59.42±1.45 | 74.14±0.77 | 42.17±4.38 | 81.27±0.16 | 52.26±2.45 | 50.87±1.45 | 43.76±0.94 | 34.39±1.97 | 50.50±1.94 |

Table S13: HyenaDNA GUE per-seed linear probe MCC (%), mouse TF-binding, splice, Virus-COVID MCC, and Avg-GUE17.

| Model | Run | TF-M0 | TF-M1 | TF-M2 | TF-M3 | TF-M4 | Splice | COVID-MCC | Avg-GUE17 |
|---|---|---|---|---|---|---|---|---|---|
| H0 (HyenaDNA causal) | (det.) | 15.03 | 59.56 | 48.88 | 24.55 | 16.15 | 2.10 | 3.36 | 40.56 |
| H1 (+bidirectional patch) | (det.) | 11.57 | 58.06 | 50.13 | 11.29 | 16.07 | 0.22 | 3.51 | 37.63 |
| H2 (+MNTP) | r1 | 15.36 | 61.14 | 51.31 | 13.85 | 19.16 | -0.78 | 2.93 | 40.93 |
| H2 (+MNTP) | r2 | 19.16 | 61.53 | 55.67 | 12.35 | 19.06 | 2.93 | 2.56 | 41.61 |
| H2 (+MNTP) | r3 | 17.65 | 61.57 | 52.71 | 7.94 | 17.78 | -0.07 | 2.83 | 40.74 |
| H2 (+MNTP) | mean±std | 17.39±1.91 | 61.41±0.24 | 53.23±2.23 | 11.38±3.07 | 18.67±0.77 | 0.69±1.97 | 2.77±0.19 | 41.09±0.46 |
| H3 (+dropout SimCSE) | r1 | 22.31 | 59.16 | 42.69 | 3.76 | 15.69 | 15.31 | 8.73 | 41.78 |
| H3 (+dropout SimCSE) | r2 | 18.03 | 59.53 | 39.81 | 10.48 | 13.51 | 12.12 | 9.40 | 42.12 |
| H3 (+dropout SimCSE) | r3 | 12.87 | 64.09 | 57.97 | 8.78 | 15.90 | 7.76 | 8.72 | 43.27 |
| H3 (+dropout SimCSE) | mean±std | 17.74±4.73 | 60.93±2.75 | 46.82±9.76 | 7.67±3.49 | 15.03±1.32 | 11.73±3.79 | 8.95±0.39 | 42.39±0.78 |
| H4 (+revcomp SimCSE) | r1 | 12.20 | 64.47 | 45.85 | 10.46 | 21.50 | 6.14 | 6.74 | 42.26 |
| H4 (+revcomp SimCSE) | r2 | 19.49 | 65.29 | 58.59 | 5.50 | 17.24 | 10.81 | 7.59 | 43.88 |
| H4 (+revcomp SimCSE) | r3 | 14.69 | 63.16 | 53.95 | 17.17 | 15.11 | 6.58 | 6.14 | 42.66 |
| H4 (+revcomp SimCSE) | mean±std | 15.46±3.71 | 64.31±1.07 | 52.80±6.45 | 11.04±5.86 | 17.95±3.25 | 7.84±2.58 | 6.82±0.73 | 42.93±0.84 |
| H5 (+crop SimCSE) | r1 | 15.79 | 57.34 | 48.82 | 10.46 | 17.33 | 12.83 | 23.82 | 42.31 |
| H5 (+crop SimCSE) | r2 | 13.90 | 55.74 | 55.55 | 11.30 | 19.31 | 11.80 | 22.20 | 42.62 |
| H5 (+crop SimCSE) | r3 | 17.50 | 53.67 | 48.78 | 11.52 | 18.20 | 11.95 | 24.36 | 41.32 |
| H5 (+crop SimCSE) | mean±std | 15.73±1.80 | 55.58±1.84 | 51.05±3.90 | 11.09±0.56 | 18.28±0.99 | 12.19±0.56 | 23.46±1.12 | 42.08±0.68 |
| H6 (+local-shift SimCSE) | r1 | 21.39 | 60.05 | 54.27 | 17.22 | 22.34 | 15.43 | 12.93 | 45.24 |
| H6 (+local-shift SimCSE) | r2 | 30.53 | 66.06 | 62.24 | 19.71 | 26.54 | 18.10 | 13.71 | 48.48 |
| H6 (+local-shift SimCSE) | r3 | 22.40 | 62.04 | 57.94 | 23.85 | 24.79 | 13.38 | 13.54 | 45.77 |
| H6 (+local-shift SimCSE) | mean±std | 24.77±5.01 | 62.72±3.06 | 58.15±3.99 | 20.26±3.35 | 24.56±2.11 | 15.64±2.37 | 13.39±0.41 | 46.49±1.74 |

Table S14: HyenaDNA Virus-COVID macro-F1 under the Step 4 mean-pooling linear-probe protocol.

| Model | Run | Virus-COVID F1 |
|---|---|---|
| H0 (HyenaDNA causal) | (det.) | 11.33 |
| H1 (+bidirectional patch) | (det.) | 11.32 |
| H2 (+MNTP) | r1 | 11.81 |
| H2 (+MNTP) | r2 | 11.64 |
| H2 (+MNTP) | r3 | 11.40 |
| H2 (+MNTP) | mean±std | 11.62±0.21 |
| H3 (+dropout SimCSE) | r1 | 17.34 |
| H3 (+dropout SimCSE) | r2 | 17.59 |
| H3 (+dropout SimCSE) | r3 | 17.15 |
| H3 (+dropout SimCSE) | mean±std | 17.36±0.22 |
| H4 (+revcomp SimCSE) | r1 | 16.02 |
| H4 (+revcomp SimCSE) | r2 | 16.49 |
| H4 (+revcomp SimCSE) | r3 | 14.76 |
| H4 (+revcomp SimCSE) | mean±std | 15.76±0.89 |
| H5 (+crop SimCSE) | r1 | 31.82 |
| H5 (+crop SimCSE) | r2 | 30.39 |
| H5 (+crop SimCSE) | r3 | 32.23 |
| H5 (+crop SimCSE) | mean±std | 31.48±0.97 |
| H6 (+local-shift SimCSE) | r1 | 20.77 |
| H6 (+local-shift SimCSE) | r2 | 21.27 |
| H6 (+local-shift SimCSE) | r3 | 21.22 |
| H6 (+local-shift SimCSE) | mean±std | 21.09±0.28 |

tuning summary; HyenaDNA rows are generated from the preserve-dtype (fp32-faithful) `exp_res/hyena_gated/step5-ep3-pd-full-*` workbooks.

Table S15: DNAGPT Step 5 full fine-tuning GUE+ EPI MCC (%) per seed.

| Model | Run | GM12878 | HeLa-S3 | HUVEC | IMR90 | K562 | NHEK | Avg-EPI |
|---|---|---|---|---|---|---|---|---|
| M0 (DNAGPT causal) | r1 | 55.65 | 61.97 | 64.81 | 75.13 | 58.29 | 74.56 | 65.07 |
| M0 (DNAGPT causal) | r2 | 64.61 | 69.50 | 67.08 | 79.97 | 65.87 | 80.42 | 71.24 |
| M0 (DNAGPT causal) | r3 | 61.30 | 68.72 | 68.94 | 78.00 | 67.32 | 82.21 | 71.08 |
| M0 (DNAGPT causal) | mean±std | 60.52±4.53 | 66.73±4.14 | 66.94±2.07 | 77.70±2.43 | 63.83±4.85 | 79.06±4.00 | 69.13±3.52 |
| M1 (+bidir attn) | r1 | 63.00 | 71.04 | 62.87 | 73.98 | 63.26 | 82.91 | 69.51 |
| M1 (+bidir attn) | r2 | 71.99 | 77.99 | 74.10 | 81.06 | 75.42 | 89.50 | 78.34 |
| M1 (+bidir attn) | r3 | 71.24 | 79.44 | 74.81 | 77.32 | 74.22 | 88.20 | 77.54 |
| M1 (+bidir attn) | mean±std | 68.74±4.99 | 76.16±4.49 | 70.59±6.70 | 77.45±3.54 | 70.97±6.70 | 86.87±3.49 | 75.13±4.88 |
| M2 (+MNTP) | r1 | 60.34 | 68.66 | 70.22 | 72.60 | 61.85 | 81.54 | 69.20 |
| M2 (+MNTP) | r2 | 68.80 | 75.15 | 73.14 | 79.32 | 69.70 | 84.94 | 75.17 |
| M2 (+MNTP) | r3 | 65.47 | 77.06 | 72.18 | 76.62 | 69.52 | 83.64 | 74.08 |
| M2 (+MNTP) | mean±std | 64.87±4.26 | 73.62±4.40 | 71.85±1.49 | 76.18±3.38 | 67.02±4.48 | 83.37±1.72 | 72.82±3.18 |
| M3 (+dropout SimCSE) | r1 | 60.30 | 72.70 | 69.87 | 74.82 | 64.17 | 83.84 | 70.95 |
| M3 (+dropout SimCSE) | r2 | 66.23 | 79.52 | 72.38 | 81.48 | 70.91 | 86.62 | 76.19 |
| M3 (+dropout SimCSE) | r3 | 69.54 | 79.35 | 75.42 | 80.49 | 68.10 | 86.24 | 76.52 |
| M3 (+dropout SimCSE) | mean±std | 65.36±4.68 | 77.19±3.89 | 72.56±2.78 | 78.93±3.59 | 67.73±3.39 | 85.57±1.51 | 74.55±3.12 |
| M4 (+RC SimCSE) | r1 | 55.87 | 76.66 | 69.82 | 78.84 | 66.89 | 84.45 | 72.09 |
| M4 (+RC SimCSE) | r2 | 71.93 | 79.45 | 76.52 | 81.64 | 76.12 | 88.20 | 78.98 |
| M4 (+RC SimCSE) | r3 | 68.34 | 81.26 | 77.26 | 83.66 | 74.29 | 87.35 | 78.69 |
| M4 (+RC SimCSE) | mean±std | 65.38±8.43 | 79.12±2.32 | 74.53±4.10 | 81.38±2.42 | 72.43±4.89 | 86.67±1.97 | 76.59±3.90 |
| M5 (+crop SimCSE) | r1 | 63.36 | 76.09 | 67.60 | 74.51 | 70.62 | 82.01 | 72.36 |
| M5 (+crop SimCSE) | r2 | 70.95 | 80.98 | 74.28 | 82.32 | 73.50 | 86.39 | 78.07 |
| M5 (+crop SimCSE) | r3 | 68.21 | 79.00 | 73.55 | 81.23 | 74.91 | 87.50 | 77.40 |
| M5 (+crop SimCSE) | mean±std | 67.51±3.84 | 78.69±2.46 | 71.81±3.66 | 79.35±4.23 | 73.01±2.19 | 85.30±2.90 | 75.94±3.12 |
| M6 (+local-shift) | r1 | 63.86 | 72.11 | 67.26 | 74.04 | 65.76 | 82.76 | 70.97 |
| M6 (+local-shift) | r2 | 72.19 | 78.57 | 74.16 | 79.97 | 74.02 | 84.74 | 77.28 |
| M6 (+local-shift) | r3 | 71.28 | 76.42 | 75.06 | 82.03 | 75.32 | 86.43 | 77.76 |
| M6 (+local-shift) | mean±std | 69.11±4.57 | 75.70±3.29 | 72.16±4.27 | 78.68±4.15 | 71.70±5.19 | 84.64±1.84 | 75.34±3.79 |

Table S16: DNAGPT Step 5 full fine-tuning GUE MCC (%) per seed, promoter and human TF-binding tasks.

| Model | Run | PC-All | PC-TATA | PC-noTA | P3-All | P3-TATA | P3-noTA | TF-H0 | TF-H1 | TF-H2 | TF-H3 | TF-H4 |
|---|---|---|---|---|---|---|---|---|---|---|---|---|
| M0 (DNAGPT causal) | r1 | 60.16 | 54.70 | 66.88 | 84.36 | 56.50 | 91.52 | 54.79 | 52.64 | 49.07 | 39.89 | 71.86 |
| M0 (DNAGPT causal) | r2 | 61.98 | 55.01 | 67.43 | 84.80 | 56.78 | 91.87 | 57.29 | 52.56 | 51.79 | 44.50 | 69.69 |
| M0 (DNAGPT causal) | r3 | 62.05 | 55.63 | 66.71 | 84.97 | 59.80 | 92.05 | 57.53 | 52.82 | 49.82 | 44.68 | 71.26 |
| M0 (DNAGPT causal) | mean±std | 61.40±1.07 | 55.11±0.47 | 67.01±0.38 | 84.71±0.31 | 57.69±1.83 | 91.81±0.27 | 56.54±1.52 | 52.67±0.13 | 50.23±1.40 | 43.02±2.72 | 70.94±1.12 |
| M1 (+bidir attn) | r1 | 60.63 | 50.42 | 66.21 | 83.24 | 55.18 | 90.99 | 56.80 | 54.26 | 48.11 | 45.71 | 70.06 |
| M1 (+bidir attn) | r2 | 60.83 | 49.27 | 66.82 | 84.87 | 58.42 | 91.60 | 57.56 | 56.04 | 46.69 | 48.48 | 70.41 |
| M1 (+bidir attn) | r3 | 60.13 | 47.71 | 66.60 | 84.49 | 58.09 | 91.36 | 58.80 | 55.66 | 48.85 | 46.97 | 70.49 |
| M1 (+bidir attn) | mean±std | 60.53±0.36 | 49.13±1.36 | 66.54±0.31 | 84.20±0.85 | 57.23±1.78 | 91.32±0.31 | 57.72±1.01 | 55.32±0.94 | 47.88±1.10 | 47.05±1.39 | 70.32±0.23 |
| M2 (+MNTP) | r1 | 59.95 | 49.10 | 66.07 | 83.86 | 53.82 | 91.75 | 56.74 | 53.60 | 47.74 | 44.11 | 71.03 |
| M2 (+MNTP) | r2 | 60.89 | 51.71 | 66.90 | 84.87 | 55.82 | 91.86 | 58.61 | 55.11 | 49.38 | 46.02 | 70.00 |
| M2 (+MNTP) | r3 | 61.70 | 47.15 | 66.76 | 84.63 | 55.80 | 92.22 | 60.34 | 54.13 | 48.54 | 45.43 | 68.80 |
| M2 (+MNTP) | mean±std | 60.85±0.88 | 49.32±2.29 | 66.58±0.44 | 84.45±0.53 | 55.15±1.15 | 91.94±0.25 | 58.56±1.80 | 54.28±0.77 | 48.55±0.82 | 45.19±0.98 | 69.94±1.12 |
| M3 (+dropout SimCSE) | r1 | 59.45 | 53.34 | 65.51 | 84.20 | 53.91 | 92.54 | 54.95 | 53.60 | 48.70 | 42.68 | 67.96 |
| M3 (+dropout SimCSE) | r2 | 60.68 | 50.19 | 66.30 | 84.06 | 53.90 | 91.82 | 57.35 | 55.41 | 50.47 | 47.30 | 67.20 |
| M3 (+dropout SimCSE) | r3 | 60.38 | 51.06 | 66.82 | 84.33 | 54.49 | 91.27 | 57.96 | 54.84 | 50.65 | 46.10 | 70.27 |
| M3 (+dropout SimCSE) | mean±std | 60.17±0.64 | 51.53±1.63 | 66.21±0.66 | 84.20±0.14 | 54.10±0.34 | 91.88±0.64 | 56.75±1.59 | 54.62±0.93 | 49.94±1.08 | 45.36±2.40 | 68.48±1.60 |
| M4 (+RC SimCSE) | r1 | 60.87 | 48.78 | 65.37 | 83.50 | 61.53 | 92.33 | 56.93 | 54.80 | 45.89 | 40.11 | 69.20 |
| M4 (+RC SimCSE) | r2 | 60.81 | 49.09 | 66.44 | 84.24 | 59.41 | 92.05 | 57.89 | 54.90 | 48.68 | 44.54 | 70.67 |
| M4 (+RC SimCSE) | r3 | 59.96 | 50.44 | 66.53 | 84.60 | 57.23 | 91.91 | 59.14 | 56.82 | 49.78 | 45.19 | 69.92 |
| M4 (+RC SimCSE) | mean±std | 60.55±0.51 | 49.44±0.88 | 66.11±0.65 | 84.11±0.56 | 59.39±2.15 | 92.10±0.21 | 57.99±1.11 | 55.51±1.14 | 48.12±2.01 | 43.28±2.76 | 69.93±0.74 |
| M5 (+crop SimCSE) | r1 | 59.87 | 45.84 | 65.77 | 83.45 | 57.86 | 91.50 | 54.73 | 53.44 | 49.68 | 38.33 | 69.48 |
| M5 (+crop SimCSE) | r2 | 60.28 | 47.85 | 66.23 | 83.52 | 58.41 | 91.82 | 57.33 | 54.20 | 50.77 | 44.83 | 68.37 |
| M5 (+crop SimCSE) | r3 | 60.04 | 52.21 | 66.22 | 84.44 | 59.06 | 91.83 | 58.36 | 54.31 | 50.79 | 47.00 | 66.59 |
| M5 (+crop SimCSE) | mean±std | 60.06±0.21 | 48.63±3.26 | 66.07±0.26 | 83.80±0.55 | 58.44±0.60 | 91.72±0.19 | 56.81±1.87 | 53.98±0.47 | 50.41±0.64 | 43.39±4.51 | 68.15±1.46 |
| M6 (+local-shift) | r1 | 60.76 | 47.47 | 66.09 | 83.32 | 56.84 | 91.26 | 55.54 | 53.40 | 48.21 | 43.92 | 68.46 |
| M6 (+local-shift) | r2 | 60.63 | 48.99 | 66.69 | 83.89 | 60.37 | 91.94 | 57.33 | 55.43 | 49.80 | 45.39 | 69.25 |
| M6 (+local-shift) | r3 | 59.90 | 49.30 | 66.54 | 84.64 | 58.77 | 91.35 | 58.94 | 53.65 | 52.86 | 44.01 | 68.36 |
| M6 (+local-shift) | mean±std | 60.43±0.46 | 48.59±0.98 | 66.44±0.31 | 83.95±0.66 | 58.66±1.77 | 91.52±0.37 | 57.27±1.70 | 54.16±1.11 | 50.29±2.36 | 44.44±0.82 | 68.69±0.49 |

### S3 HyenaDNA Full Fine-Tuning Diagnostics

To interpret why HyenaDNA behaves differently under full fine-tuning than under frozen linear probing, we summarise the preserve-dtype Step 5 full fine-tuning results (Section S2, `exp_res/hyena_gated`) at the family level. All runs use identical hyperparameters for fair

Table S17: DNAGPT Step 5 full fine-tuning GUE MCC (%) per seed, mouse TF-binding, splice, and Avg-GUE17.

| **Model** | **Run** | **TF-M0** | **TF-M1** | **TF-M2** | **TF-M3** | **TF-M4** | **Splice** | **Avg-GUE17** |
|---|---|---|---|---|---|---|---|---|
| M0 (DNAGPT causal) | r1 | 39.58 | 78.49 | 62.84 | 54.01 | 38.40 | 72.62 | 60.49 |
| M0 (DNAGPT causal) | r2 | 44.24 | 78.47 | 64.87 | 57.34 | 39.12 | 70.57 | 61.67 |
| M0 (DNAGPT causal) | r3 | 44.14 | 78.77 | 67.37 | 49.00 | 36.85 | 70.95 | 61.44 |
| M0 (DNAGPT causal) | mean±std | 42.65±2.66 | 78.58±0.17 | 65.03±2.27 | 53.45±4.20 | 38.12±1.16 | 71.38±1.09 | 61.20±0.62 |
| M1 (+bidir attn) | r1 | 41.32 | 80.04 | 64.87 | 60.67 | 35.77 | 59.96 | 60.25 |
| M1 (+bidir attn) | r2 | 41.18 | 80.19 | 69.51 | 48.68 | 34.49 | 59.73 | 60.28 |
| M1 (+bidir attn) | r3 | 45.51 | 80.49 | 69.17 | 50.06 | 36.84 | 63.62 | 60.87 |
| M1 (+bidir attn) | mean±std | 42.67±2.46 | 80.24±0.23 | 67.85±2.59 | 53.14±6.56 | 35.70±1.18 | 61.10±2.18 | 60.47±0.35 |
| M2 (+MNTP) | r1 | 36.02 | 78.83 | 63.86 | 48.52 | 33.85 | 63.99 | 58.99 |
| M2 (+MNTP) | r2 | 40.94 | 78.94 | 74.08 | 49.95 | 36.59 | 64.04 | 60.92 |
| M2 (+MNTP) | r3 | 36.80 | 79.66 | 66.74 | 46.53 | 39.12 | 62.82 | 59.83 |
| M2 (+MNTP) | mean±std | 37.92±2.64 | 79.14±0.45 | 68.23±5.27 | 48.33±1.72 | 36.52±2.64 | 63.62±0.69 | 59.92±0.97 |
| M3 (+dropout SimCSE) | r1 | 40.95 | 79.01 | 66.25 | 47.07 | 36.73 | 62.96 | 59.40 |
| M3 (+dropout SimCSE) | r2 | 42.93 | 79.78 | 72.47 | 58.54 | 35.67 | 63.01 | 61.00 |
| M3 (+dropout SimCSE) | r3 | 45.61 | 78.98 | 67.22 | 59.86 | 38.35 | 66.93 | 61.48 |
| M3 (+dropout SimCSE) | mean±std | 43.16±2.34 | 79.26±0.45 | 68.65±3.35 | 55.16±7.03 | 36.92±1.35 | 64.30±2.28 | 60.63±1.09 |
| M4 (+RC SimCSE) | r1 | 43.42 | 79.19 | 71.49 | 54.89 | 37.64 | 65.43 | 60.67 |
| M4 (+RC SimCSE) | r2 | 40.01 | 79.28 | 72.85 | 50.02 | 36.31 | 63.70 | 60.64 |
| M4 (+RC SimCSE) | r3 | 41.75 | 80.03 | 73.19 | 53.31 | 38.30 | 64.70 | 61.34 |
| M4 (+RC SimCSE) | mean±std | 41.73±1.71 | 79.50±0.46 | 72.51±0.90 | 52.74±2.48 | 37.42±1.01 | 64.61±0.87 | 60.88±0.40 |
| M5 (+crop SimCSE) | r1 | 44.96 | 78.28 | 70.91 | 58.19 | 35.91 | 60.39 | 59.92 |
| M5 (+crop SimCSE) | r2 | 41.26 | 79.77 | 76.83 | 57.55 | 36.98 | 63.50 | 61.15 |
| M5 (+crop SimCSE) | r3 | 44.84 | 79.68 | 70.85 | 56.70 | 37.62 | 63.80 | 61.43 |
| M5 (+crop SimCSE) | mean±std | 43.69±2.10 | 79.24±0.84 | 72.86±3.44 | 57.48±0.75 | 36.84±0.86 | 62.56±1.89 | 60.83±0.81 |
| M6 (+local-shift) | r1 | 42.62 | 78.41 | 70.75 | 54.28 | 37.81 | 61.97 | 60.07 |
| M6 (+local-shift) | r2 | 40.81 | 79.32 | 70.81 | 55.16 | 38.26 | 62.02 | 60.95 |
| M6 (+local-shift) | r3 | 42.10 | 79.36 | 70.80 | 58.67 | 37.61 | 65.76 | 61.33 |
| M6 (+local-shift) | mean±std | 41.84±0.93 | 79.03±0.54 | 70.79±0.03 | 56.04±2.32 | 37.89±0.33 | 63.25±2.17 | 60.78±0.65 |

Table S18: DNAGPT Step 5 full fine-tuning Virus-COVID macro-F1 (%) per seed.

| **Model** | **Run** | **Virus-COVID F1** |
|---|---|---|
| M0 (DNAGPT causal) | r1 | 65.05 |
| M0 (DNAGPT causal) | r2 | 64.10 |
| M0 (DNAGPT causal) | r3 | 64.15 |
| M0 (DNAGPT causal) | mean±std | 64.43±0.53 |
| M1 (+bidir attn) | r1 | 62.32 |
| M1 (+bidir attn) | r2 | 63.53 |
| M1 (+bidir attn) | r3 | 60.43 |
| M1 (+bidir attn) | mean±std | 62.09±1.56 |
| M2 (+MNTP) | r1 | 62.66 |
| M2 (+MNTP) | r2 | 63.49 |
| M2 (+MNTP) | r3 | 64.17 |
| M2 (+MNTP) | mean±std | 63.44±0.76 |
| M3 (+dropout SimCSE) | r1 | 64.26 |
| M3 (+dropout SimCSE) | r2 | 63.68 |
| M3 (+dropout SimCSE) | r3 | 63.12 |
| M3 (+dropout SimCSE) | mean±std | 63.69±0.57 |
| M4 (+RC SimCSE) | r1 | 63.57 |
| M4 (+RC SimCSE) | r2 | 64.56 |
| M4 (+RC SimCSE) | r3 | 62.90 |
| M4 (+RC SimCSE) | mean±std | 63.68±0.84 |
| M5 (+crop SimCSE) | r1 | 64.91 |
| M5 (+crop SimCSE) | r2 | 65.24 |
| M5 (+crop SimCSE) | r3 | 64.14 |
| M5 (+crop SimCSE) | mean±std | 64.76±0.56 |
| M6 (+local-shift) | r1 | 65.28 |
| M6 (+local-shift) | r2 | 64.49 |
| M6 (+local-shift) | r3 | 64.78 |
| M6 (+local-shift) | mean±std | 64.85±0.40 |

Table S19: HyenaDNA Step 5 full fine-tuning GUE+ EPI MCC (%) per seed.

| Model | Run | GM12878 | HeLa-S3 | HUVEC | IMR90 | K562 | NHEK | Avg-EPI |
|---|---|---|---|---|---|---|---|---|
| H0 (HyenaDNA causal) | r1 | 84.53 | 90.78 | 90.67 | 96.18 | 88.81 | 97.91 | 91.48 |
| H0 (HyenaDNA causal) | r2 | 88.05 | 94.38 | 93.69 | 94.99 | 87.10 | 98.41 | 92.77 |
| H0 (HyenaDNA causal) | r3 | 89.26 | 93.78 | 91.88 | 95.89 | 90.89 | 98.10 | 93.30 |
| H0 (HyenaDNA causal) | mean±std | 87.28±2.46 | 92.98±1.93 | 92.08±1.52 | 95.69±0.62 | 88.93±1.90 | 98.14±0.25 | 92.52±0.94 |
| H1 (+bidir patch) | r1 | 81.41 | 85.16 | 91.87 | 91.27 | 85.49 | 95.30 | 88.42 |
| H1 (+bidir patch) | r2 | 82.02 | 83.35 | 90.35 | 92.18 | 88.81 | 96.62 | 88.89 |
| H1 (+bidir patch) | r3 | 85.18 | 89.99 | 88.86 | 91.37 | 84.15 | 95.31 | 89.14 |
| H1 (+bidir patch) | mean±std | 82.87±2.02 | 86.17±3.43 | 90.36±1.51 | 91.61±0.50 | 86.15±2.40 | 95.74±0.76 | 88.82±0.37 |
| H2 (+MNTP) | r1 | 83.09 | 87.20 | 90.58 | 91.64 | 86.87 | 94.10 | 88.92 |
| H2 (+MNTP) | r2 | 81.53 | 94.79 | 88.67 | 92.18 | 83.51 | 95.55 | 89.37 |
| H2 (+MNTP) | r3 | 83.54 | 92.38 | 89.26 | 92.79 | 88.69 | 97.50 | 90.69 |
| H2 (+MNTP) | mean±std | 82.72±1.05 | 91.46±3.88 | 89.50±0.98 | 92.20±0.58 | 86.36±2.63 | 95.72±1.71 | 89.66±0.92 |
| H3 (+dropout SimCSE) | r1 | 73.48 | 80.60 | 82.99 | 87.75 | 78.18 | 88.21 | 81.87 |
| H3 (+dropout SimCSE) | r2 | 75.42 | 78.99 | 85.96 | 88.94 | 79.46 | 91.08 | 83.31 |
| H3 (+dropout SimCSE) | r3 | 76.04 | 83.52 | 86.36 | 89.61 | 80.28 | 91.40 | 84.53 |
| H3 (+dropout SimCSE) | mean±std | 74.98±1.34 | 81.04±2.30 | 85.10±1.84 | 88.77±0.94 | 79.31±1.06 | 90.23±1.76 | 83.24±1.33 |
| H4 (+revcomp SimCSE) | r1 | 74.59 | 84.09 | 84.47 | 89.66 | 81.99 | 90.85 | 84.28 |
| H4 (+revcomp SimCSE) | r2 | 76.31 | 82.46 | 87.58 | 89.41 | 81.22 | 92.83 | 84.97 |
| H4 (+revcomp SimCSE) | r3 | 79.02 | 85.53 | 84.92 | 91.67 | 79.59 | 92.52 | 85.54 |
| H4 (+revcomp SimCSE) | mean±std | 76.64±2.23 | 84.03±1.54 | 85.66±1.68 | 90.25±1.24 | 80.93±1.23 | 92.07±1.07 | 84.93±0.63 |
| H5 (+crop SimCSE) | r1 | 86.13 | 90.98 | 88.90 | 92.62 | 89.71 | 94.43 | 90.46 |
| H5 (+crop SimCSE) | r2 | 84.43 | 88.63 | 91.07 | 93.24 | 88.79 | 94.01 | 90.03 |
| H5 (+crop SimCSE) | r3 | 87.48 | 91.66 | 91.77 | 93.52 | 89.00 | 96.05 | 91.58 |
| H5 (+crop SimCSE) | mean±std | 86.01±1.53 | 90.42±1.59 | 90.58±1.50 | 93.13±0.46 | 89.17±0.48 | 94.83±1.08 | 90.69±0.80 |
| H6 (+local-shift SimCSE) | r1 | 81.47 | 86.97 | 85.21 | 90.82 | 84.83 | 93.30 | 87.10 |
| H6 (+local-shift SimCSE) | r2 | 80.96 | 86.34 | 87.80 | 91.30 | 82.97 | 93.90 | 87.21 |
| H6 (+local-shift SimCSE) | r3 | 85.21 | 86.57 | 88.80 | 90.94 | 84.12 | 93.95 | 88.26 |
| H6 (+local-shift SimCSE) | mean±std | 82.55±2.32 | 86.63±0.32 | 87.27±1.85 | 91.02±0.25 | 83.97±0.94 | 93.72±0.36 | 87.52±0.64 |

Table S20: HyenaDNA Step 5 full fine-tuning GUE MCC (%) per seed, promoter and human TF-binding tasks.

| Model | Run | PC-All | PC-TATA | PC-noTA | P3-All | P3-TATA | P3-noTA | TF-H0 | TF-H1 | TF-H2 | TF-H3 | TF-H4 |
|---|---|---|---|---|---|---|---|---|---|---|---|---|
| H0 (HyenaDNA causal) | r1 | 61.12 | 71.57 | 67.24 | 85.48 | 58.40 | 92.11 | 58.96 | 54.06 | 52.37 | 44.05 | 74.13 |
| H0 (HyenaDNA causal) | r2 | 64.17 | 73.56 | 68.11 | 84.77 | 56.46 | 91.56 | 56.61 | 56.29 | 51.78 | 45.58 | 75.60 |
| H0 (HyenaDNA causal) | r3 | 63.86 | 70.99 | 67.37 | 83.79 | 57.56 | 91.97 | 56.13 | 55.80 | 53.19 | 44.12 | 73.67 |
| H0 (HyenaDNA causal) | mean±std | 63.05±1.68 | 72.04±1.35 | 67.57±0.47 | 84.68±0.85 | 57.47±0.97 | 91.88±0.29 | 57.23±1.51 | 55.38±1.17 | 52.45±0.71 | 44.58±0.86 | 74.47±1.01 |
| H1 (+bidir patch) | r1 | 65.18 | 56.29 | 68.26 | 86.42 | 59.12 | 92.01 | 57.04 | 54.22 | 51.61 | 41.17 | 76.01 |
| H1 (+bidir patch) | r2 | 63.41 | 64.16 | 67.69 | 84.64 | 57.75 | 92.76 | 56.23 | 55.60 | 52.19 | 42.39 | 73.39 |
| H1 (+bidir patch) | r3 | 63.46 | 51.48 | 67.42 | 86.39 | 59.40 | 92.28 | 57.35 | 53.82 | 52.27 | 43.96 | 70.65 |
| H1 (+bidir patch) | mean±std | 64.02±1.01 | 57.31±6.40 | 67.79±0.43 | 85.82±1.02 | 58.76±0.88 | 92.35±0.38 | 56.87±0.58 | 54.55±0.93 | 52.02±0.36 | 42.51±1.40 | 73.35±2.68 |
| H2 (+MNTP) | r1 | 64.64 | 71.94 | 69.10 | 86.89 | 63.02 | 92.95 | 58.91 | 57.29 | 53.97 | 44.53 | 75.41 |
| H2 (+MNTP) | r2 | 64.00 | 72.70 | 67.86 | 86.29 | 62.54 | 93.32 | 58.15 | 56.62 | 54.73 | 45.98 | 76.04 |
| H2 (+MNTP) | r3 | 63.44 | 72.93 | 68.86 | 86.68 | 59.43 | 92.80 | 59.45 | 56.60 | 56.46 | 44.60 | 80.02 |
| H2 (+MNTP) | mean±std | 64.03±0.60 | 72.52±0.52 | 68.61±0.66 | 86.62±0.30 | 61.66±1.95 | 93.02±0.27 | 58.84±0.65 | 56.84±0.39 | 55.05±1.28 | 45.04±0.82 | 77.16±2.50 |
| H3 (+dropout SimCSE) | r1 | 65.64 | 75.46 | 68.38 | 85.92 | 62.35 | 92.43 | 60.22 | 56.82 | 56.72 | 48.02 | 76.81 |
| H3 (+dropout SimCSE) | r2 | 66.17 | 74.29 | 68.34 | 86.37 | 59.42 | 92.95 | 56.02 | 57.68 | 55.67 | 45.23 | 75.00 |
| H3 (+dropout SimCSE) | r3 | 66.00 | 73.48 | 68.66 | 86.39 | 61.52 | 93.10 | 57.96 | 56.29 | 55.98 | 46.10 | 78.25 |
| H3 (+dropout SimCSE) | mean±std | 65.94±0.27 | 74.41±1.00 | 68.46±0.17 | 86.23±0.27 | 61.10±1.51 | 92.83±0.35 | 58.07±2.10 | 56.93±0.70 | 56.12±0.54 | 46.45±1.43 | 76.69±1.63 |
| H4 (+revcomp SimCSE) | r1 | 65.12 | 71.48 | 67.88 | 86.72 | 61.99 | 93.21 | 58.08 | 54.02 | 56.37 | 46.78 | 76.23 |
| H4 (+revcomp SimCSE) | r2 | 66.38 | 76.63 | 68.41 | 85.76 | 62.35 | 92.97 | 58.14 | 57.13 | 55.88 | 47.88 | 78.02 |
| H4 (+revcomp SimCSE) | r3 | 65.76 | 74.32 | 69.01 | 85.57 | 60.10 | 93.63 | 57.14 | 54.62 | 57.86 | 43.07 | 77.26 |
| H4 (+revcomp SimCSE) | mean±std | 65.75±0.63 | 74.14±2.58 | 68.43±0.57 | 86.02±0.62 | 61.48±1.21 | 93.27±0.33 | 57.79±0.56 | 55.26±1.65 | 56.70±1.03 | 45.91±2.52 | 77.17±0.90 |
| H5 (+crop SimCSE) | r1 | 65.51 | 64.48 | 68.74 | 86.70 | 58.90 | 92.56 | 56.58 | 55.22 | 52.18 | 45.89 | 75.93 |
| H5 (+crop SimCSE) | r2 | 64.09 | 68.03 | 68.33 | 85.14 | 55.56 | 93.13 | 55.79 | 55.94 | 51.71 | 44.29 | 76.63 |
| H5 (+crop SimCSE) | r3 | 64.16 | 72.02 | 69.30 | 86.17 | 58.12 | 92.60 | 59.12 | 56.47 | 56.80 | 46.63 | 77.60 |
| H5 (+crop SimCSE) | mean±std | 64.59±0.80 | 68.18±3.77 | 68.79±0.49 | 86.00±0.79 | 57.53±1.75 | 92.76±0.32 | 57.16±1.74 | 55.88±0.63 | 53.56±2.81 | 45.60±1.20 | 76.72±0.84 |
| H6 (+local-shift SimCSE) | r1 | 63.54 | 68.48 | 68.22 | 85.74 | 60.37 | 92.76 | 57.65 | 52.73 | 56.60 | 45.30 | 76.04 |
| H6 (+local-shift SimCSE) | r2 | 65.31 | 70.64 | 68.02 | 85.71 | 59.18 | 93.19 | 56.89 | 56.62 | 54.39 | 44.85 | 78.21 |
| H6 (+local-shift SimCSE) | r3 | 64.87 | 72.60 | 69.64 | 85.91 | 57.63 | 92.94 | 60.25 | 54.90 | 54.38 | 48.04 | 76.20 |
| H6 (+local-shift SimCSE) | mean±std | 64.57±0.92 | 70.57±2.06 | 68.63±0.88 | 85.79±0.11 | 59.06±1.37 | 92.96±0.22 | 58.26±1.76 | 54.75±1.95 | 55.12±1.28 | 46.06±1.73 | 76.82±1.21 |

Table S21: HyenaDNA Step 5 full fine-tuning GUE MCC (%) per seed, mouse TF-binding, splice, and Avg-GUE17.

| Model | Run | TF-M0 | TF-M1 | TF-M2 | TF-M3 | TF-M4 | Splice | Avg-GUE17 |
|---|---|---|---|---|---|---|---|---|
| H0 (HyenaDNA causal) | r1 | 54.30 | 80.25 | 75.37 | 65.62 | 43.98 | 79.01 | 65.77 |
| H0 (HyenaDNA causal) | r2 | 52.85 | 80.62 | 72.20 | 68.84 | 40.73 | 79.52 | 65.84 |
| H0 (HyenaDNA causal) | r3 | 52.85 | 80.33 | 66.46 | 61.43 | 43.91 | 81.47 | 64.99 |
| H0 (HyenaDNA causal) | mean±std | 53.33±0.84 | 80.40±0.19 | 71.34±4.52 | 65.30±3.72 | 42.87±1.86 | 80.00±1.30 | 65.53±0.47 |
| H1 (+bidir patch) | r1 | 49.39 | 81.32 | 80.00 | 61.32 | 43.60 | 83.04 | 65.06 |
| H1 (+bidir patch) | r2 | 47.01 | 81.95 | 74.44 | 54.84 | 44.72 | 82.87 | 64.47 |
| H1 (+bidir patch) | r3 | 48.14 | 81.83 | 78.70 | 53.47 | 42.61 | 85.09 | 64.02 |
| H1 (+bidir patch) | mean±std | 48.18±1.19 | 81.70±0.33 | 77.71±2.91 | 56.54±4.19 | 43.64±1.06 | 83.67±1.24 | 64.52±0.52 |
| H2 (+MNTP) | r1 | 56.55 | 81.75 | 85.47 | 71.15 | 43.92 | 85.77 | 68.43 |
| H2 (+MNTP) | r2 | 57.36 | 81.95 | 87.83 | 66.77 | 45.01 | 83.44 | 68.27 |
| H2 (+MNTP) | r3 | 58.07 | 81.35 | 85.99 | 73.27 | 46.29 | 83.89 | 68.83 |
| H2 (+MNTP) | mean±std | 57.33±0.76 | 81.68±0.31 | 86.43±1.24 | 70.40±3.31 | 45.07±1.19 | 84.37±1.24 | 68.51±0.29 |
| H3 (+dropout SimCSE) | r1 | 55.18 | 81.73 | 85.37 | 69.06 | 44.24 | 85.01 | 68.79 |
| H3 (+dropout SimCSE) | r2 | 53.65 | 82.48 | 82.93 | 66.77 | 46.99 | 82.12 | 67.77 |
| H3 (+dropout SimCSE) | r3 | 47.60 | 81.54 | 86.11 | 61.01 | 47.94 | 83.67 | 67.74 |
| H3 (+dropout SimCSE) | mean±std | 52.14±4.01 | 81.92±0.50 | 84.80±1.66 | 65.61±4.15 | 46.39±1.92 | 83.60±1.45 | 68.10±0.60 |
| H4 (+revcomp SimCSE) | r1 | 56.03 | 81.80 | 86.59 | 76.85 | 45.70 | 84.53 | 68.79 |
| H4 (+revcomp SimCSE) | r2 | 56.60 | 82.03 | 81.21 | 71.64 | 46.00 | 84.12 | 68.89 |
| H4 (+revcomp SimCSE) | r3 | 53.78 | 82.48 | 79.88 | 75.25 | 47.30 | 83.67 | 68.28 |
| H4 (+revcomp SimCSE) | mean±std | 55.47±1.49 | 82.10±0.35 | 82.56±3.55 | 74.58±2.67 | 46.33±0.85 | 84.11±0.43 | 68.65±0.33 |
| H5 (+crop SimCSE) | r1 | 47.43 | 81.24 | 60.98 | 29.73 | 44.30 | 84.32 | 62.98 |
| H5 (+crop SimCSE) | r2 | 48.46 | 81.75 | 65.85 | 31.70 | 41.21 | 68.39 | 62.12 |
| H5 (+crop SimCSE) | r3 | 42.87 | 81.75 | 62.81 | 35.58 | 44.58 | 68.31 | 63.23 |
| H5 (+crop SimCSE) | mean±std | 46.25±2.97 | 81.58±0.29 | 63.21±2.46 | 32.34±2.98 | 43.36±1.87 | 73.67±9.22 | 62.78±0.58 |
| H6 (+local-shift SimCSE) | r1 | 51.57 | 81.25 | 78.92 | 55.02 | 44.25 | 83.06 | 65.97 |
| H6 (+local-shift SimCSE) | r2 | 55.53 | 81.56 | 84.89 | 73.33 | 44.55 | 82.24 | 67.95 |
| H6 (+local-shift SimCSE) | r3 | 46.50 | 81.62 | 80.63 | 61.63 | 46.01 | 79.37 | 66.65 |
| H6 (+local-shift SimCSE) | mean±std | 51.20±4.53 | 81.48±0.20 | 81.48±3.07 | 63.33±9.27 | 44.94±0.94 | 81.56±1.94 | 66.86±1.00 |

Table S22: HyenaDNA Step 5 full fine-tuning Virus-COVID macro-F1 (%) per seed.

| Model | Run | Virus-COVID F1 |
|---|---|---|
| H0 (HyenaDNA causal) | r1 | 73.26 |
| H0 (HyenaDNA causal) | r2 | 72.99 |
| H0 (HyenaDNA causal) | r3 | 73.28 |
| H0 (HyenaDNA causal) | mean±std | 73.18±0.16 |
| H1 (+bidir patch) | r1 | 75.86 |
| H1 (+bidir patch) | r2 | 76.15 |
| H1 (+bidir patch) | r3 | 75.90 |
| H1 (+bidir patch) | mean±std | 75.97±0.16 |
| H2 (+MNTP) | r1 | 76.34 |
| H2 (+MNTP) | r2 | 76.11 |
| H2 (+MNTP) | r3 | 76.21 |
| H2 (+MNTP) | mean±std | 76.22±0.12 |
| H3 (+dropout SimCSE) | r1 | 75.74 |
| H3 (+dropout SimCSE) | r2 | 75.61 |
| H3 (+dropout SimCSE) | r3 | 75.12 |
| H3 (+dropout SimCSE) | mean±std | 75.49±0.33 |
| H4 (+revcomp SimCSE) | r1 | 75.22 |
| H4 (+revcomp SimCSE) | r2 | 76.09 |
| H4 (+revcomp SimCSE) | r3 | 73.88 |
| H4 (+revcomp SimCSE) | mean±std | 75.06±1.11 |
| H5 (+crop SimCSE) | r1 | 75.95 |
| H5 (+crop SimCSE) | r2 | 75.92 |
| H5 (+crop SimCSE) | r3 | 75.86 |
| H5 (+crop SimCSE) | mean±std | 75.91±0.05 |
| H6 (+local-shift SimCSE) | r1 | 73.24 |
| H6 (+local-shift SimCSE) | r2 | 72.04 |
| H6 (+local-shift SimCSE) | r3 | 73.16 |
| H6 (+local-shift SimCSE) | mean±std | 72.81±0.67 |

across-variant comparison: learning rate $3 \times 10^{-5}$, cosine schedule, warmup 50, gradient checkpointing, three seeds, and the released GUE_v2 EPI splits.

The pattern is not a continuation of the frozen H5 EPI ranking. Under full supervision every adapted variant falls below the causal H0 baseline on Avg-EPI MCC (Table S23): the frozen long-context EPI advantage does not survive task-specific fine-tuning, with crop (H5) staying closest to H0 and dropout (H3) regressing most. On the shorter GUE families the ordering reverses. The adapted initialisations mostly improve over H0 on the 17-task GUE average and on splice-site prediction, the exception being crop (H5), which is the only adapted variant below H0 on both GUE17 and splice. There is no splice collapse under the preserve-dtype pipeline; the earlier collapse was a `bfloat16` measurement artefact.

Table S23: HyenaDNA preserve-dtype Step 5 full fine-tuning family summary, mean over three seeds. GUE17 and Splice are MCC; Avg-EPI is MCC; Virus-COVID is macro-F1. Per-seed values are in Section S2.

| Model | GUE17 MCC | Splice MCC | Avg-EPI MCC | Virus-COVID F1 |
|---|---|---|---|---|
| H0 (causal) | 65.53 | 80.00 | **92.52** | 73.18 |
| H1 (+bidir patch) | 64.52 | 83.67 | 88.82 | 75.97 |
| H2 (+MNTP) | 68.51 | **84.37** | 89.66 | **76.22** |
| H3 (+dropout) | 68.10 | 83.60 | 83.24 | 75.49 |
| H4 (+revcomp) | **68.65** | 84.11 | 84.93 | 75.06 |
| H5 (+crop) | 62.78 | 73.67 | 90.69 | 75.91 |
| H6 (+local-shift) | 66.86 | 81.56 | 87.52 | 72.81 |

This diagnostic supports the main-text interpretation: full fine-tuning can override the frozen-representation ranking. The supervised EPI advantage does not transfer from the frozen setting, whereas the adapted initialisations help on the short GUE families—most clearly for MNTP (H2)—which is why the main task-conditioned claims rest on the Step 4 frozen linear probe.

### S4 Task-Family Paired Statistical Checks

To accompany the task-family matrix in the main paper, we report selected paired task-level Wilcoxon signed-rank checks across biologically matched tasks. These tests are intended as compact support for the main qualitative regimes rather than as exhaustive multiple-testing-corrected inference across all model pairs. The generated table `task_family_paired_tests.tex` was not found. It can be regenerated with `uv run python src/analyze_task_family_paired_tests.py`.

### S5 Weighted-Mean Pooling Ablation

The main Step 4 results use simple mean pooling over token-level hidden states. As a robustness check, we also evaluated the adapted mask-only checkpoints with a weighted-mean pooling alternative, for both backbones (Tables S24 and S25). The EPI pattern is unchanged: crop contrastive learning (M5/H5) remains the strongest variant on long-range EPI under weighted pooling, matching the main mean-pooling results (DNAGPT 29.05, HyenaDNA 28.75 Avg-EPI MCC). Shorter-sequence averages are more pooling-sensitive but preserve the same qualitative ordering.

Table S24: Step 4 weighted-mean pooling ablation (DNAGPT), mask-only setting. Mean ± std over three seeds. GB accuracy is over all 8 GB tasks; NT, GUE, and EPI use MCC; GUE is averaged over the 17 non-COVID GUE tasks; Virus-COVID is macro-F1.

| Model | GB acc | NT MCC | GUE MCC | COVID F1 | Avg-EPI MCC |
|---|---|---|---|---|---|
| M0 (causal) | 78.18±0.00 | 47.22±0.00 | **51.74**±0.00 | 26.97±0.00 | 13.35±0.00 |
| M1 (+bidir attn) | 78.26±0.00 | 44.64±0.00 | 49.59±0.00 | 11.80±0.00 | 14.03±0.00 |
| M2 (+MNTP) | 78.07±0.16 | 44.89±0.56 | 49.28±0.45 | 30.40±1.84 | 20.02±0.20 |
| M3 (+dropout SimCSE) | 78.12±0.02 | 45.58±0.52 | 50.74±0.22 | 27.05±0.81 | 23.57±0.32 |
| M4 (+RC SimCSE) | **78.92**±0.03 | **47.83**±0.23 | 51.20±0.29 | 29.88±1.06 | 21.77±0.27 |
| M5 (+crop SimCSE) | 77.98±0.13 | 46.13±0.15 | 50.56±0.26 | **42.55**±0.45 | **29.05**±0.51 |
| M6 (+local-shift) | 78.25±0.03 | 46.52±0.42 | 50.95±0.61 | 38.48±2.03 | 25.06±0.77 |

Table S25: Step 4 weighted-mean pooling ablation (HyenaDNA), same protocol and columns as Table S24. Mean ± std over three seeds; H0 and H1 are deterministic.

| Model | GB acc | NT MCC | GUE MCC | COVID F1 | Avg-EPI MCC |
|---|---|---|---|---|---|
| H0 (causal) | 77.23±0.00 | 37.05±0.00 | 42.53±0.00 | 8.33±0.00 | 16.62±0.00 |
| H1 (+bidir patch) | 74.55±0.00 | 37.01±0.00 | 38.10±0.00 | 11.06±0.00 | 12.45±0.00 |
| H2 (+MNTP) | 76.58±0.18 | 35.91±0.20 | 40.48±0.30 | 11.13±0.52 | 16.90±0.54 |
| H3 (+dropout SimCSE) | 78.48±0.62 | 41.24±0.68 | 42.50±0.82 | 17.40±0.09 | 23.46±0.61 |
| H4 (+RC SimCSE) | 78.82±0.23 | 43.12±1.27 | 43.39±0.92 | 15.84±0.73 | 23.08±0.75 |
| H5 (+crop SimCSE) | 76.38±0.16 | 42.79±0.46 | 42.60±0.71 | **30.62**±0.99 | **28.75**±0.22 |
| H6 (+local-shift) | **78.95**±0.23 | **44.82**±0.26 | **46.59**±1.08 | 20.87±0.19 | 25.61±1.58 |

### S6 Embedding-Table Update Policy: Mask-Only vs. Full-Embedding

The Stage 2 MNTP design has one degree of freedom that is invisible to the LLM2Vec formulation but unavoidable once the mask token is a newly added vocabulary entry: *which rows of the token embedding matrix should be allowed to change during MNTP training?* The original LLM2Vec paper does not face this question because it reuses an existing token (the underscore `_`) as the mask placeholder and freezes the entire embedding matrix. For DNAGPT no equivalent semantically neutral token exists in the BPE vocabulary, so we add a dedicated `[MASK]` token whose embedding row must be learned during Stage 2. We evaluated two policies for which rows of the embedding matrix are updated during this learning:

**Mask-only (used in the main paper).** Every gradient applied to the `wte` matrix has all rows except the `[MASK]` row zeroed via a backward hook, and the matrix is placed in a `weight_decay`=0 optimiser group so AdamW's decoupled decay cannot drift the frozen rows. Effective trainable parameters added by this policy: 768 (one row).

**Full-embedding (*maskfix* below).** The full `wte` matrix (25,001 rows × 768, after adding `[MASK]`) is added to the set of trainable parameters via PEFT's `modules_to_save` mechanism, which keeps specified non-LoRA modules trainable and saves them alongside the adapter weights. Under this policy, the entire embedding table is updated during MNTP training. Effective trainable parameters added by this policy: ∼38.6 M (≈37% of the 105 M backbone), in addition to the LoRA adapters on attention projections.

**Why mask-only is the main-paper default.** Mask-only is the conservative reading of the LLM2Vec recipe: it keeps the M0→M2 delta attributable to bidirectional LoRA adaptation plus a single learned mask embedding, rather than a global re-tuning of all 25,000 pretrained BPE embeddings ($\sim$38.6 M extra parameters, $\approx$37% of the backbone). We report full-embedding numbers only for transparency; they should be read as a deliberate departure from LLM2Vec rather than as the default.

**Quantitative comparison.** Table S26 reports the full Step 4 linear probe results under both policies, on the same M0–M6 model variants, the same training data (hg38 with N-filtering), and the same evaluation protocol. M2–M6 are mean $\pm$ std over three independent seeds (42, 43, 44).

Table S26: Step 4 linear probe results (mean pooling) under the two embedding-table update policies. GB: accuracy (%); NT and EPI: MCC (%); GUE: MCC (%) averaged over the 17 non-COVID GUE tasks. M2–M6 are mean $\pm$ std over 3 seeds (42, 43, 44). M0 and M1 are deterministic and identical across the two policies (no embedding-table update is performed in either). Bold values within a row indicate the better policy for that (model, benchmark) cell.

| Model | Policy | GB acc | NT MCC | GUE MCC | Avg-EPI MCC |
|---|---|---|---|---|---|
| M0 (DNAGPT causal) | — | 78.08 | 45.99 | 50.80 | 13.42 |
| M1 (+bidir attn) | — | 78.34 | 44.50 | 49.14 | 13.96 |
| M2 (+MNTP LoRA) | maskfix | 78.19$\pm$0.10 | **45.78**$\pm$0.16 | 49.41$\pm$0.12 | **19.84**$\pm$0.37 |
| | mask-only | **78.26**$\pm$0.11 | 45.75$\pm$0.19 | **49.67**$\pm$0.55 | 19.70$\pm$0.57 |
| M3 (+dropout SimCSE) | maskfix | **78.49**$\pm$0.09 | **46.42**$\pm$0.33 | 50.51$\pm$0.38 | 23.01$\pm$0.15 |
| | mask-only | 78.44$\pm$0.03 | 46.29$\pm$0.30 | **50.72**$\pm$0.15 | **23.22**$\pm$0.26 |
| M4 (+revcomp SimCSE) | maskfix | 78.81$\pm$0.08 | 46.81$\pm$0.33 | **50.79**$\pm$0.32 | **21.19**$\pm$0.31 |
| | mask-only | **78.96**$\pm$0.13 | **47.11**$\pm$0.30 | 50.74$\pm$0.14 | 21.01$\pm$0.12 |
| M5 (+crop SimCSE) | maskfix | **78.14**$\pm$0.14 | **45.89**$\pm$0.17 | **50.00**$\pm$0.45 | **27.91**$\pm$0.63 |
| | mask-only | 78.13$\pm$0.10 | 45.74$\pm$0.51 | 49.88$\pm$0.32 | 27.46$\pm$0.20 |
| M6 (+local-shift SimCSE) | maskfix | 78.50$\pm$0.17 | 46.14$\pm$0.18 | 50.36$\pm$0.50 | 24.28$\pm$0.90 |
| | mask-only | **78.54**$\pm$0.10 | **46.15**$\pm$0.39 | **50.50**$\pm$0.36 | **24.34**$\pm$1.42 |

M6 maskfix and mask-only rows both report mean $\pm$ std over three seeds.

**Reading the comparison.** Across the 20 model$\times$benchmark cells (M2–M6) the two policies split evenly (10/20 each) and differ by at most $\sim$0.5 pp; on the headline EPI metric, M5 scores $27.91 \pm 0.63$ % (maskfix) versus $27.46 \pm 0.20$ % (mask-only), a 0.45 pp gap with overlapping intervals. The embedding-update policy therefore does not materially change the adaptation picture—including for M6, now complete under both policies—and the small-gain (short-sequence) versus large-gain (long-range) pattern is preserved under both.

### S7 MNTP Checkpoint-Convergence Diagnostic

We evaluated M2 checkpoints saved every 1000 steps during a 10 000-step MNTP run as a single-seed checkpoint-convergence diagnostic (r1). The purpose of this sweep is to assess whether the masked adaptation stage has a single uniformly optimal training duration across benchmark families.

GB is essentially flat, NT reaches a broad plateau after 4 000–7 000 steps, GUE improves slowly, COVID F1 peaks at 3 000 steps, and Avg-EPI MCC also peaks early and remains within 0.85 pp thereafter. The default 5 757-step M2 checkpoint therefore provides a practical compromise across task types rather than a family-specific optimum.

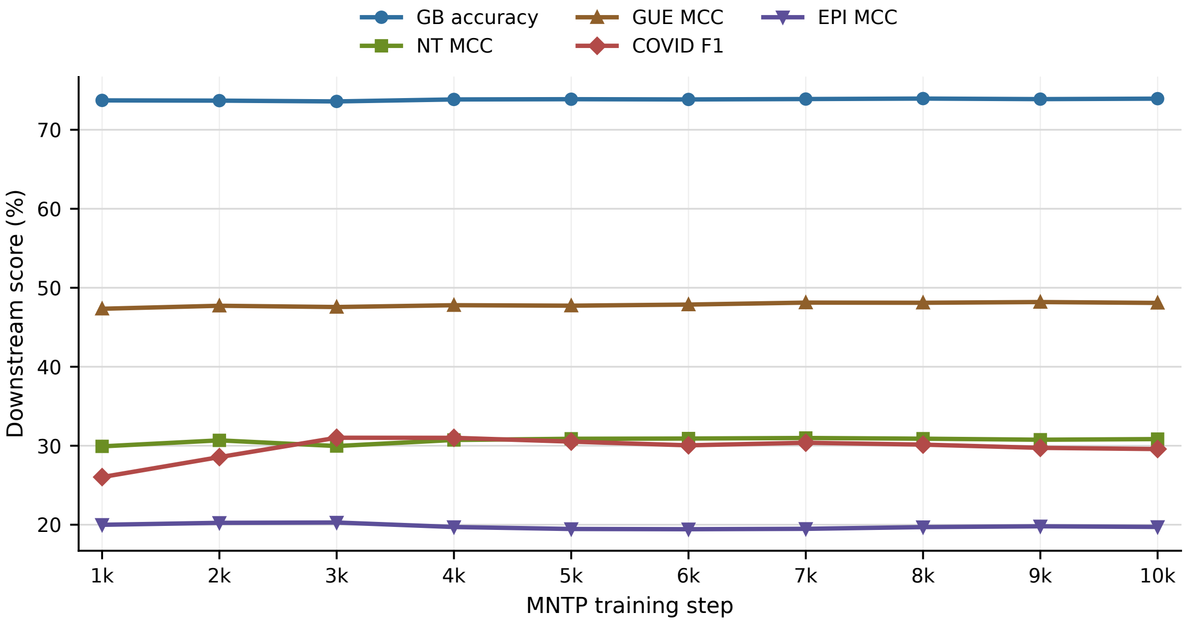


Figure S1: Downstream performance of M2 checkpoints at 1 000-step intervals during MNTP training under the mean-pooling linear-probe protocol. This is a single-seed checkpoint-convergence diagnostic (r1). Short-sequence benchmarks change modestly across the sweep, whereas COVID F1 and Avg-EPI MCC peak early and then remain close to their best values.

### S8 EPI Pair-Representation Geometry

To complement the linear-probe EPI results, we evaluated the geometry of frozen enhancer–promoter pair embeddings. For each EPI cell-line task, complete enhancer–promoter pairs were embedded using the same junction-cropped input representation as the main EPI analyses. We then measured whether interacting and non-interacting pairs were separable under two embedding-space diagnostics: (i) prototype scoring, using the relative similarity to positive and negative class centroids, and (ii) local $k$-nearest-neighbour (kNN) scoring in the pair-embedding space. We also report a pair-ranking diagnostic, where each positive pair is ranked against sampled negative pairs.

Table S27: EPI pair-representation geometry (DNAGPT), averaged across the six EPI cell-line tasks. Prototype metrics measure global positive-vs-negative centroid separation. kNN metrics measure local neighbourhood separability of interacting and non-interacting enhancer–promoter pairs. Rank Top10 is the percentage of positive pairs ranked in the top 10 among sampled candidate pairs.

| Model | Proto AUC | Proto MCC | kNN AUC | kNN MCC | Rank Top10 |
|---|---|---|---|---|---|
| M0 (causal) | 0.559 | 0.095 | 0.686 | 0.264 | 13.57 |
| M1 (+bidir attn) | 0.548 | 0.074 | 0.662 | 0.239 | 13.90 |
| M2 (+MNTP) | **0.603** | 0.152 | 0.734 | 0.336 | **17.66** |
| M3 (+dropout SimCSE) | 0.595 | 0.130 | 0.796 | 0.422 | 17.29 |
| M4 (+RC SimCSE) | 0.599 | **0.152** | 0.740 | 0.344 | **17.66** |
| M5 (+crop SimCSE) | 0.600 | 0.135 | **0.855** | **0.498** | 17.57 |
| M6 (+local-shift) | 0.588 | 0.130 | 0.811 | 0.446 | 16.78 |

The local-neighbourhood (kNN) metrics show the clearest separation, and in both backbones the crop-contrastive variant is strongest there. For DNAGPT, M5 attains the highest kNN AUC and kNN MCC on every individual EPI cell-line task, raising the

Table S28: EPI pair-representation geometry (HyenaDNA), same protocol and columns as Table S27. H2–H6 are single-seed (r1); H0 and H1 are deterministic.

| Model | Proto AUC | Proto MCC | kNN AUC | kNN MCC | Rank Top10 |
|---|---|---|---|---|---|
| H0 (causal) | 0.600 | 0.148 | 0.748 | 0.349 | 17.30 |
| H1 (+bidir patch) | 0.574 | 0.120 | 0.770 | 0.393 | 15.68 |
| H2 (+MNTP) | 0.591 | 0.141 | 0.752 | 0.359 | 16.31 |
| H3 (+dropout SimCSE) | 0.587 | 0.123 | 0.713 | 0.312 | 15.48 |
| H4 (+RC SimCSE) | 0.595 | 0.137 | 0.740 | 0.349 | 17.57 |
| H5 (+crop SimCSE) | **0.614** | **0.167** | **0.836** | **0.501** | **19.19** |
| H6 (+local-shift) | 0.597 | 0.143 | 0.799 | 0.434 | 17.90 |

average kNN AUC from 0.686 (M0) to 0.855 and kNN MCC from 0.264 to 0.498. For HyenaDNA, H5 is likewise the strongest variant, raising kNN AUC from 0.748 (H0) to 0.836 and kNN MCC from 0.349 to 0.501, and it also leads on the prototype and ranking diagnostics. In both backbones the prototype (global-centroid) metrics improve only modestly relative to the local kNN gains, consistent with enhancer–promoter interaction being a heterogeneous pair relation: the contrastive crop objective most strongly improves local pair-neighbourhood structure rather than global cluster separation.

### S9 Additional Staged-Pipeline Control Notes

The main paper's staged-pipeline ablation shows that Stage 3 crop contrastive learning is the dominant driver of the frozen EPI gain in both backbones (DNAGPT: Avg-EPI MCC 13.42 → 27.46 from the causal M0 to the staged M5; HyenaDNA: 15.88 → 26.65 from H0 to H5). We add three observations that clarify what the earlier stages contribute. Tables S29 and S30 give the full per-family breakdown of the frozen staged pipeline (M0–M6 / H0–H6), which the main text summarises only as Fam-mean.

Table S29: DNAGPT: frozen linear-probe performance per task family across the staged pipeline (M0–M6, mean-pooling, mask-only Stage 2 setting). Each family is reported in its native metric (accuracy for Genomics-Benchmarks tasks, MCC for Nucleotide-Transformer/GUE tasks, macro-F1 for Virus-COVID). Fam-mean is the descriptive mean over the seven families and mixes metrics, as in the main text. M2–M6 are the three-seed mean; M0/M1 are deterministic. Bold marks the best value per column.

| Model | EPI | COVID | Histone | Local-reg | TF | Promoter | Splice | Fam-mean |
|---|---|---|---|---|---|---|---|---|
| M0 (causal) | 13.42 | 27.40 | 39.50 | 62.92 | 47.21 | 70.38 | **30.30** | 41.59 |
| M1 (+bidir attn) | 13.96 | 11.81 | 38.57 | 62.35 | 46.32 | 69.27 | 25.34 | 38.23 |
| M2 (+MNTP) | 19.70 | 32.10 | 39.93 | 62.60 | 46.66 | 69.15 | 28.77 | 42.70 |
| M3 (+dropout SimCSE) | 23.22 | 28.85 | 40.56 | 62.41 | 47.73 | 70.40 | 28.86 | 43.15 |
| M4 (+RC SimCSE) | 21.01 | 30.28 | **41.81** | **63.32** | **47.78** | **70.71** | 28.00 | 43.27 |
| M5 (+crop SimCSE) | **27.46** | **42.63** | 40.02 | 62.31 | 46.81 | 69.72 | 28.02 | **45.28** |
| M6 (+local-shift) | 24.34 | 39.93 | 40.55 | 62.54 | 47.68 | 70.12 | 28.41 | 44.79 |

**Crop already works on the causal backbone.** Applying crop directly to the causal checkpoint (Stage 0 branch) recovers almost all of the EPI gain in both models: DNAGPT reaches 25.93 of its staged 27.46, and HyenaDNA reaches 26.57 of its staged 26.65. The staged Stage 1/2 initialisation adds a modest Fam-mean improvement in both backbones (43.91 → 45.28 for DNAGPT; 38.80 → 40.55 for HyenaDNA), plus a small further EPI bump for DNAGPT (+1.53 pp) but essentially none for HyenaDNA (+0.08 pp). The crop EPI effect is therefore robust to where adaptation branches, while the earlier stages mainly broaden multi-family competence.

Table S30: HyenaDNA: frozen linear-probe performance per task family across the staged pipeline (H0–H6), same protocol, metrics, and column definitions as Table S29. H2–H6 are the three-seed mean; H0/H1 are deterministic. Bold marks the best value per column.

| Model | EPI | COVID | Histone | Local-reg | TF | Promoter | Splice | Fam-mean |
|---|---|---|---|---|---|---|---|---|
| H0 (causal) | 15.88 | 11.33 | 28.08 | 63.25 | 38.54 | 59.20 | 17.36 | 33.38 |
| H1 (+bidir patch) | 11.39 | 11.32 | 33.16 | 62.46 | 34.12 | 58.93 | 14.53 | 32.27 |
| H2 (+MNTP) | 16.38 | 11.62 | 31.02 | 62.28 | 38.52 | 59.96 | 17.51 | 33.90 |
| H3 (+dropout SimCSE) | 22.40 | 17.36 | 36.37 | 64.67 | 36.66 | 64.55 | 24.11 | 38.02 |
| H4 (+RC SimCSE) | 21.12 | 15.76 | 39.05 | **65.16** | 38.66 | 63.91 | 23.08 | 38.10 |
| H5 (+crop SimCSE) | **26.65** | **31.48** | 38.52 | 62.56 | 35.99 | 65.63 | 23.00 | 40.55 |
| H6 (+local-shift) | 24.04 | 21.09 | **40.19** | 64.10 | **42.22** | **67.28** | **27.64** | **40.94** |

**Pretraining matters more for HyenaDNA's EPI signal.** Re-running the adaptation from a randomly initialised backbone leaves DNAGPT's crop EPI close to its pretrained value (23.39 vs. 27.46) but collapses HyenaDNA's (15.24 vs. 26.65, from a random causal baseline of only 6.88). Broad Fam-mean competence requires pretraining in both backbones (45.28 → 32.90 for DNAGPT; 40.94 → 36.20 for HyenaDNA).

**MNTP repairs out-of-domain embedding geometry.** The Stage 2 MNTP objective raises the DGEB-DNA mean primary metric in both backbones, more strongly for DNAGPT (0.164 → 0.268, M0→M2) than for HyenaDNA (0.174 → 0.204, H0→H2; Tables S42 and S43).

Together these controls support the main-text interpretation: crop contrastive learning drives the largest long-range gain, while the earlier stages stabilise representation geometry and broaden transfer beyond the specific crop objective. A further EPI order-swap diagnostic is reported in Section S12.

### S10 Value of Pretraining: Full Per-Family Breakdown

The main text summarises the value-of-pretraining ablation as EPI and Fam-mean only. Tables S31 and S32 give the full per-family breakdown for the *randomly initialised* backbone, using the same native metrics, family definitions, and column layout as Tables S29 and S30 (the pretrained backbone), so the two can be compared directly. Without pretraining, every family drops and the best Fam-mean falls from 45.28 to 32.90 for DNAGPT and from 40.94 to 36.20 for HyenaDNA. The drop is far larger for HyenaDNA's EPI (best 16.39 vs. pretrained 26.65) than for DNAGPT's (crop M5 still reaches 23.39 vs. pretrained 27.46), consistent with the architecture-dependent role of pretraining reported in the main text.

Table S31: DNAGPT, *random initialisation*: frozen linear-probe performance per task family for the Stage-2/3 adaptation applied to a randomly initialised backbone (single seed, r1). Same native metrics and column definitions as Table S29. Bold marks the best value per column.

| Model | EPI | COVID | Histone | Local-reg | TF | Promoter | Splice | Fam-mean |
|---|---|---|---|---|---|---|---|---|
| M0 (causal) | 14.99 | 28.21 | 9.92 | 47.08 | 21.39 | 31.83 | 8.19 | 23.09 |
| M1 (+bidir attn) | 23.00 | 43.51 | 15.71 | 51.21 | 26.64 | 39.54 | 11.72 | 30.19 |
| M2 (+MNTP) | 8.34 | 9.02 | 10.61 | 54.70 | 5.53 | 30.98 | 3.95 | 17.59 |
| M3 (+dropout SimCSE) | **25.14** | **45.39** | 17.03 | 54.99 | 28.05 | 40.05 | 12.37 | 31.86 |
| M4 (+RC SimCSE) | 22.43 | 39.92 | **20.35** | **58.11** | **28.32** | **48.01** | **13.16** | **32.90** |
| M5 (+crop SimCSE) | 23.39 | 44.39 | 16.17 | 53.67 | 26.24 | 40.01 | 12.50 | 30.91 |
| M6 (+local-shift) | 23.29 | 43.41 | 16.05 | 51.87 | 26.81 | 39.53 | 12.20 | 30.45 |

Table S32: HyenaDNA, *random initialisation*: frozen linear-probe performance per task family, same protocol, metrics, and column definitions as Table S31. Bold marks the best value per column.

| Model | EPI | COVID | Histone | Local-reg | TF | Promoter | Splice | Fam-mean |
|---|---|---|---|---|---|---|---|---|
| H0 (causal) | 6.88 | 2.28 | 17.25 | 61.46 | 23.49 | 50.87 | 10.48 | 24.67 |
| H1 (+bidir patch) | 9.03 | 2.28 | 18.85 | 62.32 | 23.82 | 46.74 | 4.97 | 24.00 |
| H2 (+MNTP) | **16.39** | 11.17 | **35.68** | **64.61** | **43.62** | **63.33** | **18.58** | **36.20** |
| H3 (+dropout SimCSE) | 14.85 | **13.23** | 21.48 | 62.56 | 14.03 | 52.06 | 11.44 | 27.09 |
| H4 (+RC SimCSE) | 15.42 | 12.56 | 33.14 | 62.92 | 23.86 | 59.02 | 11.11 | 31.15 |
| H5 (+crop SimCSE) | 15.24 | 12.20 | 26.11 | 64.29 | 20.50 | 56.30 | 13.81 | 29.78 |
| H6 (+local-shift) | 13.38 | 12.93 | 23.01 | 63.15 | 19.66 | 54.48 | 10.89 | 28.21 |

### S11 EPI Input-Length Sweep

As a further control that the EPI gain tracks usable long-range context, we swept the visible input length for DNAGPT. When EPI inputs are centre-cropped to 512 bp the M5–M0 margin nearly vanishes (+0.30 pp); restoring the 4096 bp junction crop returns it to +14.12 pp (Fig. S2).

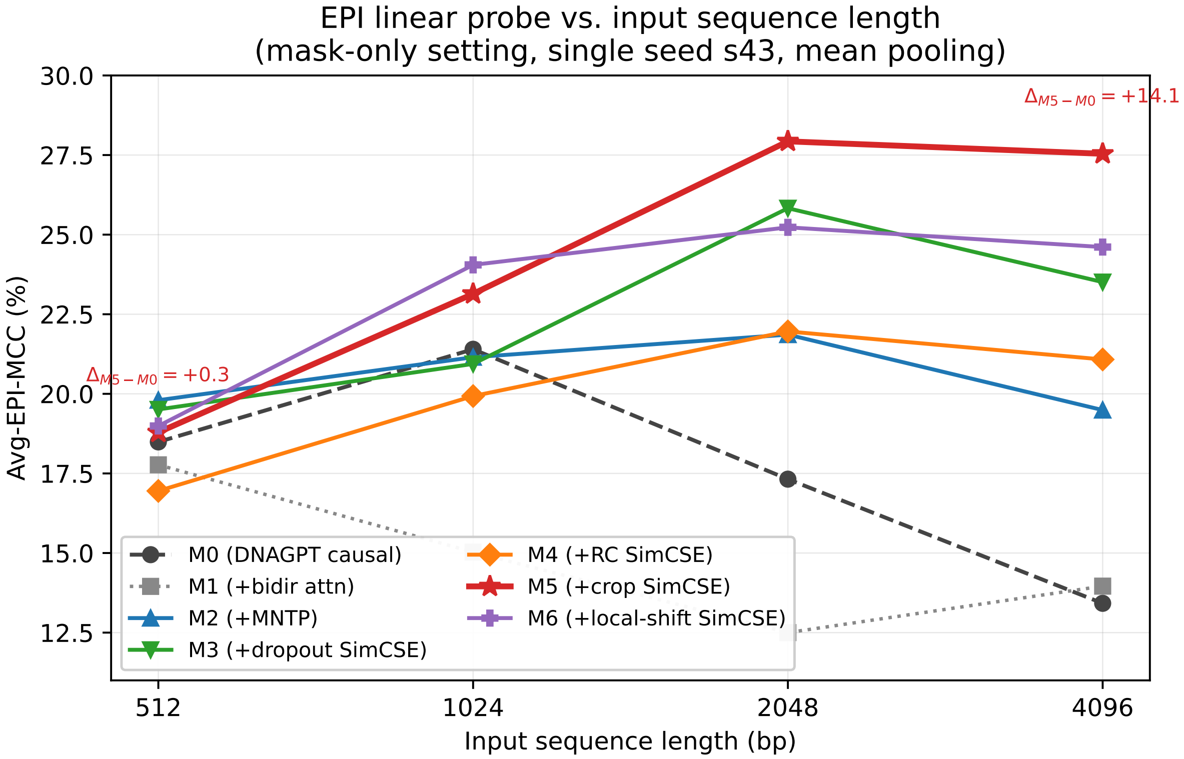


Figure S2: EPI linear probe performance (Avg-EPI MCC, mean pooling) as a function of input sequence length for all seven DNAGPT variants under the mask-only Stage 2 setting. Single-seed diagnostic (s43). The M5–M0 margin is near zero at 512 bp and grows as longer cross-junction context is restored.

### S12 EPI Enhancer–Promoter Order-Swap Diagnostic

To test whether the EPI gain is sensitive to the arbitrary concatenation order of enhancer and promoter sequences, we reversed the default input order from *enhancer + promoter* to *promoter* + *enhancer* and re-evaluated all DNAGPT and HyenaDNA variants under the same frozen linear-probe protocol. This is a single-seed diagnostic (s43) using the mask-only Stage 2 setting.

Table S33: DNAGPT: Avg-EPI MCC (%) under the original and reversed concatenation order. $\Delta$ = reversed − original.

| Model | Enh+Prom | Prom+Enh | $\Delta$ |
|---|---|---|---|
| M0 (causal) | 13.42 | 20.60 | **+7.18** |
| M1 (+bidir attn) | 13.96 | 13.77 | −0.19 |
| M2 (+MNTP) | 19.49 | 21.11 | +1.62 |
| M3 (+dropout SimCSE) | 23.51 | 25.00 | +1.49 |
| M4 (+RC SimCSE) | 21.08 | 20.56 | −0.52 |
| M5 (+crop SimCSE) | 27.54 | 28.06 | **+0.52** |
| M6 (+local-shift) | 24.61 | 25.37 | +0.76 |

Table S34: HyenaDNA: Avg-EPI MCC (%) under the original and reversed concatenation order, same protocol as Table S33. $\Delta$ = reversed − original. Single-seed (r1); H0 and H1 are deterministic.

| Model | Enh+Prom | Prom+Enh | $\Delta$ |
|---|---|---|---|
| H0 (causal) | 15.88 | 15.98 | +0.10 |
| H1 (+bidir patch) | 11.39 | 11.30 | −0.09 |
| H2 (+MNTP) | 16.27 | 16.32 | +0.05 |
| H3 (+dropout SimCSE) | 22.47 | 22.89 | +0.42 |
| H4 (+RC SimCSE) | 21.95 | 22.00 | +0.05 |
| H5 (+crop SimCSE) | 26.98 | 28.03 | +1.05 |
| H6 (+local-shift) | 24.74 | 23.62 | −1.12 |

For DNAGPT, M0 is strongly affected by concatenation order, whereas M5 remains nearly unchanged and stays above M0 even under M0's more favourable ordering, supporting the interpretation that the adapted bidirectional representation is less dependent on arbitrary enhancer–promoter ordering. HyenaDNA shows only small shifts throughout (all $|\Delta| \leq 1.1$ pp). Despite being a causal backbone, HyenaDNA H0 exhibits negligible change under enhancer–promoter order swapping when evaluated with mean-pooled sequence representations. This is not a contradiction with causality: the causal constraint restricts each token's hidden state to its left context, but averaging over all tokens at the pooling step largely washes out block-order and positional differences, so a causal model can still yield an approximately order-invariant *pooled* embedding. The size of any residual effect is backbone-dependent (large for DNAGPT M0, negligible for HyenaDNA H0). We therefore do not read H0's stability as bidirectionality or strict order-invariance, and keep the controlled claim to the within-DNAGPT reduction in order-sensitivity from M0 to the adapted variants.

### S13 Reflection-Equivariance Diagnostic (Full Table)

Table S35 gives the full reflection-equivariance diagnostic for the bidirectional HyenaDNA variants summarised in the main text. Per variant on held-out hg38 windows we measure: **tok_eq**, the per-position cosine between the reversed forward representation $R\,H(\mathbf{x})$ and the representation of the reversed input $H(R\mathbf{x})$ before pooling; **pooled_refl**$=\cos(f(\mathbf{x}), f(R\mathbf{x}))$ on the mean-pooled embedding; and **anchor**, the mean cosine between two distinct window embeddings (a measure of anisotropy). For all three, a lower value is the informative direction.

Table S35: Reflection-equivariance diagnostic for the bidirectional HyenaDNA variants (mean ± std over 3 seeds). tok_eq ≈ 1: token-level reflection equivariance; pooled_refl ≈ 1: orientation-insensitive pooled embedding; anchor ≈ 1: anisotropic (collapsed) embedding space.

| Variant | tok_eq | pooled_refl | anchor |
|---|---|---|---|
| H0 (causal) | 0.86±0.00 | 0.99±0.00 | 0.99±0.00 |
| H1 (bidir. patch) | 1.00±0.00 | 1.00±0.00 | 0.99±0.00 |
| H2 (MNTP) | 0.74±0.00 | 0.98±0.01 | 0.95±0.01 |
| H3 (dropout) | 0.11±0.05 | 0.42±0.01 | 0.29±0.03 |
| H4 (reverse-complement) | 0.59±0.05 | 0.71±0.05 | 0.38±0.03 |
| H5 (crop) | 0.19±0.01 | 0.58±0.03 | 0.45±0.03 |
| H6 (local-shift) | 0.15±0.01 | 0.50±0.02 | 0.39±0.02 |

### S14 Cross-Architecture Motif-Contact Effect Sizes

Figure S3 reports the mean motif-pair enrichment effect size across variants for both backbones, complementing the pooled Stouffer $z$ in the main text; effect sizes track the pooled $z$.

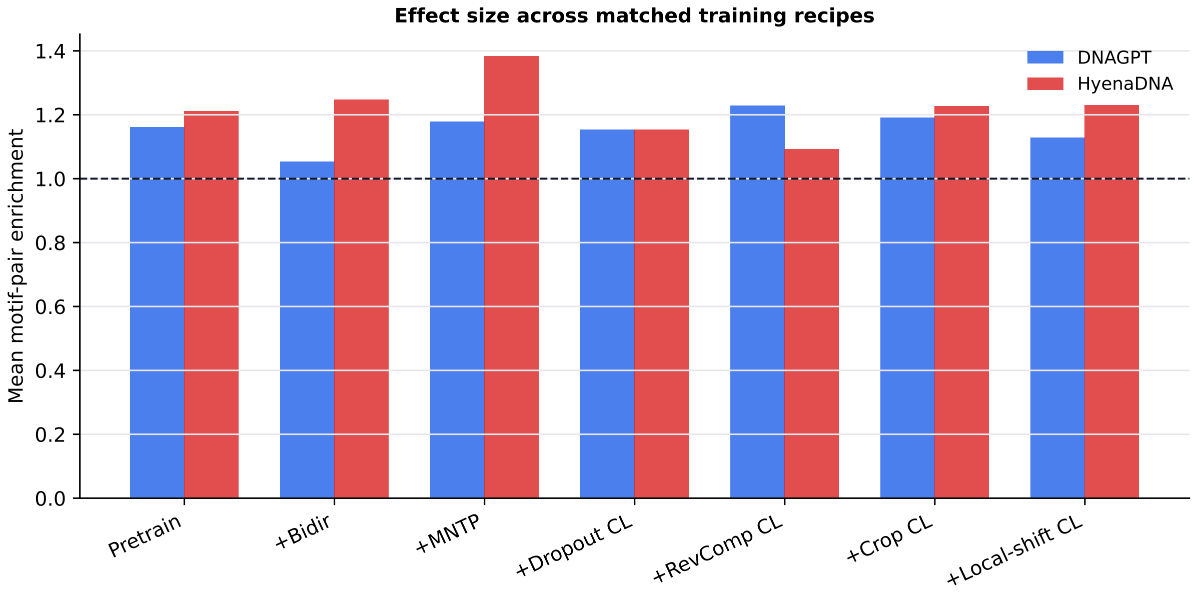


Figure S3: Cross-architecture motif-pair enrichment effect size (dashed line: distance-matched background). Effect sizes track the pooled Stouffer $z$ reported in the main text.

### S15 Full M0–M6 Attention-Map Case Study on EPI-GM12878

This section extends the main-paper attention-map case study from the two-model M0 vs. M5 comparison to the complete M0–M6 series on the same EPI-GM12878 example, junction-cropped to 4096 bp, under the mask-only embedding-update policy. The same positive (interacting) and negative (non-interacting) enhancer–promoter pairs are used. For every model variant we plot the head-averaged last-layer attention map, with a robust 99.5th-percentile colour cap applied per panel so that one sharp hotspot does not drive the rest of the panel to black.

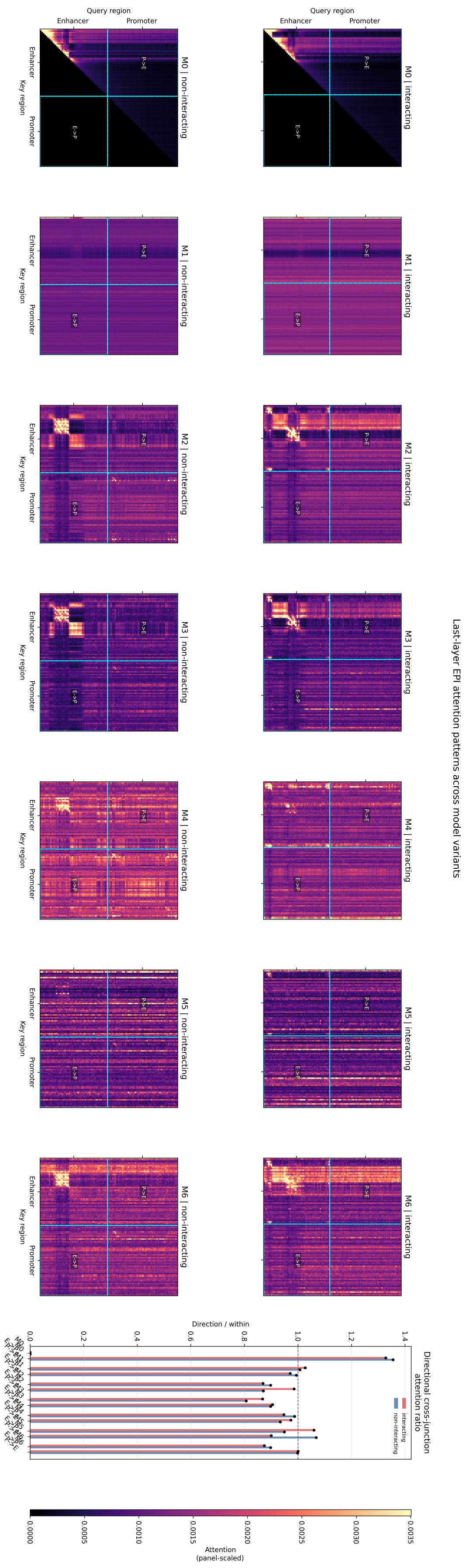


Figure S4: Last-layer attention maps (mean over heads) for all seven model variants (M0–M6) on one EPI-GM12878 example, junction-cropped to 4096 bp. Top row: positive (interacting) pair; bottom row: negative (non-interacting) pair. This panel extends the M0-vs.-M5 case study in the main paper.

**Reading the panel.** Three patterns are worth highlighting. First, the M0 vs. M1 comparison directly visualises why the bidirectional patch alone is insufficient: removing the causal mask on top of weights that were never trained for bidirectional attention produces noisy, untargeted attention. Second, M2 (MNTP-recovered) already restores most of the structure, with attention concentrating on regions that match short DNA motifs near the junction. Third, the contrastive variants M3–M6 differ qualitatively in how much cross-junction off-diagonal attention they produce: M5 (crop) shows the most cross-junction mass on positive examples, M4 (RC) shows pronounced attention along the diagonal mirror line consistent with its reverse-complement training objective, and M6 (local-shift) sits between M3 and M5. These qualitative differences are consistent with the relative ordering of the Avg-EPI MCC numbers in the main table.

### S16 DNAGPT Reverse-Complement Contact-Map Consistency

To test whether reverse-complement contrastive adaptation produces strand-consistent perturbation geometry, we recomputed each DNAGPT base-pair mutagenesis contact map on the reverse-complemented sequence, flipped the reverse-complement map back to forward coordinates, and measured Pearson correlation between the aligned maps. All models were evaluated on the same 24 hg38 windows and 4096 aligned positions per window, with the DNAGPT checkpoints loaded in float32.

Table S36: DNAGPT reverse-complement consistency of base-pair mutagenesis contact maps.

| **Model** | **Fragments** | **RC corr. mean±std** | **Δ vs M0** |
|---|---|---|---|
| M0 | 24 | 0.073±0.022 | 0.000 |
| M1 | 24 | 0.039±0.017 | −0.033 |
| M2 | 24 | 0.070±0.030 | −0.003 |
| M3 | 24 | 0.095±0.015 | +0.023 |
| M4 (+RC SimCSE) | 24 | **0.271±0.067** | **+0.199** |
| M5 | 24 | 0.075±0.009 | +0.003 |
| M6 | 24 | 0.092±0.017 | +0.020 |

Table S37: Paired reverse-complement consistency tests for DNAGPT variants versus M0.

| **Comparison** | $n$ | **Mean Δ** | **Median Δ** | **Pos./neg.** | **Wilcoxon BH** $q$ |
|---|---|---|---|---|---|
| M1 vs M0 | 24 | −0.0334 | −0.0297 | 0/24 | $7.15 \times 10^{-7}$ |
| M2 vs M0 | 24 | −0.0030 | −0.0080 | 8/16 | 0.2643 |
| M3 vs M0 | 24 | +0.0229 | +0.0271 | 21/3 | $3.08 \times 10^{-4}$ |
| M4 vs M0 | 24 | **+0.1988** | **+0.2242** | 23/1 | $\mathbf{1.07 \times 10^{-6}}$ |
| M5 vs M0 | 24 | +0.0029 | +0.0078 | 15/9 | 0.1007 |
| M6 vs M0 | 24 | +0.0196 | +0.0210 | 22/2 | $4.53 \times 10^{-6}$ |

Table S38: Paired reverse-complement consistency tests comparing M4 with other adapted DNAGPT variants.

| **Comparison** | $n$ | **Mean Δ** | **Median Δ** | **Pos./neg.** | **Wilcoxon BH** $q$ |
|---|---|---|---|---|---|
| M4 vs M3 | 24 | +0.1759 | +0.1855 | 24/0 | $1.59 \times 10^{-7}$ |
| M4 vs M5 | 24 | +0.1959 | +0.2123 | 24/0 | $1.59 \times 10^{-7}$ |
| M4 vs M6 | 24 | +0.1793 | +0.1946 | 23/1 | $3.58 \times 10^{-7}$ |
| M4 vs M2 | 24 | +0.2018 | +0.2208 | 24/0 | $1.59 \times 10^{-7}$ |

### S17 HyenaDNA Reverse-Complement Contact-Map Consistency

We repeated the reverse-complement contact-map consistency diagnostic for HyenaDNA variants H0–H6. As in the DNAGPT analysis, each base-pair mutagenesis contact map was recomputed on the reverse-complemented sequence, flipped back to forward coordinates, and compared with the forward map by Pearson correlation. All models were evaluated on the same 24 hg38 windows, with 4094 aligned positions per window.

Table S39: HyenaDNA reverse-complement consistency of base-pair mutagenesis contact maps.

| **Model** | **Fragments** | **RC corr. mean±std** | **Δ vs H0** |
|---|---|---|---|
| H0 | 24 | 0.224±0.051 | 0.000 |
| H1 | 24 | 0.265±0.044 | +0.040 |
| H2 | 24 | 0.360±0.075 | +0.136 |
| H3 | 24 | 0.823±0.026 | +0.599 |
| H4 (+RC SimCSE) | 24 | **0.935±0.018** | **+0.711** |
| H5 | 24 | 0.308±0.021 | +0.083 |
| H6 | 24 | 0.466±0.027 | +0.242 |

Table S40: Paired reverse-complement consistency tests for HyenaDNA variants versus H0.

| **Comparison** | $n$ | **Mean Δ** | **Median Δ** | **Pos./neg.** | **Wilcoxon BH** $q$ |
|---|---|---|---|---|---|
| H1 vs H0 | 24 | +0.0403 | +0.0531 | 18/6 | $2.81 \times 10^{-3}$ |
| H2 vs H0 | 24 | +0.1357 | +0.1555 | 22/2 | $3.62 \times 10^{-5}$ |
| H3 vs H0 | 24 | +0.5986 | +0.6093 | 24/0 | $1.79 \times 10^{-7}$ |
| H4 vs H0 | 24 | **+0.7111** | **+0.7085** | 24/0 | $\mathbf{1.79 \times 10^{-7}}$ |
| H5 vs H0 | 24 | +0.0834 | +0.0775 | 24/0 | $1.79 \times 10^{-7}$ |
| H6 vs H0 | 24 | +0.2415 | +0.2638 | 24/0 | $1.79 \times 10^{-7}$ |

Table S41: Paired reverse-complement consistency tests comparing H4 with other adapted HyenaDNA variants.

| **Comparison** | $n$ | **Mean Δ** | **Median Δ** | **Pos./neg.** | **Wilcoxon BH** $q$ |
|---|---|---|---|---|---|
| H4 vs H2 | 24 | +0.5753 | +0.5775 | 24/0 | $1.19 \times 10^{-7}$ |
| H4 vs H3 | 24 | +0.1125 | +0.1145 | 24/0 | $1.19 \times 10^{-7}$ |
| H4 vs H5 | 24 | +0.6277 | +0.6316 | 24/0 | $1.19 \times 10^{-7}$ |
| H4 vs H6 | 24 | +0.4695 | +0.4667 | 24/0 | $1.19 \times 10^{-7}$ |

The large H4 effect should be interpreted with this diagnostic's proximity to the training objective in mind. Reverse-complement consistency is a proximal strand-symmetry readout for the RC SimCSE objective rather than an independent downstream benchmark, so the H4 increase does not by itself establish a general strand-aware representation gain. Instead, it verifies that the RC contrastive objective strongly imprints the intended symmetry on the base-pair-mutagenesis contact geometry. The cross-architecture magnitude also differs: DNAGPT M4 improves by +0.199 over M0, whereas HyenaDNA H4 improves by +0.711 over H0. HyenaDNA's nucleotide-level representation is measured in a one-base coordinate system, making reverse-complement alignment more direct than in DNAGPT's BPE-tokenised model; the two backbones also have different baseline correlation levels. We therefore interpret each effect relative to its backbone-matched baseline rather than treating the raw deltas as controlled cross-architecture effect sizes, and use this analysis as a mechanism check for the RC objective, not as a stand-alone claim that H4 is universally better.

### S18 DGEB-DNA Out-of-Domain Frozen Embedding Evaluation

We evaluated DNAGPT variants on the nucleotide-sequence subset of DGEB-DNA [West-Roberts et al., 2024], using frozen final-layer embeddings and the official DGEB task evaluators. The selected tasks are out-of-domain relative to the human-genome adaptation corpus and span bacterial, archaeal, and eukaryotic nucleotide sequences. The set includes five evolutionary-distance similarity tasks, where the primary metric is the best correlation between embedding distance and evolutionary distance (`top_corr`), and one E. coli RNA segment clustering task, where the primary metric is V-measure. Values for M2–M6 are reported as mean ± std over three independently adapted checkpoints; M0 and M1 are deterministic checkpoints.

Table S42: DGEB-DNA out-of-domain frozen embedding evaluation. Values report each task's primary metric. `top_corr` is used for the 16S/18S and RpoB phylogeny tasks; V-measure is used for E. coli RNA segment clustering. The final column averages the six primary metrics per seed and then reports mean ± std across seeds.

| Model | Bac 16S | Arch 16S | Euk 18S | RpoB Bac | RpoB Arch | E. coli RNA | Mean |
|---|---|---|---|---|---|---|---|
| M0 (causal) | 0.330±0.000 | 0.100±0.000 | 0.284±0.000 | 0.090±0.000 | 0.088±0.000 | 0.093±0.000 | 0.164±0.000 |
| M1 (+bidir attn) | 0.144±0.000 | 0.108±0.000 | 0.025±0.000 | 0.150±0.000 | -0.037±0.000 | 0.085±0.000 | 0.079±0.000 |
| M2 (+MNTP) | 0.477±0.004 | **0.330**±0.018 | 0.367±0.010 | **0.178**±0.010 | 0.138±0.005 | **0.116**±0.030 | **0.268**±0.011 |
| M3 (+dropout SimCSE) | **0.483**±0.011 | 0.316±0.021 | 0.370±0.014 | 0.158±0.010 | 0.130±0.005 | 0.070±0.031 | 0.254±0.010 |
| M4 (+RC SimCSE) | 0.454±0.010 | 0.260±0.021 | 0.359±0.006 | 0.126±0.002 | **0.185**±0.008 | 0.112±0.022 | 0.249±0.006 |
| M5 (+crop SimCSE) | 0.429±0.006 | 0.310±0.030 | 0.367±0.003 | 0.144±0.005 | 0.109±0.004 | 0.054±0.053 | 0.235±0.005 |
| M6 (+local-shift) | 0.472±0.003 | 0.309±0.035 | **0.376**±0.006 | 0.165±0.002 | 0.131±0.009 | 0.088±0.011 | 0.257±0.007 |

The DGEB-DNA results provide a complementary view of the adaptation stages. The bidirectional patch alone substantially degrades out-of-domain nucleotide embedding geometry (M1 mean 0.079 vs. M0 0.164), whereas MNTP repairs and improves this geometry (M2 mean 0.268). This is consistent with the motif-contact ablation reported in the main text, where M0→M1 also reduces pooled motif signal before MNTP raises it above the causal baseline. Together, the two diagnostics indicate that the bidirectional patch alone disrupts both local perturbation structure and out-of-domain frozen embedding geometry, while MNTP repairs the representation. Contrastive variants remain above the causal baseline but do not uniformly improve over MNTP on these cross-taxa phylogeny and clustering tasks. This supports the task-conditioned interpretation in the main text: MNTP acts as a general representation-repair stage, while crop contrastive adaptation is most beneficial for long-range regulatory pair-relation tasks such as EPI.

We also ran the same DGEB-DNA protocol for the gated-tied HyenaDNA branch (Table S43). Here, the bidirectional patch does not reproduce the DNAGPT degradation: H1 improves the mean score over H0 (0.187 vs. 0.174). The strongest overall variant is the crop contrastive checkpoint H5 (0.223), followed by H4 and H2. Thus, in this out-of-domain diagnostic, gated HyenaDNA adaptation preserves or improves frozen sequence-embedding geometry on average, although the best variant remains task-dependent.

### S19 Caduceus-PS Matched Linear-Probe Baseline

The Caduceus-PS row in the main frozen-baseline table is our matched linear-probe rerun of the public checkpoint `kuleshov-group/caduceus-ps_seqlen-131k_d_model-256_n_layer-16`. We used the `ps` checkpoint, not the `ph` checkpoint, because the reported baseline is labelled

Table S43: DGEB-DNA out-of-domain frozen embedding evaluation for gated-tied HyenaDNA variants. Values report each task's primary metric, with mean ± std over three seeds for H2–H6. H0 and H1 are deterministic under this protocol.

| Model | Bac 16S | Arch 16S | Euk 18S | RpoB Bac | RpoB Arch | E. coli RNA | Mean |
|---|---|---|---|---|---|---|---|
| H0 (causal) | 0.261±0.000 | 0.094±0.000 | 0.297±0.000 | **0.192**±0.000 | 0.043±0.000 | 0.157±0.000 | 0.174±0.000 |
| H1 (gated bidir) | 0.361±0.000 | 0.145±0.000 | 0.305±0.000 | 0.139±0.000 | 0.058±0.000 | 0.112±0.000 | 0.187±0.000 |
| H2 (gated MNTP) | **0.434**±0.007 | 0.088±0.009 | **0.316**±0.012 | 0.129±0.002 | 0.095±0.001 | 0.163±0.060 | 0.204±0.008 |
| H3 (gated dropout) | 0.307±0.042 | 0.091±0.022 | 0.204±0.033 | 0.135±0.015 | **0.164**±0.024 | 0.198±0.026 | 0.183±0.023 |
| H4 (gated RC) | 0.400±0.039 | 0.156±0.058 | 0.275±0.022 | 0.127±0.050 | 0.121±0.025 | 0.191±0.015 | 0.212±0.006 |
| H5 (gated crop) | 0.406±0.070 | **0.189**±0.031 | 0.273±0.018 | 0.137±0.034 | 0.161±0.018 | 0.171±0.031 | **0.223**±0.014 |
| H6 (gated local-shift) | 0.280±0.062 | 0.103±0.038 | 0.203±0.034 | 0.082±0.014 | 0.115±0.018 | **0.204**±0.038 | 0.164±0.022 |

Caduceus-PS. The model was loaded with Hugging Face `AutoModel` using `trust_remote_code=True`, with `AutoModelForMaskedLM` as a loader fallback if needed, and kept frozen throughout evaluation.

To keep the comparison matched to the Step 4 frozen-encoder protocol, sequences were tokenised with the Caduceus tokenizer using truncation at `max_length=1024` and padding to the longest sequence in each batch. The sequence embedding was the mean of the final-layer hidden states over non-padding positions, using the tokenizer attention mask; no CLS, EOS, or task-specific pooling head was used. The pooled embedding was then L2-normalised before fitting the same suite-specific linear probes used for DNAGPT, HyenaDNA, and DNABERT-2 reruns. Thus the Caduceus-PS numbers should be read as a matched frozen mean-pooling baseline under our evaluation code, not as a full exploitation of the 131k-context Caduceus architecture or as task-specific Caduceus fine-tuning.

### S20 Use of Large Language Models

Following the journal's policy on the use of large language models (LLMs), we document here how LLM-based assistants were used in preparing this work, and how we verified their output. We used Claude and ChatGPT in this work.

#### S20.1 Where we used LLM assistance

- **Code.** Drafting parts of the analysis and plotting scripts, for example result aggregation and figure generation. We wrote the core training and evaluation code ourselves.
- **Processing and tabulating results.** Aggregating per-seed experimental output into the summary tables reported in the main text and the Supplementary Material.
- **Language editing.** Editing the wording, grammar and concision of text we had already written.
- **Consistency checking.** Checking the internal consistency of numbers, labels, cross-references and terminology across the manuscript and the Supplementary Material.

#### S20.2 Where we did not use LLM assistance

The research question, the staged adaptation framework, the experimental design, the choice of benchmarks and diagnostics, the interpretation of results and all scientific claims are our own. No experimental result was generated, altered or imputed by an LLM, and no section of the manuscript was drafted from a prompt.

##### S20.3 Verification

We ran all code ourselves and checked its output against the raw experiment logs. We traced every numerical value reported in the manuscript and the Supplementary Material back to the corresponding raw result files and recomputed it. We reviewed all language edits line by line and confirm that they did not change the technical meaning of the text.

### References

Jacob West-Roberts, Joshua Kravitz, Nishant Jha, Andre Cornman, and Yunha Hwang. Diverse genomic embedding benchmark for functional evaluation across the tree of life. *bioRxiv*, 2024. doi: 10.1101/2024.07.10.602933. URL https://doi.org/10.1101/2024.07.10.602933.
